# A glycogenin homolog controls *Toxoplasma gondii* growth via glycosylation of an E3 ubiquitin ligase

**DOI:** 10.1101/764241

**Authors:** Msano Mandalasi, Hyun W. Kim, David Thieker, M. Osman Sheikh, Elisabet Gas-Pascual, Kazi Rahman, Peng Zhao, Nitin G. Daniel, Hanke van der Wel, H. Travis Ichikawa, John N. Glushka, Lance Wells, Robert J. Woods, Zachary A. Wood, Christopher M. West

## Abstract

Skp1, a subunit of E3 Skp1/Cullin-1/F-box protein ubiquitin ligases, is uniquely modified in protists by an O_2_-dependent prolyl hydroxylase that generates the attachment site for a defined pentasaccharide. Previous studies demonstrated the importance of the core glycan for growth of the parasite *Toxoplasma gondii* in fibroblasts, but the significance of the non-reducing terminal sugar was unknown. Here, we find that a homolog of glycogenin, an enzyme that can initiate and prime glycogen synthesis in yeast and animals, is required to catalyze the addition of an α-galactose in 3-linkage to the subterminal glucose to complete pentasaccharide assembly in cells. A strong selectivity of the enzyme (Gat1) for Skp1 in extracts is consistent with other evidence that Skp1 is the sole target of the glycosyltransferase pathway. *gat1*-disruption results in slow growth attesting to the importance of the terminal sugar. Molecular dynamics simulations provide an explanation for this finding and confirm the potential of the full glycan to control Skp1 organization as in the amoebozoan *Dictyostelium* despite the different terminal disaccharide assembled by different glycosyltransferases. Though Gat1 also exhibits low α-glucosyltransferase activity like glycogenin, autoglycosylation is not detected and *gat1*-disruption reveals no effect on starch accumulation. A crystal structure of the ortholog from the crop pathogen *Pythium ultimum* explains the distinct substrate preference and regiospecificity relative to glycogenin. A phylogenetic analysis suggests that Gat1 is related to the evolutionary progenitor of glycogenin, and acquired a role in glycogen formation following the ancestral disappearance of the underlying Skp1 glycosyltransferase prior to amoebozoan emergence.

## INTRODUCTION

A novel glycosylation pathway whose glycosyltransferases (GTs) reside in the cytoplasm evolved early in eukaryotic evolution and persisted in most branches of protist radiation before disappearing in land plants, yeasts, and animals. Studies of the pathway in the amoebozoan *Dictyostelium discoideum* and the apicomplexan *Toxoplasma gondii* show a role in O_2_-sensing, owing to the requirement for an O_2_-dependent prolyl 4-hydroxylase, PhyA, that modifies a critical proline residue of the highly conserved nucleocytoplasmic protein Skp1 (West and Blader, 2015). PhyA action generates a 4(*trans*)-hydroxyproline (Hyp) residue, which serves as the attachment point of the pentasaccharide that is assembled by the sequential action of five GT activities. Thus, Skp1 is a unusual example of a glycoprotein that is not only glycosylated in a complex manner in the cytoplasm, but also remains to function in this compartment and the nucleus, rather than being exported (West and Hart, 2017). Skp1 is the only protein that is detectably glycosylated by any of the five GT activities in *D. discoideum* (West and Blader, 2015), which is at a variance from most GTs, which typically modify a spectrum of proteins. Genetic manipulation of Skp1 expression, substitution of the target Pro, and genetic interactions with a Skp1 GT, affect *D. discoideum* O_2_-sensing in ways that are consistent with Skp1 being the functional target of prolyl hydroxylation in O_2_-sensing (Wang et al., 2011). Skp1 is a subunit of the SCF class of E3 ubiquitin (Ub) ligases, and hydroxylation/glycosylation promotes its interaction with three different F-box proteins (FBPs) *in vitro* and *in vivo* (Sheikh et al., 2014, 2015), implicating an effect on their respective E3(SCF)Ub ligases. The mechanism by which the Hyp-linked pentasaccharide controls the activity of Skp1 is unclear, but structural studies suggest an effect on Skp1 conformation that is consistent with receptivity to binding FBPs (Sheikh et al., 2017; Xu et al., 2018; West and Kim, 2019).

Studies of the GTs that catalyze assembly of the glycan in *T. gondii* revealed that, while a linear pentasaccharide is assembled at a conserved prolyl residue as in *D. discoideum*, the glycan has a distinct structure (Fig. 1A). The core trisaccharides are evidently identical, and the enzymes that assemble them are orthologous, but the GT that assembles the fourth and fifth sugars in *D. discoideum* is not detected in the *T. gondii* genome. Further studies revealed that the fourth sugar is an α3-linked Glc, rather than α3-linked Gal, whose linkage is catalyzed by novel enzyme, Glt1 (Rahman et al., 2017). The question of the identity of the fifth sugar, known to be a hexose from MS studies, and its mechanism of addition, remains unanswered. Genomics studies predict the existence of four cytoplasmically localized GTs whose functions are not assigned (Gas-Pascual et al., 2019), and one of these, referred to as Gat1, was predicted to be the missing Skp1 GT on account of the apparent co-distribution of its gene in protists that possess Glt1-like genes (Rahman et al., 2017). However, this gene is highly similar to glycogenin, a dimeric α4-glucosyl transferase that can prime the synthesis of glycogen in the cytoplasm of yeast and animals by a mechanism that involves auto-glucosylation. The cyst-forming stage of *T. gondii* accumulates crystalline amylopectin (Coppin et al., 2003), an α1,4Glc polymer with α1,6-linked branches that resembles glycogen, in its cytoplasmic compartment. *T. gondii* amylopectin is assembled by a UDP-Glc based metabolism that is related to the floridean starch of the red alga *Cyanidioschyzon merolae* and, to a lesser extent, to that of glycogen storing animals and fungi. Homologs of glycogen synthase (Coppin et al., 2003) and glycogen phosphorylase (Sugi et al., 2017) regulate the accumulation of amylopectin in this parasite, and TgGat1 has been annotated as a glycogenin (Coppin et al., 2005; Sugi et al., 2017). Related genes have been implicated in promoting starch formation in red algae (Pancha et al., 2019a, 2019b).

**Fig. 1.**
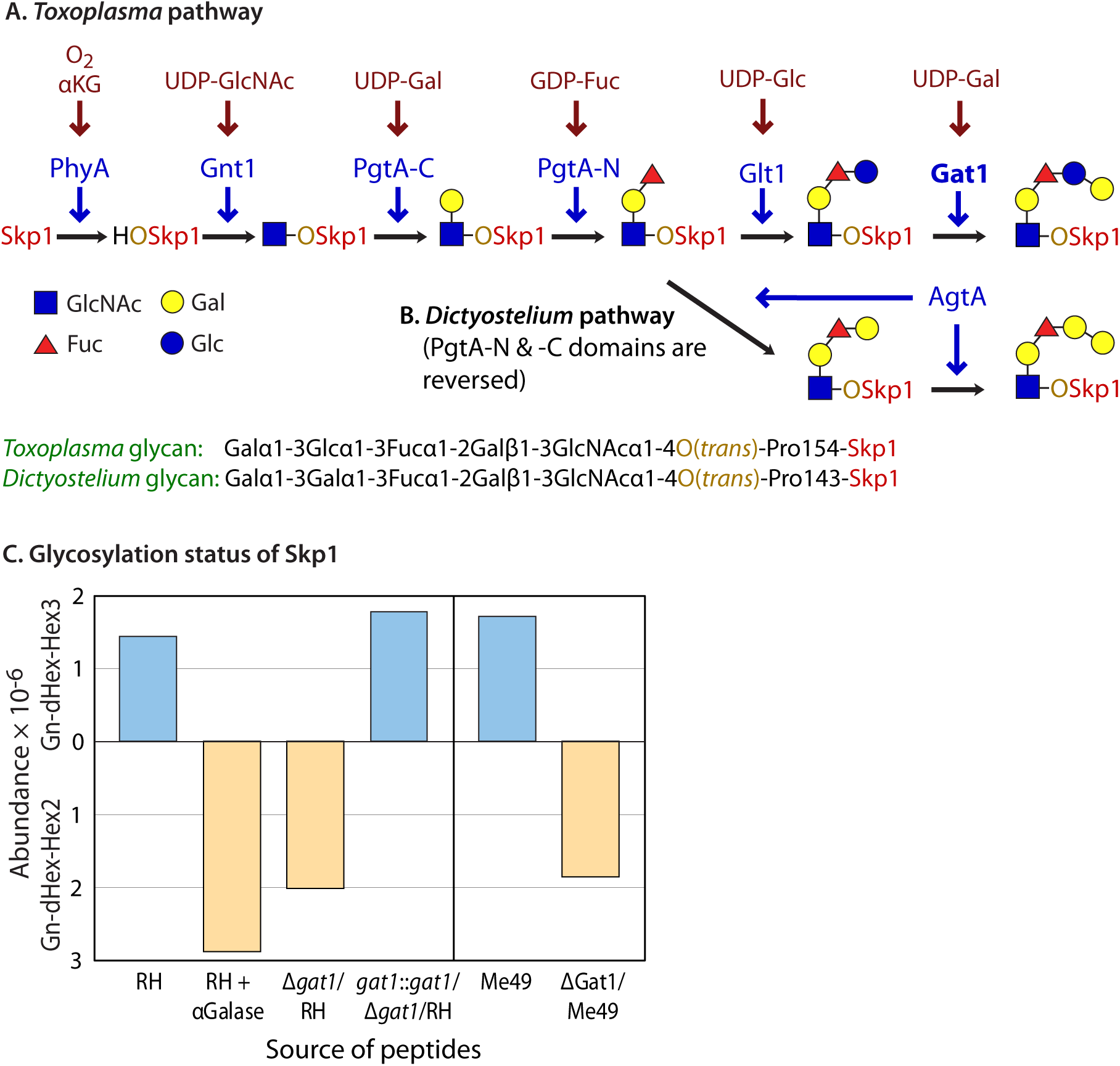
Gat1 is required for terminal α-galactosylation of Skp1. (**A**, **B**) Schematic of Skp1 glycosylation pathway in *Toxoplasma* (Rahman et al., 2017, and herein) and *Dictyostelium* (West and Blader 2015). Gat1, the final enzyme in the *Toxoplasma* pathway, is in bold. (**C**) Glycopeptide analysis of Skp1 immunoprecipitated from type 1 RH and type 2 ME49 strains and their genetic derivatives, after trypsinization and mass analysis by nLC-MS. Pentasaccharide levels are shown above the x-axis, and tetrasaccharide levels are below, after normalization to total Skp1 peptides detected. In addition, tryptic peptides from RH were treated with green coffee bean α-galactosidase prior to nLC-MS analysis. Similar results were obtained in an independent trial from RH and Δ*gat1*/RH (not shown). See Table S2 and Fig. S5 for primary data.

The identity of the missing Skp1 GT is critical for determining the terminal sugar and its linkage in *T. gondii*, and to assess the target specificity of the GT and the role of the glycan in cellular regulation. Furthermore, defining the precise glycan structure would enable a test of the *cis*-acting structure model for how it controls Skp1 activity in *D. discoideum* (West and Kim, 2019). Our new studies show that Gat1 is a conserved, cytoplasmic, retaining α3-galactosyl transferase, not an α4-glucosyl transferase as for glycogenin, that modifies the Glc terminus of Glcα1,3Fucα1,2Galβ1,3GlcNAcα1-Skp1, to support optimal growth on fibroblast monolayers. Despite the glycan differences between *T. gondii* and *D. discoideum*, we find that both have similar potential to modulate Skp1 structure. Finally, we demonstrate the existence of a similar process in the oomycete plant pathogen *Pythium ultimum* (Pu), and rationalize the difference in Gat1 and glycogenin activities based on the crystal structure of PuGat1. *P. ultimum* is the agent for root rot disease in agriculturally important crops (Lévesque et al., 2010; Kamoun et al., 2015), and is related to *P. insidiosum*, the agent for debilitating pythiosus in humans and other mammals (Gaastra et al., 2010). We suggest that the glycogenin lineage originally evolved as a modifier of Skp1 that contributed to a novel mechanism of O_2_-sensing in a range of aerobic unicellular and pathogenic eukaryotes, before transitioning into glycogenin during Opisthokont evolution.

## RESULTS

### The fifth Skp1 sugar is an α-linked Gal and depends on TgGat1

Previous studies described the mechanism of assembly of the first four sugars on TgSkp1-Pro154 (Fig. 1A), but the left the identity of the final sugar (other than its being a hexose) unresolved (Rahman et al., 2016, 2017). The corresponding sugar in *Dictyostelium* (Fig. 1B) is a 3-linked αGal that is susceptible to removal with green coffee bean α-galactosidase. Similar treatment of tryptic peptides from TgSkp1, isolated from the standard type 1 strain RH by immunoprecipitation, resulted in complete conversion of the pentasaccharide form of the glycopeptide to the tetrasaccharide form, indicating that the terminal Hex is an αGal residue (Fig. 1C).

According to a recent genomic analysis (Rahman et al., 2017), TgGat1 is a candidate for the unknown glycosyltransferase that catalyzes the addition of the fifth and final sugar (Fig. 1A). To test the dependence of Skp1 glycosylation on Tg*gat1*, the gene was disrupted using CRISPR/Cas9 in the RH strain, yielding *gat1*Δ as described in Fig. S4A. PCR studies confirmed replacement with a *dhfr* cassette, and enzyme assays (see below) showed a loss of enzyme activity. The resulting clone produced a version of Skp1 in which the 5-sugar glycopeptide was no longer detectable, but ions corresponding to the 4-sugar glycopeptide were detected at similar abundance (Fig. 1C). Similar results were obtained in the type 2 ME49 strain (Fig. S4A, Fig. 1C), and in strain RHΔΔ in which *gat1* was disrupted by homologous recombination using a different selection marker, referred to as Δ*gat1-1* (Table 1; Fig. S3B; not shown). The similar results obtained by different genetic methods in distinct genetic backgrounds indicate that Tg*gat1* is required for addition of the terminal sugar, but whether this was a direct or indirect effect was unclear.

**Table 1.**
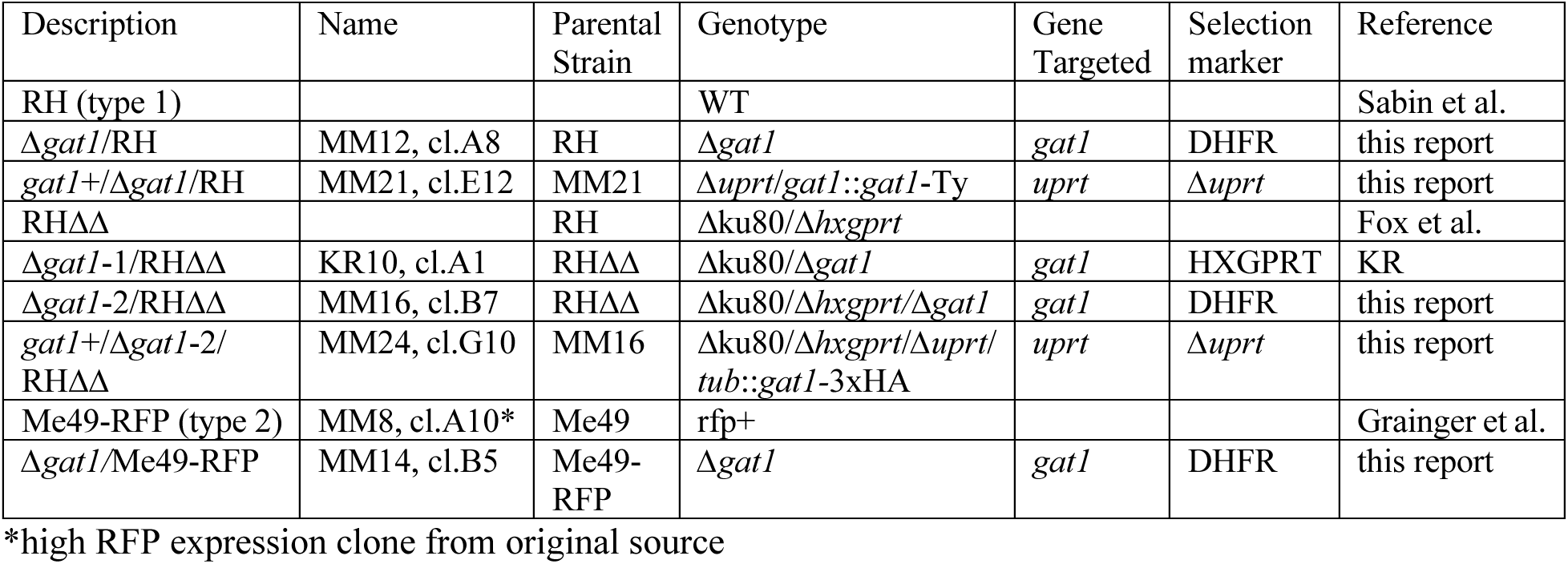
Strains employed in this study

| Description | Name | Parental Strain | Genotype | Gene Targeted | Selection marker | Reference |
| --- | --- | --- | --- | --- | --- | --- |
| RH (type 1) |  |  | WT |  |  | Sabin et al. |
| $\Delta gat1$/RH | MM12, cl.A8 | RH | $\Delta gat1$ | *gat1* | DHFR | this report |
| *gat1*<sup>+</sup>/$\Delta gat1$/RH | MM21, cl.E12 | MM21 | $\Delta uprt/gat1::gat1$-Ty | *uprt* | $\Delta uprt$ | this report |
| RH$\Delta\Delta$ | | RH | $\Delta ku80/\Delta hxgprt$ | | | Fox et al. |
| $\Delta gat1$-1/RH$\Delta\Delta$ | KR10, cl.A1 | RH$\Delta\Delta$ | $\Delta ku80/\Delta gat1$ | *gat1* | HXGPRT | KR |
| $\Delta gat1$-2/RH$\Delta\Delta$ | MM16, cl.B7 | RH$\Delta\Delta$ | $\Delta ku80/\Delta hxgprt/\Delta gat1$ | *gat1* | DHFR | this report |
| *gat1*<sup>+</sup>/$\Delta gat1$-2/RH$\Delta\Delta$ | MM24, cl.G10 | MM16 | $\Delta ku80/\Delta hxgprt/\Delta uprt/tub::gat1$-3xHA | *uprt* | $\Delta uprt$ | this report |
| Me49-RFP (type 2) | MM8, cl.A10* | Me49 | rfp<sup>+</sup> |  |  | Grainger et al. |
| $\Delta gat1$/Me49-RFP | MM14, cl.B5 | Me49-RFP | $\Delta gat1$ | *gat1* | DHFR | this report |

\*high RFP expression clone from original source

### Dependence of parasite growth on *gat1*

Parasites require contact with and invasion of mammalian host cells to establish a niche within an intracellular parasitophorous vacuole in order to proliferate. In the context of two-dimensional monolayers of fibroblasts, parasites divide and eventually lyse out to invade neighboring cells to repeat the cycle. The area of the resulting plaque provides a measure of efficiency of a number of cellular processes. Past studies showed that plaque size growth is compromised by mutational blockade of Skp1 hydroxylation and earlier steps of the glycosylation pathway (Xu et al., 2012; Rahman et al., 2016, 2017). Here we find that disruption of *gat1* resulted in modestly smaller plaques, on average, after 5 d of growth, in either the RH or RHΔΔ background (Fig. 2A,B). Furthermore, plaque sizes were similarly reduced in an independent *gat1*Δ strain prepared in RHΔΔ using homologous recombination without CRISPR-Cas9 (Fig. 2C). To determine whether the effects were specific to the genetic lesion at the *gat1* locus, the RH and RHΔΔ KO strains generated using CRISPR/Cas9 were modified again by CRISPR/Cas9 to introduce single copies of epitope tagged versions of the *gat1* coding locus, downstream of an endogenous *gat1* promoter cassette or a tubulin promoter cassette, respectively, into the *uprt* locus. The expected insertions were confirmed using PCR (Figs. S3C, S4B). As a result, TgGat1-3xHA could be detected by Western blotting of tachyzoites in the RHΔΔ background (Fig. S4D) and, as discussed below, the complemented RHΔΔ strain restored Skp1 glycosylation according to a biochemical complementation test. Although TgGat1-Ty expressed under its own promoter cassette in the RH background was not detected (not shown), enzyme activity was partially restored in the TgGat1-Ty strain (Fig. S4C). Both strains exhibited larger plaque sizes than their respective KO parents (Fig. 2A, B).

**Fig. 2.**
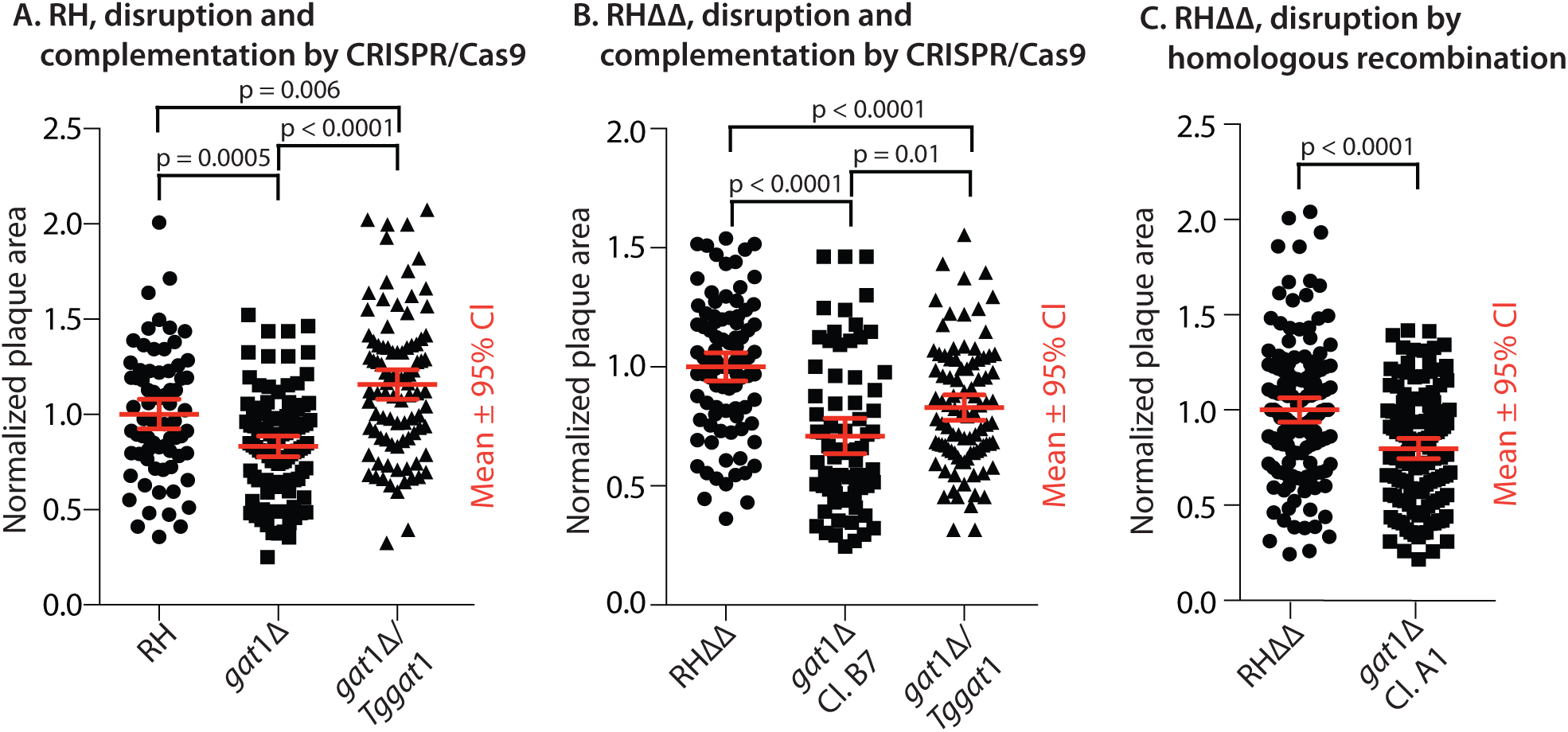
Dependence of parasite growth on Gat1. Parasites were plated at clonal density on two-dimensional monolayers of human foreskin fibroblasts (HFFs), and allowed to invade, proliferate, lyse out, and reinfect neighboring fibroblasts. After 5.5 d, cultures were fixed, stained, and analyzed for the areas occupied by lysed fibroblasts (plaques). Data from each of 3 independent trials, which were each normalized to the parental strain, were merged for presentation. (**A**) Comparison of the type 1 RH strain before and after *gat1* replacement using CRISPR/Cas9, and complementation with *gat1* under control of its own promoter cassette in the uprt locus. (**B**) Comparison of RHΔΔ, *gat1-2*Δ/RHΔΔ, and the latter complemented with gat1 under control of a tubulin promoter in the uprt locus. (**C**) Comparison of *gat1-1*Δ/RHΔΔ, prepared by homologous recombination.

### Evolution of the Gat1 sequence

An evolutionary analysis was conducted to gain insight into the function of the TgGat1 gene product. Based on searches of genomic databases using BLASTP, TgGat1 is most closely related to CAZy GT8 sequences. The top-scoring hits, with Expect values of <10^-32^, were found only in protists that contain *Toxoplasma* PgtA-like and Glt1-like sequences (Fig. S7) and lack *Dictyostelium* AgtA-like sequences, suggesting a common function. The most similar sequences, in searches seeded with the putative catalytic domain, belong to glycogenin, with Expect values of ≥E^-27^. All other homologous sequences had Expect values of ≥10^-22^. Glycogenin is a self-modifying α4-glucosyltransferase that is a primer for glycogen synthesis by glycogen synthase and branching enzyme. *T. gondii* produces amylopectin, whose sequence is similar to that of glycogen, raising the possibility that Gat1 has a similar function in *T. gondii*.

Glycogenin consists of a CAZy GT8 family catalytic domain plus a C-terminal glycogen synthase binding domain separated by a linker, whereas Gat1 consists only of a single catalytic domain (Fig. 3A). TgGat1 is predicted to be a 345-amino acid protein encoded by a single exon gene in the Type I GT1 strain (TGGT1_310400). It is 34% identical to rabbit (*Oryctolagus cuniculus*) glycogenin over the catalytic domain, but includes a poorly conserved 90-amino acid sequence that interrupts the catalytic domain (Fig. 3A). This region is likely unstructured based on secondary structure prediction by the XtalPred server (Slabinski et al., 2007). A homologous Gat1-like sequence found in another, distantly related, protist, *Pythium ultimum*, (Uniprot K3WC47) was analyzed because it lacks this sequence. PuGat1 is predicted to be a 266-amino acid protein encoded by a 2-exon gene, annotated as PYU1_G002535-201 (Transcript ID PYU1_T002538).

**Fig. 3.**
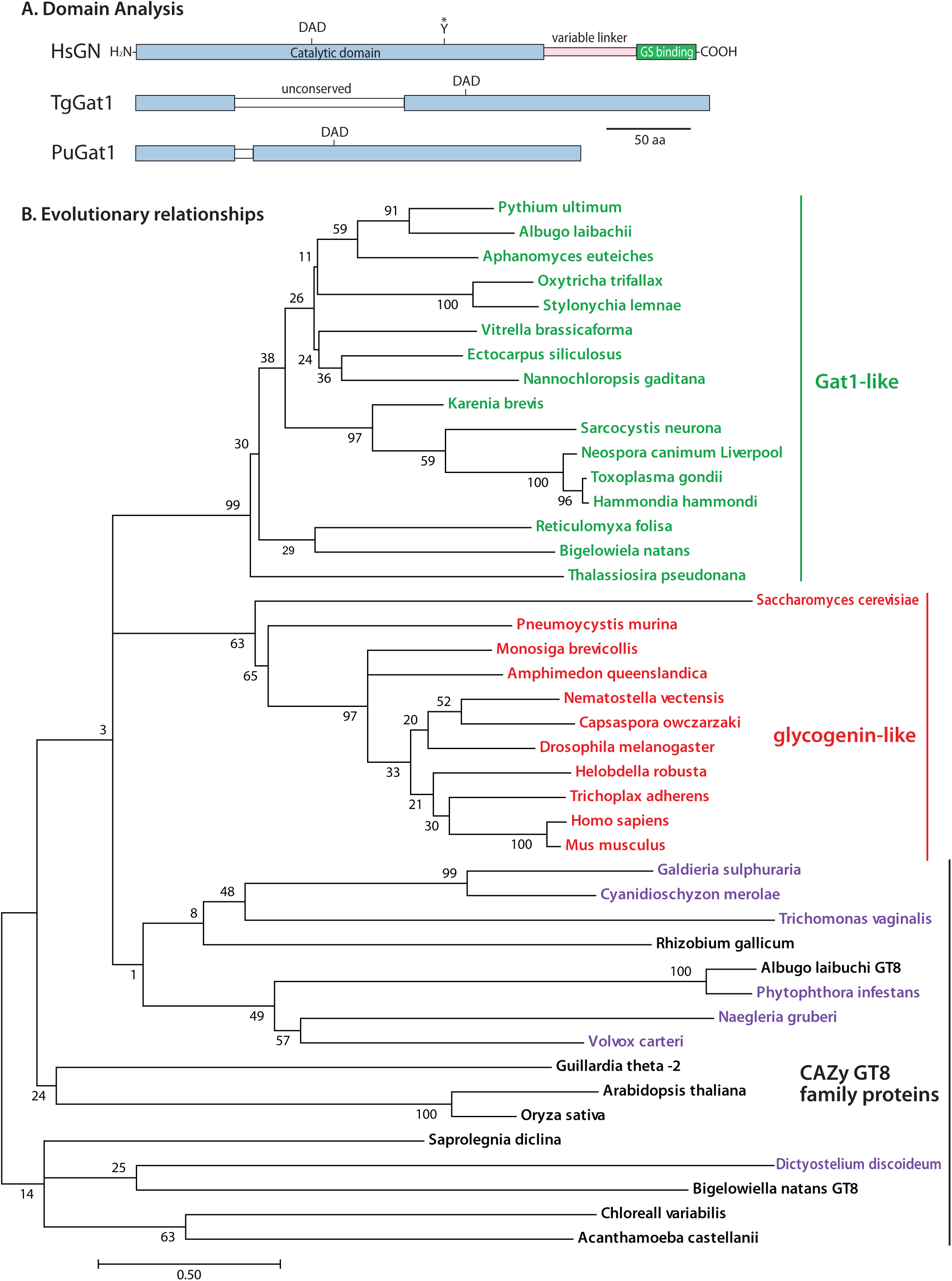
Phylogenetic analysis of Gat1- and glycogenin-like sequences. (**A**) Domain analysis of Gat1 from *T. gondii* and *P. ultimum* in comparison with human glycogenin-1. (**B**) The evolutionary history of the sequence of the Gat1 catalytic domain was inferred by using a Maximum Likelihood method. The tree with the highest log likelihood (−13279.56) is shown. Gat1 and Gat1-like sequences are colored green, glycogenin and glycogenin-like sequences are in red, and characterized and other selected other CAZy GT8 sequences are in black, or purple if predicted to reside in the secretory pathway rather than the cytoplasm. The percentage of trees in which the associated taxa clustered together is shown at each branch. Branch lengths are measured by the number of substitutions per site. See Figs. S6-S8 for alignments.

To evaluate the evolutionary relationship of these putative glycosyltransferases, their catalytic domains those of the most closely related or known sequences from the CAZy GT8 family (Fig. S7) were aligned (Fig. S8) and analyzed by a Maximum Likelihood method (Fig. 3B). The results suggest that Gat1 and glycogenin evolved separately from a common ancestor. Though the last common ancestor was not resolved, Gat1 was presumably the predecessor to glycogenin owing to its presence in more primitive unicellular eukaryotes which bear no evidence of glycogenin-like sequences, and putative orthologs of Gat1 and glycogenin have not been observed in the same clade. Gat1 and glycogenin each possess unique conserved sequence motifs (Fig. S6) that likely support functional differences.

### Enzymatic characteristics of Gat1

To address whether TgGat1 has the potential to directly modify Skp1, the predicted full-length protein was expressed as a His_6_-tagged conjugate in *E. coli*, purified on a TALON resin, and treated with TEV protease leaving an N-terminal GlyHis-dipeptide stub before the start Met (Fig. 4A). The presumptive ortholog from *Pythium ultimum* was prepared similarly. A screen for UDP-sugar hydrolysis activity of TgGat1 yielded, after extended reaction times, only UDP-Gal and UDP-Glc as candidate substrates from a panel of six common UDP-sugars (Fig. S9A). A quantitative comparison showed approximately 7-fold greater activity toward UDP-Gal than UDP-Glc (Fig. 4B).

**Fig. 4.**
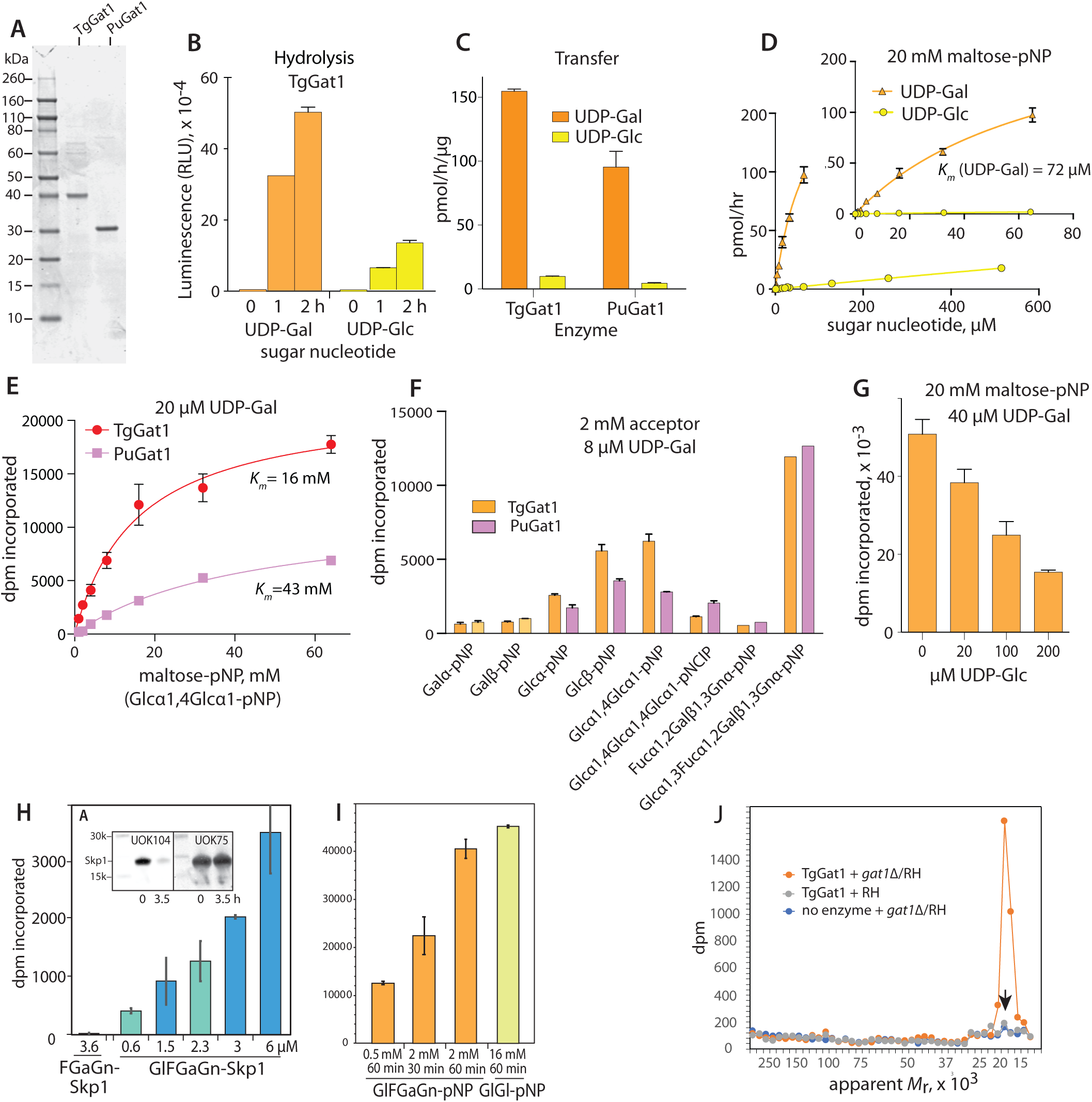
Substrate specificity of Gat1. (**A**) Recombinantly expressed and purified preparations of TgGat1 and PuGat1 were analyzed by SDS-PAGE and staining with Coomassie blue. (**B**) Temporal dependence of UDP-Gal and UDP-Glc hydrolysis. The averages and standard deviations of 3 technical replicates are shown. A similar profile was observed with a different enzyme concentration. See Fig. S9E for a trial with higher enzyme concentrations. (**C**) Transferase activity utilizing 8 µM UDP-Gal or UDP-Glc toward 20 mM maltose-pNP for TgGat1 and PuGat1. The averages and standard deviations of two technical replicates are shown; similar profiles were in 2 independent assays with a different TgGat1 preparation. (**D**) UDP-Gal and UDP-Glc concentration dependence of TgGat1 transferase activity toward 20 mM maltose-pNP. The averages and standard deviations of two technical replicates are shown, and an independent trial with TgGat1 and PuGat1 against UDP-Gal is shown in Fig S9F. (**E**) Maltose-pNP concentration dependence of TgGat1 and PuGat1 transferase activity from 20 µM UDP-Gal. The averages and standard deviations of two technical replicates are shown. (**F**) Relative Gal-transferase activity of TgGat1 and PuGat1 toward different acceptors. The averages and standard deviations of three technical replicates are shown. Similar results were obtained in three independent trials. (**G**) Effect of UDP-Glc concentration on the Gal-transferase activity of TgGat1. Reactions were incubated for 1 h. The averages and standard deviations of two technical replicates are shown. **(H**) Gal-transferase activity of TgGaT1 toward varied concentrations of GlFGaGn-Skp1, in the presence of 40 µM UDP-Gal (1 µCi) after 1 h incubation. Data from independent preparations of TgSkp1 are colored in different shades. FGaGn-Skp1 is included for comparison. Error bars represent S.D. of duplicate measurements. Inset shows Western blots of the Skp1 preparations used, where FGaGn-Skp1, which is recognized specifically by pAb UOK104, is largely converted in a 3.5-h reaction using Glt1 and UDP-Glc to GlFGaGn-Skp1, which is recognized only by the pan-specific pAb UOK75. (**I**) Reactions with synthetic oligosaccharides conjugated to pNP were conducted in parallel using the same conditions. (**J**) Biochemical complementation to detect Gat1 substrates. Desalted S100 extracts of RH and *gat1*Δ/RH were reacted with recombinant Gat1 in the presence of UDP-[^3^H]Gal, and the product of the reaction was separated on an SDS-PAGE gel which was sliced into 40 bands for liquid scintillation counting. The migration position of Skp1 is marked with an arrow. See Figs. S9H and S9I for trials using different strains.

The ability of Gat1 to transfer Gal or Glc to another sugar, rather than water, was tested using a substrate for glycogenin, Glcα1,4Glcα1-pNP (maltose-pNP), which mimics the terminal disaccharide of glycogen and starch and has a terminal αGlc as found on the Skp1 tetrasaccharide. Although Gat1 from either *T. gondii* or *P. ultimum* could modify maltose-pNP using either sugar nucleotide (Fig. 4D), the enzymes strongly preferred UDP-Gal. TgGat1 exhibited a *K_m_* value for UDP-Gal of 30-72 µM but was not saturated with UDP-Glc at 0.5 mM. PuGat1 had a slightly lower apparent *K_m_* for UDP-Gal (Fig. S9F), and both values were greater than the 2-4 µM values reported for rabbit and yeast glycogenins for UDP-Glc (Lomako et al., 1988, de Paula et al., 2005). TgGat1 and PuGat1 exhibited higher *K_m_* values for maltose-pNP in the range of 16-43 mM (Fig. 4D), which were greater than the 4 mM value reported for rabbit glycogenin (Cao et al., 1995).

Extending the acceptor to 3 sugars or decreasing it to one resulted in less activity, but either anomer of Glc-pNP was acceptable (Fig. 4F). A similar pattern was observed for the *Pythium* and *Toxoplasma* enzymes. The enzymes were specific for terminal Glc acceptors as activity towards Gal-pNP was not detected. In comparison, GlFGaGn-pNP, which mimics the natural acceptor on Skp1, was a superior acceptor substrate (Fig. 4F) with a *K_m_* of 1.5 mM (Fig. S9G), and the truncated trisaccharide FGaGn-pNP was inactive indicating that Glc was the position of attachment. Thus the Gat1 enzymes preferred their native tetrasaccharide acceptor substrate and UDP-Gal as a donor, but tolerated, with low efficiency, the preferred substrates of glycogenin, UDP-Glc and α4-linked oligomers Glc.

The importance of acceptor glycan context was examined using GlFGaGn-Skp1, which was prepared by reaction of FGaGn-Skp1 with UDP-Glc and Glt1 resulting in loss of the trisaccharide epitope (Fig. 4H inset). In a direct comparison of acceptor concentration dependence, TgGat1 was about 33x more active toward GlFGaGn-Skp1 than GlFGaGn-pNP (Fig. 4H,I), based on ∼33x less activity (dpm incorporated) at 0.001x the substrate concentration (substrate concentrations were both in the linear response range). The reaction with GlFGaGn-Skp1 did not approach saturation at the highest concentration tested, 6 µM. Thus the TgGat1 reaction was much more efficient when the tetrasaccharide was associated with its native substrate Skp1, and thus consistent with Gat1 being directly responsible for modifying Skp1 in the cell.

A characteristic of glycogenin is its ability to modify the HO-group of a Tyr side chain near its active site with αGlc, and then to repeatedly modify the 4-position of the Glc with another αGlc, and repeat the process up to 8-12 sugars. When isolated in their recombinant forms from *E. coli*, neither TgGat1 nor PuGat1 were found to be glycosylated, based on an exact mass measurements using nLC/MS (Fig. S10A-E). Furthermore, following incubation with either UDP-Gal or UDP-Glc, no change in SDS-PAGE mobility (Fig. S10D) or exact mass were observed (Fig. S10E). Thus, no evidence for autoglycosylation activity of Gat1 from either species could be detected.

### Skp1 is the only detectable protein substrate in parasite extracts

GlFGaGn-Skp1 is a substrate for Gat1, but are there others? This was addressed by complementing extracts of *gat1Δ* parasites with recombinant TgGat1 in the presence of UDP-[^3^H]Gal, and measuring incorporation of ^3^H after display of the proteome on a 1D SDS-PAGE gel. A high level of incorporation of ^3^H that depended on the addition of enzyme was observed at the position of Skp1 (Fig. 4J), as expected, but negligible dpm were detected elsewhere in the gel. Furthermore, negligible dpm were incorporated into Skp1 in RH parental cells, indicating that little GlFGaGn-Skp1 accumulates in wild-type cells. Similar results were observed in studies of *gat1*Δ clones in the RHΔΔ and Me49 backgrounds (Fig. S9H, I). Finally, complementation of the *gat1*Δ clone in RHΔΔ with Gat1 expressed under the tubulin promoter resulted in absence of measurable incorporation into Skp1, confirming specificity for Gat1 expression *per se*.

### Starch accumulates in Tg*gat1*Δ parasites

Although Gat1 was unable to serve as its own GT acceptor in the manner of glycogenin, its ability to modify α4Glc oligomers *in vitro*, albeit with low efficiency, raised the possibility that it may affect amylopectin (starch) biosynthesis in cells by applying αGal residues to its non-reducing termini. Starch normally accumulates to substantial levels in bradyzoites, a slow growing form of the parasite that is induced by stress. Since induction of bradyzoite differentiation in cell culture is more efficient in type 2 strains, Me49 and *gat1*Δ/Me49 cells were induced by pH up-shift and examined for differentiation by labeling of bradyzoite cyst walls with FITC-DBA-lectin, and for starch with the Periodic acid/Schiff’s base reagent. As shown in Fig. 5, ME49 bradyzoites accumulated substantial levels of starch relative to tachyzoites, and no difference in the pattern or level was ascertained in *gat1*Δ cells using this qualitative assessment. Thus Gat1 is not required for starch synthesis, which is consistent with the absence of terminal Gal residues in sugar composition analyses in *T. gondii* starch (Coppin et al., 2005; Guerardel et al., 2005), and does not dramatically influence starch abundance.

**Fig. 5.**
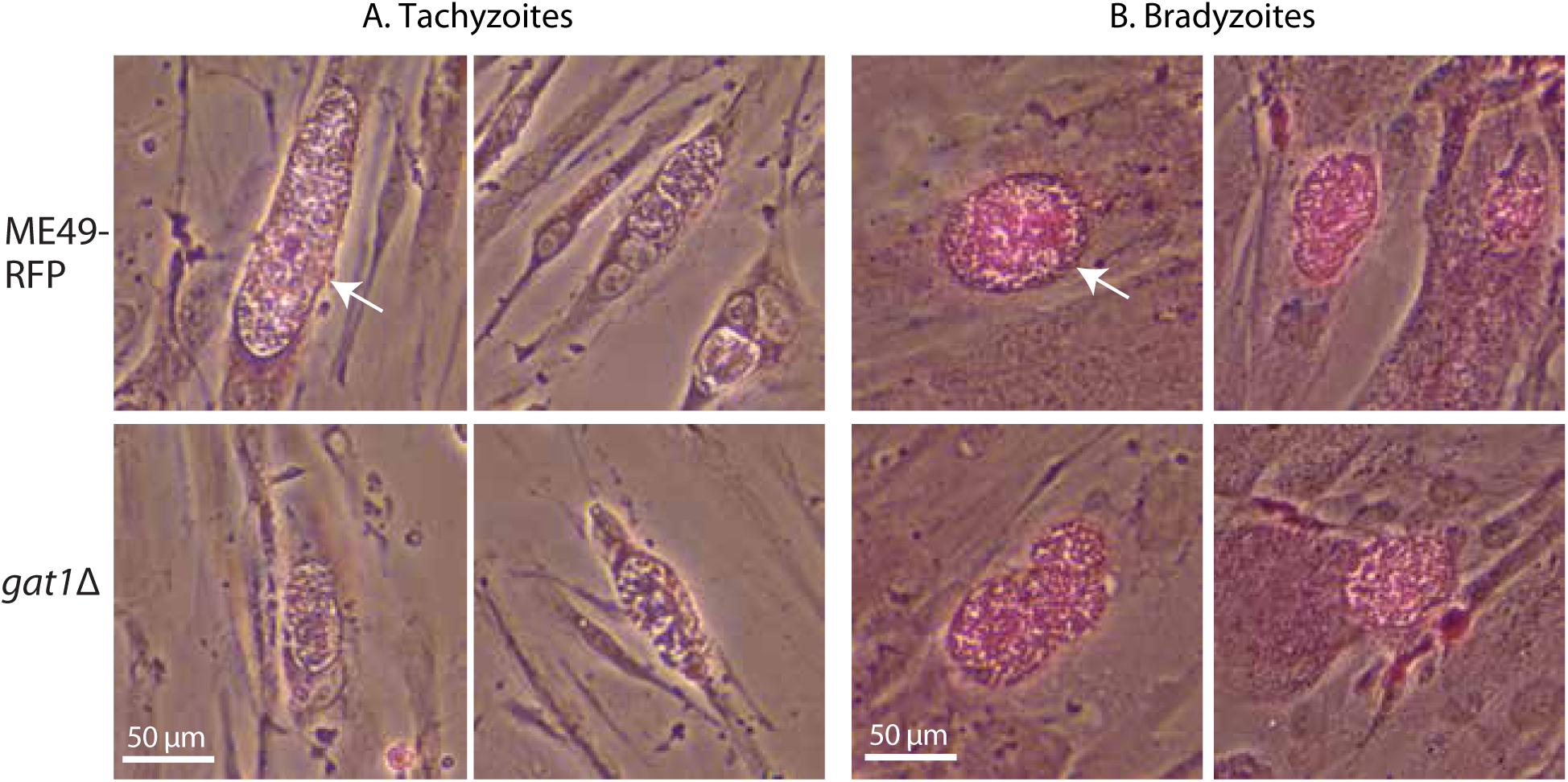
Starch accumulates normally in *gat1*Δ parasites. To promote normal starch accumulation, rapidly proliferating tachyzoites (panel **A**) of the type II strain Me49 (RFP expressing) and its *gat1*Δ derivative were induced to differentiate as slow-proliferating bradyzoite cysts (panel **B**) in human foreskin fibroblasts. Cultures were fixed and stained with Periodic acid/Schiff’s base to reveal starch as a purple adduct. Arrow indicates a parasitophorous vacuole containing dozens to hundreds of tachyzoites within a fibroblast. Arrowhead indicates a cyst containing dozens of slow-growing bradyzoites, as confirmed by labeling of the cyst wall with FITC-DBA lectin (not shown). Scale bar = 50 µm. Two independent trials yielded similar results.

### Gat1 generates a Galα1,3Glc-linkage

To determine the glycosidic linkage of the αGal residue transferred by TgGat1, the previously prepared (^13^C_6_)GlFGaGn-pNP (Rahman et al., 2017) was modified with TgGat1 using UDP-[U-^13^C]Gal as the donor substrate. The pentasaccharide reaction product (approximately 25% conversion) was analyzed together with the tetrasaccharide starting material by NMR. The previous assignment of the chemical shifts of the tetrasaccharide (Rahman et al., 2017) facilitated provisional assignment of the additional terminal Gal chemical shifts using the *CASPER* program (Jansson et al., 2006) and confirmed by analysis of the 2D COSY, TOCSY and HMBC spectra (Fig. S11). One-dimensional ^1^H-NMR spectra reveals the presence of the mixture of tetra- and pentasaccharides, followed by a downfield shift in the ^13^C-Glc-H1 peaks upon linkage to the terminal ^13^C-Gal (Fig. S11A). The HMBC ^1^H –^13^C correlation spectrum shows the (Fig. S11B, center panel) connection from Gal-H1 to Glc-H3, consistent with the downfield peak in the ^1^H –^13^C-HSQC spectrum (Fig. S11B, top panel), and the proton resonances in the HSQC-TOCSY (Fig. S11B, bottom panel), establishing the glycosidic linkage between the terminal αGal and underlying αGlc as 1à3. Finally, a ^1^H –^1^H-COSY spectrum confirms the assignments of the underlying Glc-H1, H2 and H3 (Fig. S11C). Similar results were obtained when the tetrasaccharide was modified in the presence of PuGat1 (data not shown). Taken together, our NMR analyses are most consistent with the glycan structure: Galα1,3Glcα1,3Fucα1,2Galβ1,3GlcNAcα-, indicating that TgGat1 is a retaining UDP-Gal:glucoside α1,3-galactosyltransferase. Not only does Gat1 transfer the α-anomer of a different sugar compared to glycogenin, it attaches it to a different position (4-not 3-) of the acceptor αGlc residue.

### Skp1 glycan conformation

Previous NMR and molecular dynamics (MD) analyses of DdSkp1 suggest the glycan forms a relatively stable interaction with the polypeptide that correlates with opening the F-box combining region by extension of the terminal helix (helix-8) (Sheikh et al., 2017). To assess whether the glycan of TgSkp1 is capable of this functionality despite the difference of the fourth sugar, computational methods applied to analyze glycosylated DdSkp1 were repeated on glycosylated TgSkp1. Energy-minimized structures of the glycans from DdSkp1 and TgSkp1 revealed only a differences in the position of O4 of the fourth sugar (due to the Glc/Gal configurational inversion) (Fig. S12A). As before, six all-atom MD simulations were performed without coordinate constraints for 250 ns each. Three of the simulations began with a 50-ns pre-equilibration of the glycan with Cα atom constraints on the polypeptide while the others proceeded directly. Running six independent simulations for 250 ns rather than one simulation for 1.5 μs allowed greater sampling of conformational space. As before, the simulations did not converge on a common structure, so a combined analysis of the six simulations was conducted to identify trends of glycan-protein interactions that occurred during helix extension (Fig. 6A) and were absent when retracted.

**Fig. 6.**
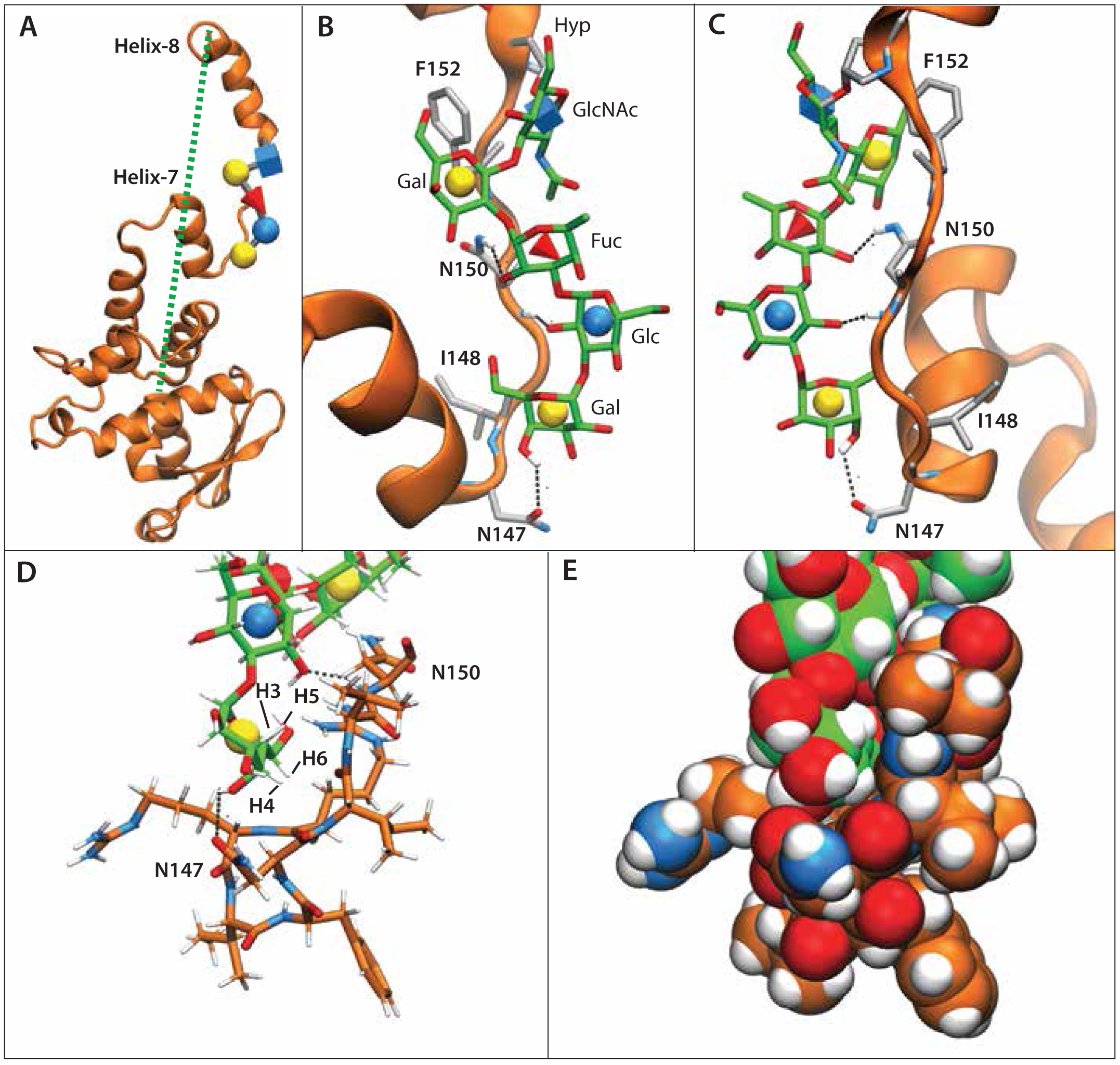
Conformational analysis of glycosylated TgSkp1 by MD simulation. GaGlFGaGn-Skp1 was subjected to six 250-ns all-atoms molecular dynamics simulations. (**A**) A frame representative of the glycan-protein interaction and associated helix-8 extension, from a simulation (Equil-1,see Fig. S12E) in which the glycan was pre-equilibrated for 50 ns prior to the start of the simulation. The dotted green line refers to the distance from C-terminus to the center of mass of residues 1-136, and ranged from 18 to 61 Å. (**B**) Zoom-in of panel A depicting the glycan (C-atoms in green) and amino acids (C-atoms in gray) described in Table 2. Dotted black lines depict H-bonds contributing to the polar energies described in Table 2. (**C**) The back side of panel B is depicted. **(D,E)** Packing of terminal sugars against the polypeptide. C-atoms of the peptide are in orange. **(D)** Glycan and peptide represented as sticks. **(E)** As in D, with Glycan and peptide represented by spheres.

**Table 2.**
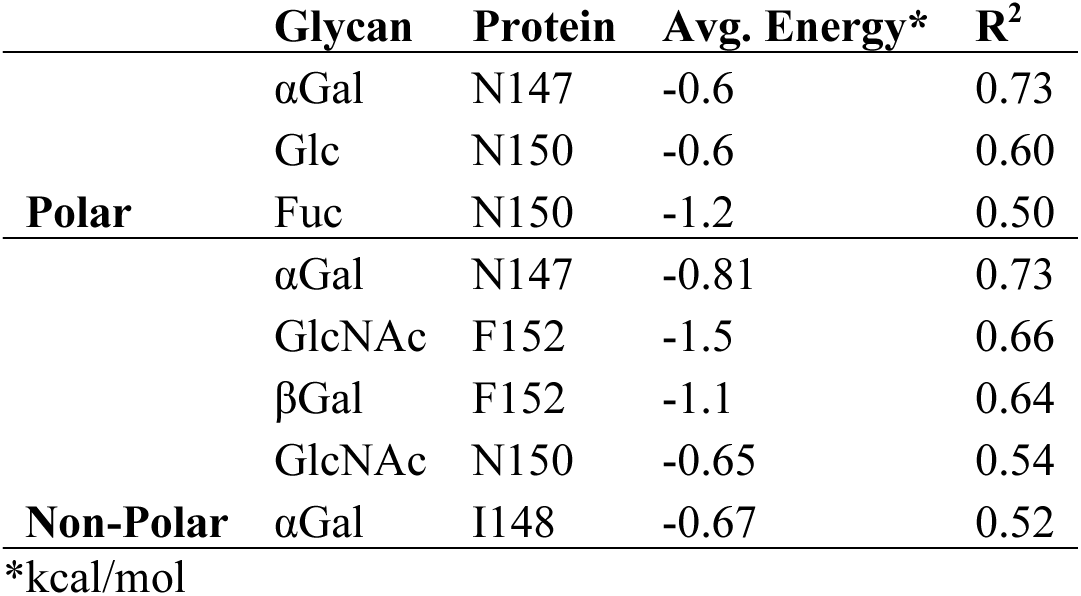
Table of MMGBSA-derived per-residue energies between the protein and glycan that exhibit a strong correlation with helix extension (distance in Fig. 6A) according to a linear regression analysis of 48 bins from the six MD simulations (Fig. S12E). Polar energies represent a sum of the electrostatic and polar solvation energies while the nonpolar energies are composed of the van der Waals and nonpolar solvation energies. Only interactions with an average polar/non-polar energy less than −0.5 were considered.

A linear regression analysis identified correlations between helix distance and predicted polar and nonpolar interaction energies with monosaccharides in the glycan (Table 2). The results indicate that extension of the C-terminus of TgSkp1 involves both hydrogen bonds and van der Waals interactions between sugar and amino acid residues. The strongest correlated polar interactions between the glycan and Skp1 (Table 2) correspond to three hydrogen bonds to amino acids within the unstructured peptide chain located between helix 7 and 8 (residues 147-152), as depicted in Figs. 6B, 6C. The correlations with nonpolar interactions also occur within this region, with the terminal αGal moiety showing the highest correlation (Table 2). To illustrate this correlation, the local environment near the αGal moiety was examined using the representative frame of the average structure (Fig. 6A). The hydrophobic face of the terminal αGal can pack against a nonpolar pocket consisting of planar faces of peptide backbone (Figs. 6D, 6E). Specifically, its C3-C5 H-atoms can pack with the backbone of residues 147-149, and a hydrogen on C6 can be buried within residues 143 and 144 (Fig. 6D). Both the polar and nonpolar interactions identified here involve amino acids that are conserved across both organisms, with the exception of V149 which is Lys in DdSkp1 (Fig. S12D). This substitution would not be expected to affect Gal burial since the majority of interactions are with the protein backbone at this position (not shown). The previously noted hydrogen bonds-1 and −3, involving E140 (E129 in DdSkp1), were also observed in this study (Fig. S12C), but were poorly correlated (R^2^=0.0, vs. 0.63 average for the other three) with helix extension (Fig. S12E). The energetic analyses applied to the glycan variant found in *Toxoplasma* confirm that, despite its sequence difference, the glycan forms an organized structure with the potential to influence local Skp1 polypeptide organization. The new studies emphasize the interaction of the glycan with the interhelix region and contributions of packing of the terminal sugars including the αGal. As previously hypothesized, these effects have the potential to influence the conformational ensemble of TgSkp1 to be more receptive to F-box protein binding.

### Structural relationship of Gat1 to glycogenin

To further probe the relationship between Gat1 and glycogenin, we compared their structures by X-ray crystallography. Attempts to crystallize TgGat1 were unsuccessful, even after deletion of its unconserved insert (Fig. 3A). However, PuGat1, which lacks this insert, was co-crystallized in the presence of Mn^2+^ and UDP.

The crystal structure of PuGat1 in complex with Mn^2+^ and UDP was solved using single-wavelength anomalous dispersion phasing of a Pt^2+^ derivative, and the resolution was extended to 1.76 Å using a native data set (Table S3). The asymmetric unit contains a single chain of PuGat1 with unambiguous electron density for the nucleotide and Mn^2+^ ion (Fig. 7A). The overall structure of PuGat1 reveals a canonical GT-A fold (Bourne and Henrissat, 2001) consisting of eight α-helices and eight β-sheets. The N-terminus (residues 1-8) and two loops (residues 80-96 and 242-244) are disordered and were not modeled. The structure is similar to glycogenin-1 from *Oryctolagus cuniculus* (Oc*-*glycogenin-1), which superimposes 213 corresponding C*α* atoms with an RMSD of 3.3 Å despite a sequence identity of only 34% (Fig. 7B). The application of crystallographic symmetry shows that PuGat1 forms the same dimer described (Gibbons et al., 2002) for the Oc*-*glycogenin-1 structure (Fig. 7B). According to PISA (Krissenel and Henrick, 2007) analysis, the PuGat1 dimer interface buries 1090 Å^2^ with a favorable P-value of 0.107, which suggests that the dimer contact is stable. Sedimentation velocity analysis of 3.5 μM PuGat1 reveals a *c(s)* distribution consisting of single species at 4.0 S, which corresponds to the predicted value of 4.2 S for a dimer (Fig. 7C). The slightly slower sedimentation indicates that the enzyme in solution is less compact than that observed in the crystal structure. PuGat1 was dimeric even at 0.3 µM (Fig. S14), suggesting that it forms a dimer with an affinity >2-fold higher than that of Oc-glycogenin-1 (which has a reported *K_d_* of 0.85 µM) (Bazan et al., 2008). Based on gel filtration and preliminary sedimentation velocity experiments (not shown), TgGat1 is also a dimer.

**Fig. 7.**
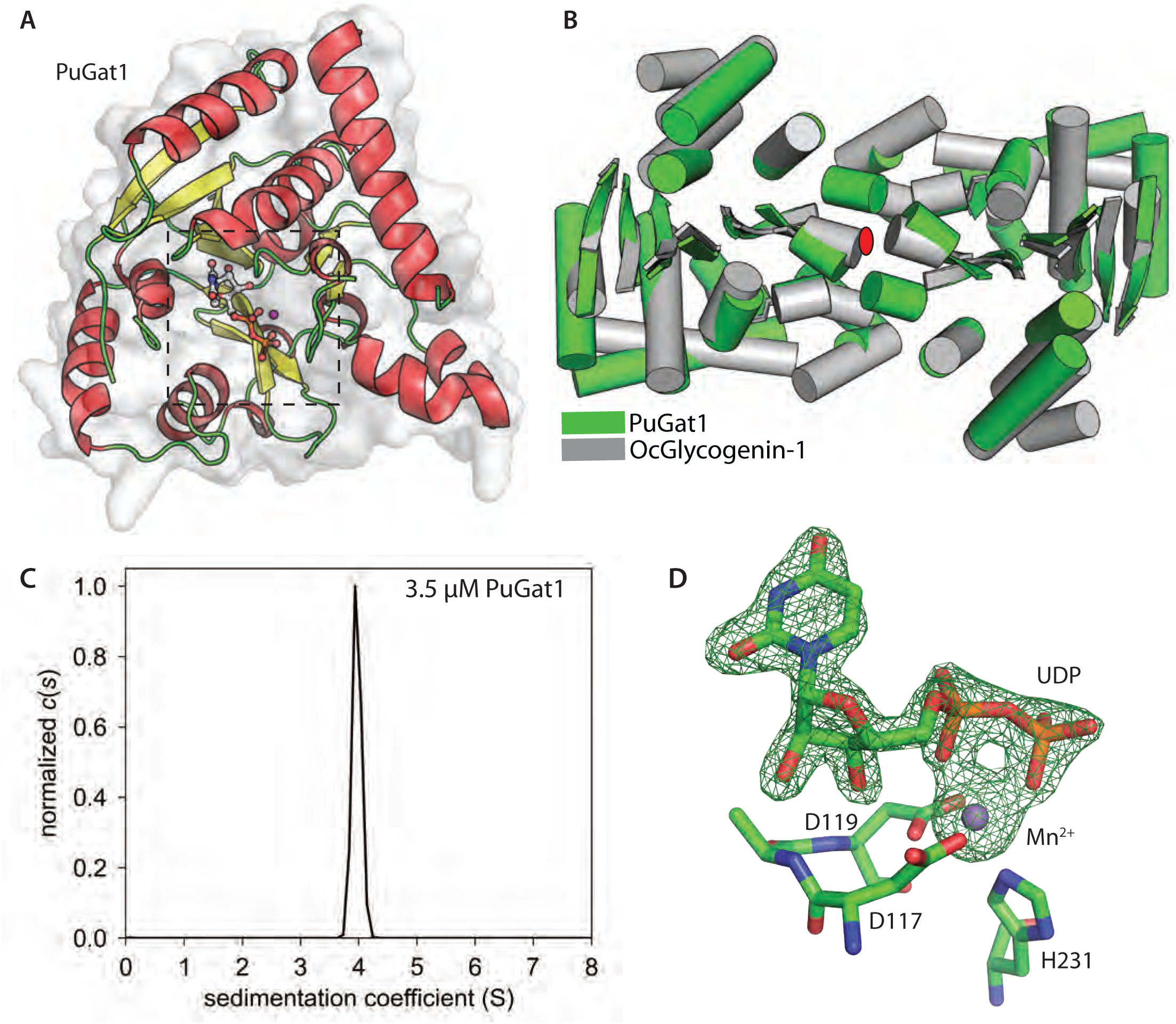
Crystal structure of PuGat1 and its oligomeric state in solution. (**A**) The asymmetric unit of PuGat1:UDP:Mn^2+^ is shown with α-helices in red, β-strands in yellow, and loops in green. Secondary structures are assigned based on DSSP. The box highlights the active site with bound ligands. (**B**) PuGat1 and Oc-glycogenin-1 (PDB entry 1LL2) dimers are superimposed. The cylinders represent α-helices, and the arrows represent β-sheets. The red ellipse is the two-fold symmetry perpendicular to the page. (**C**) Sedimentation velocity experiment of 3.5 μM PuGat1 is displayed as a continuous *c*(*s*) distribution (normalized to 1.0). The black dashed represent the predicted dimer S-values based on the crystal structure. See Fig. S14 for additional data. (**D**) The difference density map (*F_o_-F_c_*) contoured at 5σ was calculated after omitting UDP and Mn^2+^ and subjecting the structure to simulated annealing Octahedral coordination of Mn^2+^ is satisfied by Asp117 and Asp119 of DxD motif, His231, and the *α* and *β* phosphates of UDP. See Fig. S13 for additional information.

The PuGat1 active site shows that the conserved DxD motif (Bourne and Henrissat, 2001) and a conserved His residue coordinate the Mn^2+^ ion using the Oδ2 atom of D117, both Oδ1 and Oδ2 atoms of D119, and Nε2 atom of His231 (Fig. 7D). The Mn^2+^ ion is also coordinated by the oxygen atoms from the *α* and *β* phosphates of UDP. Comparing the PuGat1 and Oc-glycogenin-1 active sites shows that all of the interactions with the nucleotide are conserved, with the exception of the interactions with N3 and O4 of the uracil ring (Fig. S13). Other changes in Gat1 active site include a Leu to Ser substitution at residue 233, which can potentially remove a packing interaction with the donor sugar in PuGat1 (Fig. 8). The hydroxyl of the substituted Ser233 forms a hydrogen bond with the adjacent side chain of Gln206. A water molecule (W509) replaces the Leu side chain and forms a hydrogen bond with Asn149.

**Fig. 8.**
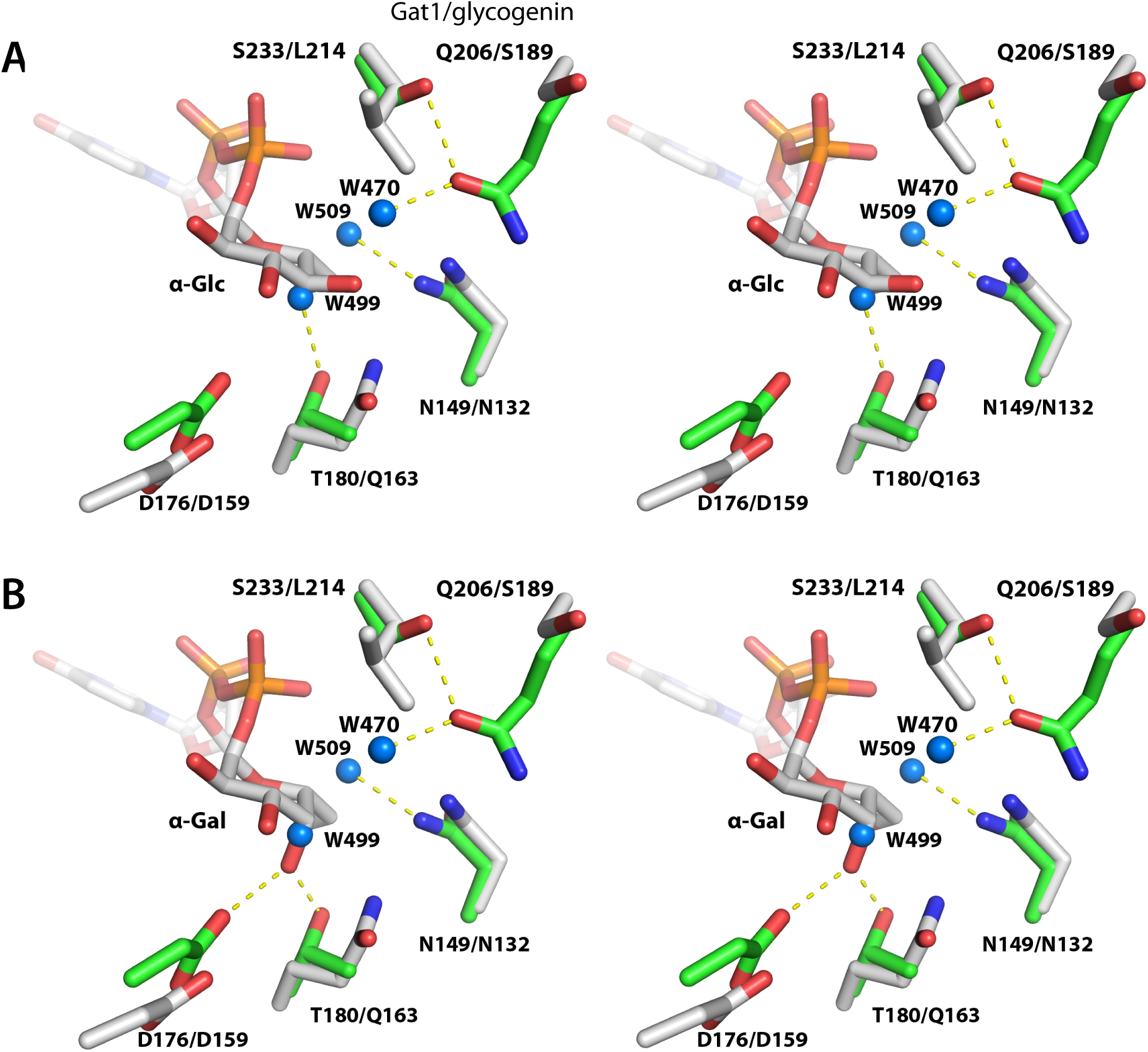
Active site comparison of PuGat1 and Oc*-*glycogenin-1. Comparison of the sugar binding pockets of PuGat1 and Oc-glycogenin-1 displayed as wall-eyed stereoview. (**A)** The Glc moiety is modeled based on the Oc-glycogenin-1 crystal structure with its intact sugar nucleotide. **(B)** The Gal moiety is modeled by flipping the stereochemistry of Glc at the C4’ position. PuGat1 and glycogenin-1 side chains are represented by green and gray sticks, respectively, and the yellow dashes represent hydrogen bonds. Water molecules are represented by blue spheres.

Comparing the crystal structure of UDP-Glc bound Oc-glycogenin-1 with PuGat1 immediately suggests a reason for why these enzymes have different donor specificities (Fig. 8). Superimposing the Oc-glycogenin-1:UDP-Glc structure onto PuGat1 shows that UDP-Glc would displace water499 coordinated by Thr180 (Fig. 8A). This would leave the Thr180 hydroxyl group unsatisfied, and the unfavorable burial of a polar group likely explains why UDP-Glc is a poor donor (Fig. 8A). In contrast, we modelled in UDP-Gal by flipping the stereochemistry at C4 position. The O4 atom of Gal would be ideally positioned to satisfy the Thr180 hydroxyl group. Concurrently, Asp176, whose Cα atom underwent a 2.3 Å shift relative to its location in glycogenin, would be in position to receive a hydrogen bond from the O4 atom of the Gal (Fig. 8B). Altogether, Gat1’s sugar donor preference for UDP-Gal is likely due to formation of favorable hydrogen bonds with O4 atom of Gal in contrast to the burial of Thr180 hydroxyl group when binding UDP-Glc.

### Computational modeling of acceptor binding

To address the basis of Gat1’s preference for the GlFGaGn-glycan, the lowest energy conformation of the reducing form of the glycan generated by GLYCAM-web (www.glycam.org) was docked using AutoDock Vina. A plausible docking mode was selected based on the requirement that the C’3-hydroxyl group must be oriented towards the anomeric carbon of the donor sugar to serve as the nucleophile for addition of the Gal, and that the glycan does not clash with the other subunit of the dimer. Out of 100 docking simulations, only the top scoring pose with a binding energy score of −5.7 kcal/mol satisfied the selection requirement. In this pose, the glycan adopted an alignment in a groove formed by Gat1 dimerization (Figs. 9A, B). The glycan is stabilized by hydrogen bond contributions from the sidechains or peptide backbones of residues D141, F143, and S234 from subunit A, and residues L212, K216, N217, and Y220 from subunit (Fig. 9C). In addition, non-polar interactions against the faces of the sugar moieties are provided by residues T208, L212, F143, and W221 from subunit A, and residue Y220 from subunit B (Fig. 9D). F143, Y220, W221, and S234 are uniquely conserved in Gat1 proteins relative to glycogenins (Fig. S6). The packing interactions from conserved hydrophobic residues are likely the major contributors in terms of binding energy. The extensive electrostatic and packing complementarity, which could not be achieved using the same approach using the *α*4Glc-tertramer recognized by glycogenin, can explain the distinct preference of Gat1 for the Skp1 tetrasaccharide acceptor substrate.

**Fig. 9.**
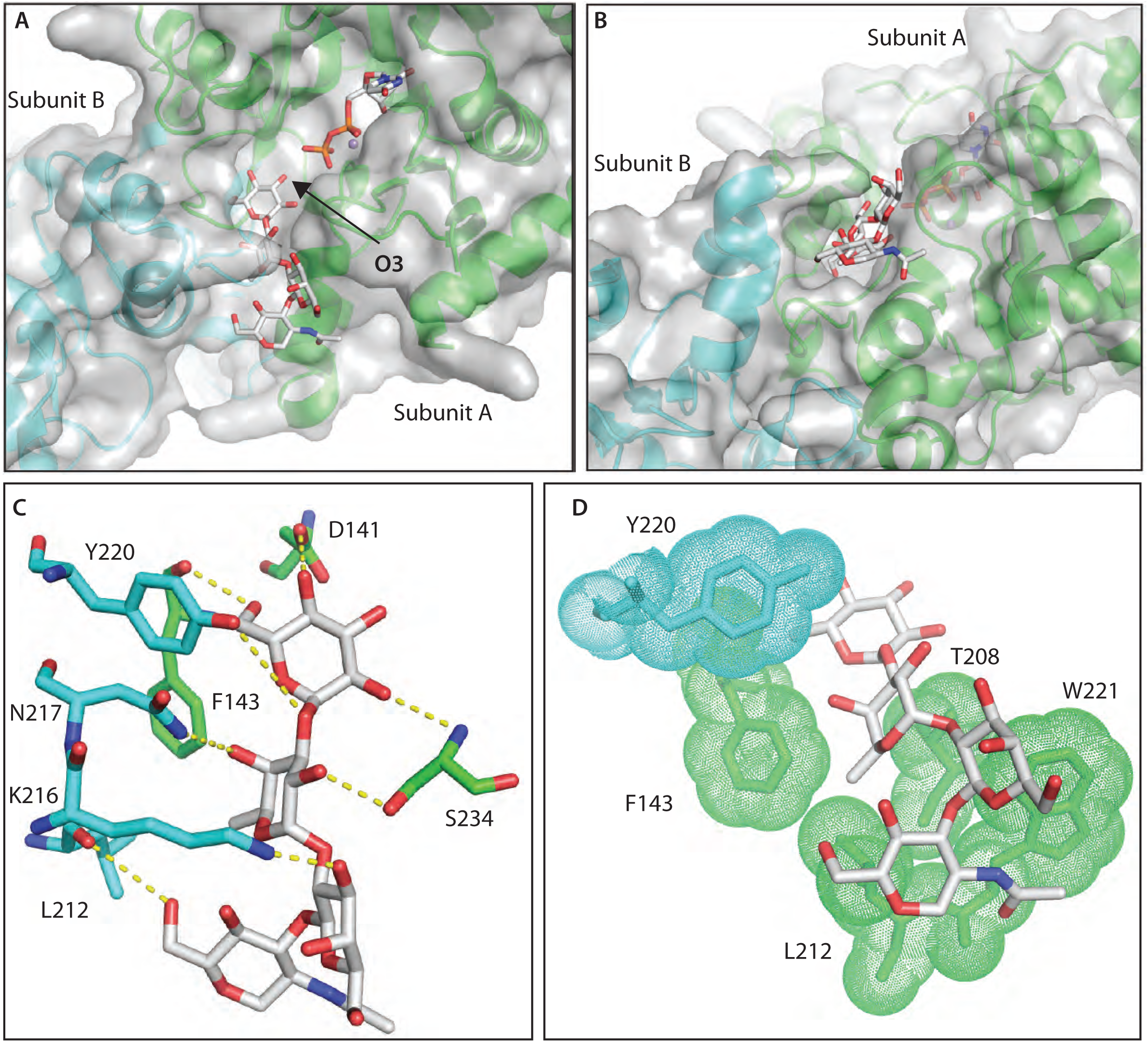
Top docking pose reveals Gat1 and Skp1-tetrasaccharide acceptor complementarity. (**A**) Skp1-tetrasaccharide binds in the PuGat1 active site and is accommodated by a groove formed by the dimer. (**B**) 90° turn of the image shown in panel A. (**C**) Hydrogen bonding interaction with the glycan is shown with the residues/ligand in sticks. (**D**) Hydrophobic packing with the faces of the sugar moieties and the methyl moiety of fucose is shown with residues in sticks/dots and ligand in sticks. Gat1 subunit A is represented in green, and subunit B in cyan.

## DISCUSSION

Gat1 is unusual for glycosyltransferases. First, it appears to be dedicated to the glycosylation of a single target protein, Skp1. Second, it resides in the cytoplasmic compartment, rather than the secretory pathway, of the cell. Third, it is widespread among protists, but not found outside of the kingdom. Fourth, it substitutes for another unrelated glycosyltransferase that modifies Skp1 in amoebozoa, but appears to contribute a related regulatory function. Nevertheless, it is descended from a widely distributed lineage of sugar nucleotide-dependent glycosyltransferases. It is most closely related to glycogenin, a sublineage of retaining glycosyltransferases that potentially evolved from Gat1 to modulate glycogen formation outside of the protist kingdom, in yeast, fungi and animals.

Recombinant Gat1s from *T. gondii* and *P. ultimum* are CAZy GT8 family enzymes that, *in vitro*, can catalyze the transfer of αGal from UDP-Gal to non-reducing terminal Glc acceptors in an α1,3-linkage, as determined using NMR analysis of a synthetic version of the TgSkp1 glycan as an acceptor (Fig. S11). The enzyme was highly selective for UDP-Gal as the donor, but was able to transfer Glc from UDP-Glc at low efficiency (Figs. 4, S9). The enzyme also preferred the TgSkp1 tetrasaccharide relative to other mono- and oligo-saccharides, but is able to modify any non-reducing terminal Glc residue, in α- or β-configuration. These reactions are characterized by *K*_m_ values in the range of 70 µM for UDP-Gal and 1.5 mM for the free TgSkp1 acceptor glycan, and exhibit pH and ionic strength optima that are consistent with action in the cytoplasmic compartment. In side-by-side comparisons, no substantial differences were found for Gat1’s from *T. gondii* and *P. ultimum*, indicating a high degree of conservation despite the wide phylogenetic distance between these two species.

In *T. gondii*, the *gat1* gene is required for addition of the terminal hexose of TgSkp1. Since the terminal hexose is αGal, based on its removal with green coffee bean α-galactosidase (Fig. 1C), and because Gat1 can efficiently and selectively catalyze the corresponding reaction *in vitro* (Fig. 4), it is concluded that Gat1 is directly responsible for this modification *in vivo*. No other Gat1 targets were detected by biochemical complementation of *gat1*Δ extracts (Figs. 4J, S9H, I). Because Gat1 is related to glycogenin, which is involved in glycogen formation in yeast and animals, amylopectin (starch) formation was examined in bradyzoites. However, there was no evidence for an effect on its accumulation (Fig. 5). These findings are consistent with Gat1’s preference for Skp1 glycan-related acceptors *in vitro* (Fig. 4F) and modeling of acceptor oligosaccharides into the active site of the PuGat1 crystal structure (Fig. 9). Furthermore, Gat1 was even more reactive when the glycan was attached to Skp1 (Fig. 4H, I), indicating that the apoprotein contributes to enzyme recognition. These results are consistent with evidence that the underlying glycosyltransferases are also specific for Skp1 in *T. gondii* (Rahman et al., 2016; Rahman et al., 2017). In *D. discoideum*, the conclusion of specificity for Skp1 is strengthened by findings that genetic manipulations of Skp1 levels interact genetically with its glycosylation with respect to O_2_-sensing (Wang et al., 2011).

The slow growth phenotype of *gat1*Δ parasites in the monolayer plaque assay is likely attributable to the absence of Gat1 because similar results were obtained in independent knockouts in three different genetic backgrounds, and the defect was corrected by genetic complementation under its own promoter or a strong tubulin promoter in the *uprt* locus (Fig. 2). Since the only detectable substrate for Gat1’s enzymatic activity was GlFGaGn-Skp1, it is likely that failure to fully glycosylate Skp1 is responsible for the growth defect. This interpretation is consistent with similar effects of knocking out early GTs in the pathway, but the phenotype is milder than the *phyA*Δ phenotype (Rahman et al., 2016; Rahman et al., 2017; Xu et al., 2012). However, we cannot rule out some other function of Gat1, though we note that at 345 amino acids, nearly all of its sequence appears devoted to the enzymatic domain (Fig. 3A). Future studies will be needed to address whether Gat1 contributes to fibroblast adhesion, invasion, proliferation within the parasitophorous vacuole, or egression from fibroblasts.

The mechanism by which full glycosylation affects Skp1 shows similarities to the model proposed for *D. discoideum*. According to the previous NMR relaxation studies and molecular dynamics simulations, the glycan adopts a rather restricted conformation which interacts with an intrinsically disordered region of the Skp1 polypeptide, resulting in an opening of F-box binding region (Sheikh et al., 2017). Lowest energy conformation calculations and MD simulations indicated that the TgSkp1 glycan, despite the presence of Glc rather than Gal at the fourth position, adopts a similar conformation (Fig. S12A,B). The all-atoms simulations of TgSkp1 also revealed a dynamic relationship of the glycan with the intrinsically disordered region upstream of its attachment site, that was mediated, according to a new residue-by-residue energetics analysis, by both non-polar and polar contacts (Fig. 6; Table 2). In particular, the new analysis emphasized the importance of the fifth sugar, whose addition is mediated by Gat1, in contributing substantial polar and non-polar contacts to stabilize the glycan relationship (Fig. 6D,E). In addition, the conservation of hydrogen bonds confirmed the predicted orientations of individual sugars with the polypeptide (Fig. S12C). A time resolved analysis of the presence of these non-polar and polar interactions with extension of helix-8 and the C-terminus of Skp1 showed a strong correlation (Table 2; Fig. S12E), suggesting a causal relationship in preparing Skp1 to interact with F-box domains.

The importance of the fifth sugar is reinforced by the observation that the amoebozoa, which lack both Glt1 and Gat1 (Figs. 1A, B), evolved a new unrelated enzyme, AgtA, to fulfill the role. AgtA is a dual function GT that applies both the 4^th^ and 5^th^ sugars: each an α3linked Gal. This raises the interesting possibility that AgtA evolved to compensate for the unexplained loss of Glt1 and Gat1, with recursive addition of the same sugar being an accessible evolutionary pathway to recover the pentasaccharide.

The most closely related protein that we found in genomic sequence databases is glycogenin (Fig. 3), the α4-glucosyltransferase that can autoglucosylate a specific Tyr near its active site and then form a linear chain of up to 12 residues that may prime glycogen synthesis in fungi and metazoa. Conversely, the Gat1 sequence is observed in a range of alveolates (includes *T. gondii*), stramenopiles (includes *P. ultimum*), and rhizaria (altogether known as the SAR group), and archaeplastids (with the SAR group known as bikonts), but not in the more recently emerging unikonts that include the amoebozoa, fungi and animals (Burki, 2014; Brown et al., 2018). Since glycogenin has the complementary distribution being found only in fungi and animals, it is tempting to speculate that Gat1 is the evolutionary precursor of glycogenin. This concept is supported by our failure to find both sequences in the same group of organisms in extant databases (Fig. 3). The aforementioned loss of Glt1 might have created the opportunity for Gat1 to evolve to a new biochemical role, and it is interesting to speculate whether this conserved a cellular function.

The hypothesis that Gat1 is the evolutionary precursor of glycogenin is strengthened by comparison of their crystal structures. The dimer interface of Gat1 precisely overlaps that of rabbit glycogenin-1 (Figs. 7, S14). The UDP moiety of the sugar nucleotide donor is coordinated in the same manner in the active site (Fig. S13). Residues that recognize the Glc-moiety of UDP-Glc in glycogenin-1 are distinct in Gat1 in a way that can explain the preference for UDP-Gal (Fig. 8) while allowing inefficient processing of UDP-Glc as observed in the *in vitro* reactions (Fig. 4B-D). This duality presages the recently discovered ability of glycogenin to utilize UDP-Gal during intermediate phases of glucan extension (Bilyard et al., 2018). Interestingly, UDP-Glc is able to inhibit the αGalT activity of Gat1 at concentrations that may be achieved *in vivo* (Fig. 4G), potentially explaining why this degeneracy is retained. On the other hand, presentation of the acceptor oligosaccharide is distinct. Computational modeling suggests that the dimer interface presents a unique opportunity for insertion of the Skp1 tetrasaccharide (Fig. 9), in a manner that is incompatible with docking of an α4-glucan. The 3-OH of the acceptor Glc is positioned appropriately to act as the nucleophile. If glycogenin evolved from Gat1, substantial changes occurred to allow both auto-glucosylation and extension of the initial Tyr-linked αGlc with additional α4-linked Glc residues in a manner that evidently depends on *trans*-interactions across a pre-existing dimer interface (Issoglio et al., 2012). Although the target Tyr-residue is present in a similar position in some Gat1’s, this immediate region is poorly conserved among Gat1’s (Fig. S6). Possibly, these features provided an evolutionary opportunity for the novel mechanism adopted by glycogenin once it was free to evolve new roles following the evident disappearance of Glt1. In addition, glycogenin acquired a C-terminal glycogen synthase binding domain of unknown function (Fig. 3A; Zequiraj et al., 2014)

Past and recent studies indicate that, despite the compelling *in vitro* data, glycogenin is not required for glycogen synthesis in yeast and mice (Torija et al., 2005; Testoni et al., 2017). Nevertheless, glycogen levels are affected by glycogenin by an unknown mechanism. This role was evidently acquired later in evolution because Gat1 does not appear to have a significant effect on amylopectin accumulation in *T. gondii,* and *gat1* is expressed equally in starch-poor tachyzoites and starch-rich bradyzoites based on transcript analysis (Coppin et al., 2005). Although the role of this enzyme lineage in Skp1 modification was lost by the time amoebozoa evolved *en route* to the appearance of fungi/yeast and animals, it is possible that it retains a currently unknown *trans* glycosyltransferase activity. Future studies to investigate additional potential targets of glycogenin in metazoa are warranted.

## EXPERIMENTAL PROCEDURES

### Maintenance of host cells and parasite manipulations

Cultures of human foreskin fibroblast (HFF, ATCC SCRC-1041) or hTERT HFF (BJ-5ta, ATCC CRL-4001) were maintained Dulbecco’s modified Eagle’s Medium supplemented with 10% (v/v) fetal bovine serum, 2 mM L-glutamine and 100 units/ml penicillin/streptomycin (Corning) at 37°C in a humidified CO_2_ (5%) incubator. Type 1 RH (Sabin, 1941), RHΔ*Ku80*Δ*HXGPRT* (RHΔΔ) (Fox et al., 2009) and type 2 ME49-RFP (Grainger et al., 2013) strains of *Toxoplasma gondii* were cultured on HFF or hTERT HFF monolayers in the same medium as described for host cells except that 1% (v/v) fetal bovine serum was used where stated, and cloned by limiting dilution in 96-well plates. Parasites were kept in media without drug for plaque assays, which were performed as before (Gas-Pascual et al., 2019). Transfections were conducted by electroporation using a BioRad Gene Pulser Xcell at 1.5 kV and 25 µF with 2 mm electroporation cuvettes (VWR international) in Cytomix buffer (10 mM KH_2_PO_4_/K_2_HPO_4_ (pH 7.6), 120 mM KCl, 0.15 mM CaCl_2_, 5 mM MgCl_2_, 25 mM HEPES, 2 mM EDTA).

### Disruption and complementation of *gat1*

TGGT1_310400 (Toxodb.org), referred to as *gat1* (Fig. S1), was disrupted by two independent approaches. In the first method, a disruption DNA was prepared from the vector pmini-GFP.*ht* (a gift from Dr. Gustavo Arrizabalaga) in which the *hxgprt* gene is flanked by multiple cloning sites as described (Rahman et al., 2016). 5’-flank and 3’-flank targeting sequences were PCR amplified from strain RHΔΔ with primer pairs Fa and Ra and pairs Fb and Rb, respectively (Table S1). The 5’-fragment was digested with ApaI and XhoI and cloned into similarly digested pminiGFP.ht. The resulting plasmid was digested with XbaI and NotI and ligated to the similarly digested 3’-flank DNA. The resulting vector was linearized with PacI and electroporated into strain RHΔΔ, selected under 25 µg/ml mycophenolic acid and 25 µg/ml xanthine, and cloned by limiting dilution. Genomic DNA was screened by PCR to identify Tg*gat1* disruption clones (Fig. S3A), using primers listed in Table S1.

The second approach was based on a double-CRISPR/Cas9 method as previously detailed (Gas-Pascual et al., 2019), with minor modifications. To generate the dual guide (DG) plasmid, a fragment of p2 containing the guide RNA gat1-63 expression cassette was PCR amplified using primers plasmid 3 FOR and plasmid 3 REV (Table S1), digested with NsiI, and ligated into the NsiI site of a dephosphorylated p3 containing guide RNA gat1-968. The type 1 RH and type 2 ME49 strains were co-transfected with pDG-Gat1 (10 µg) and a dihydroxyfolate reductase (*dhfr*) amplicon (1 µg) by electroporation (Fig. S4A). CRISPR/Cas9-mediated disruption in RHΔΔ parasites was done similarly except that pDG-Gat1 was co transfected with a *dhfr* amplicon containing 45 bp homology arms targeting *gat1* (Fig. S3B). *Gat*1Δ parasites were subsequently selected in 1 µM pyrimethamine (Sigma). The expected replacement of nt 63-968 (relative to A of ATG start codon) with the *dhfr* cassette was confirmed by PCR using primers listed in Table S1.

To complement *gat1*Δ RH parasites, a Tg*gat1* DNA fragment consisting of the coding sequence of *gat1* plus approximately 1.2 kb each of 5’- and 3’-flanking DNA sequences was generated by PCR from RH genomic DNA using primers Fa and Rb, which contained Apa1 and Not1 restriction sites respectively. After treatment with Apa1 and Not1, the PCR product was cloned into similarly digested pmini-GFP-*ht* plasmid in place of its *hxgprt* cassette, to generate pmini-Tg*gat1*. The plasmid was transformed into *E. coli* Top10 cells and purified by using a Monarch miniprep kit (NEB). A Ty tag DNA sequence was inserted at the 3’-end of the *gat1* coding sequence using a Q5 site directed mutagenesis kit (NEB) and Fn and Rn primers, yielding pmini-Tg*gat1*-Ty. The sequence was confirmed using primers Fl and Rl (Table S1). RH *gat1*Δ clone 8 was complemented by co-electroporation with a PCR amplicon from pmini-Tg*gat1-Ty* (1µg) and a sgUPRT CRISPR/Cas9 plasmid (10 µg) targeting the *uprt* locus using the guide DNA sequence 5’-ggcgtctcgattgtgagagc (Shen et al., 2014) (Fig. S4B). Transformants were selected with 10 µM fluorodeoxyuridine (FUdR, Sigma), and drug resistant clones were screened by PCR with primers listed in Table S1.

To complement *gat1* in RHΔΔ parasites, the UPRT Vha1 cDNA shuttle vector containing TgVhaI cDNA (Stasic et al., 2019) was modified to generate a Gat1-HA complementation plasmid using NEB HiFi Builder method. The vector backbone, containing 5’-flank and 3’-flank *uprt* targeting sequences and a Tg-tubulin promoter and 3×HA sequence, was PCR amplified from the shuttle vector using primers Ft and Rt. The coding sequence of *Tg*Gat1 was PCR amplified from pmini-Tg*gat1* plasmid using primers Fu and Ru, which had 18-21 nts complementary to the terminal ends of the vector (Fig. S3C). The gel-purified PCR fragments were incubated with HiFi DNA assembly enzyme mix (NEB) and transformed into *E. coli* Top10 cells, yielding pUPRT*gat1*3xHA. The Tgg*at1* sequence was confirmed using primers Fl and Rl. Complementation in a *gat1*Δ clone derived from RHΔΔ was done similarly except that a *gat1*-3xHA PCR amplicon with 5’and 3’ *uprt* homology arms was used in the transfection.

### Bradyzoite induction

ME49-RFP tachyzoites were differentiated to bradyzoites using alkaline pH (Soête et al., 1994). HFF monolayers were pre-incubated with sodium bicarbonate free RPMI (Corning) containing 50 mM HEPES-NaOH (pH 8.1) for 24 h, and then infected with tachyzoites and maintained at ambient atmosphere at 37°C with medium replacement every 24 h. Differentiation was monitored by labeling with *Dolichos biflorus* agglutinin (DBA) (Zhang et al., 2001). Infected HFF monolayers formed on 25 mm coverslips were washed with PBS (Corning), fixed with 4% paraformaldehyde in PBS for 10 min, washed with PBS, and permeabilized with 1% Triton X-100 (BioRad) in PBS for 10 min, all at room temperature. Samples were blocked with 3% (w/v) bovine serum albumin (BSA) in PBS for 1 h, incubated for 2 h with 5 μg/ml FITC-DBA lectin (Vector Laboratories Inc.) in 1% BSA in PBS, and washed with PBS. The coverslips were mounted with ProLong Gold antifade reagent (Invitrogen) on glass slides. The slides were imaged by phase contrast and fluorescence microscopy on a Zeiss Axioskop 2 Mot plus.

### Periodic acid staining

To assess amylopectin levels, parasite-infected HFF monolayers on 25 mm glass coverslips were washed with plain PBS, fixed with ice-cold MeOH for 5 min, washed with PBS, and incubated with 1% periodic acid in deionized H_2_O in the dark for 10 min (Dubey et al., 1998). The coverslips were washed with deionized H_2_O, incubated with Schiff reagent for 15 min, washed once with deionized H_2_O, and finally rinsed with running tap water for 10 min. The stained coverslips were dehydrated by sequential immersion in 70% (v/v), 80%, 90% and 100% EtOH, mounted on glass slides using Permount mounting media (Fisher Scientific), and imaged on an EVOS XL Core microscope (Invitrogen).

### Expression and purification of recombinant TgGat1 and PuGat1

The single exon coding sequence of Tg*gat1* cDNA was amplified by PCR from RH genomic DNA using primers, Gat1 Fw and Gat1 Rv (Table S1), cloned into PCR4-TOPO TA (Invitrogen), and transformed into *E. coli* Top 10 cells. The plasmid was double digested with BamH1 and Nhe1 to yield the *gat1* fragment that was cloned into similarly digested pET15-TEVi plasmid (Invitrogen), resulting in the original 346 amino acid coding sequence of Gat1 extended at its N-terminus with a His_6_-tag and TEV protease cleavage site (MGSSHHHHHHSSGRENLYFQGH-). A similar N-terminal modification of rabbit glycogenin did not significantly alter its enzymatic activity (Cao et al., 1995). The predicted coding sequence for Pu*gat1* was inferred from PYU1_G002535-201 at protists.ensembl.org/Pythium_ultimum. The coding sequence was codon optimized for *E. coli* expression, chemically synthesized by Norclone Biotech (Ontario, Canada), and inserted in the NdeI and XhoI sites of pET15b-TEV. The expressed protein was extended at its N-terminus with MGSSHHHHHHSSGENLYFQGH-.

TgGat1 or PuGat1 were expressed in and purified from *E. coli* BL21-Gold cells as previously described for TgGlt1 (Rahman et al., 2017), through the purification on a 5-ml Co^+2^ TALON resin column. The eluted protein was dialyzed in 50 mM Tris-HCl (pH 8.0), 300 mM NaCl, 1 mM EDTA and 2 mM β-mercaptoethanol, followed by 25 mM Tris-HCl (pH 8.0), 200 mM NaCl, 1 mM β-mercaptoethanol and 5 mM MnCl_2_. The sample was treated with 2 µM His_6_-TEV protease, 5 µM TCEP in the same buffer overnight at 20°C, and reapplied to another Co^+2^ TALON column. The flow-through fraction was dialyzed in 25 mM Tris-HCl (pH 8.0), 50 mM NaCl, 1 mM β-mercaptoethanol, 2 mM MnCl_2_. The sample was concentrated by centrifugal ultrafiltration and aliquots were stored at −80°C. As indicated, the preparations were further purified by gel filtration on a Superdex200 column in the same buffer.

### SDS-PAGE and Western blotting

Samples were suspended in diluted with Laemmli sample buffer and typically electrophoresed on a 4–12% gradient SDS-polyacrylamide gel (NuPAGE Novex, Invitrogen). Gels were either stained with Coomassie blue or transferred to a nitrocellulose membrane using an iBlot system (Invitrogen). Blots were typically blocked in 5% non-fat dry milk in Tris-buffered saline and probed with a 1:1000-fold dilution of the antibody of interest in the milk solution, followed by secondary probing with a 1:10,000-fold dilution of Alexa-680-labeled goat anti-rabbit IgG secondary antibody (Invitrogen). Blots were imaged on a Li-Cor Odyssey infrared scanner and analyzed in Adobe Photoshop with no contrast enhancement. For measuring incorporation of radioactivity, 1-mm thick 7-20% acrylamide gels were prepared manually. Detailed information is found in Rahman et al., (2016).

### Preparation of Skp1 peptides

To monitor Skp1 glycosylation status, endogenous TgSkp1 was purified from parasite extracts essentially as described (Rahman et al., 2016). Briefly, frozen pelleted tachyzoites (1 × 10^8^) were resuspended in 8 M urea in 50 mM HEPES-NaOH (pH 7.4), incubated on ice for 1 h and at 50**°**C for 5 mins, and diluted 8-fold in IP buffer (0.2% Nonidet P-40 in 50 mM HEPES-NaOH, pH 7.4). The lysates were centrifuged at 21,000 × *g* for 20 min at 4**°**C, and 100 µl of the supernatant (5 × 10^7^ cells) were incubated with 5 µl UOK75 rabbit antibody (anti-TgSkp1) coupled to protein A/G magnetic agarose beads (Pierce, 78609) for 1 h at 4**°**C. Beads were captured in a DynaMag-2 magnet (Life Technologies) according to the manufacturer’s directions, and washed 3× with 50 mM HEPES-NaOH (pH 7.4), 3× with 10 mM Tris-HCl (pH 7.4), 50 mM NaCl, and once with 50 mM NaCl. Bound Skp1 was eluted twice with 60 µl 133 mM triethylamine (TEA, Sequencing Grade, Pierce, 25108) by incubating for 15 mins at RT, and immediately neutralized with 40 µl of 0.2 M acetic acid. The eluted material was pooled and dried under vacuum, reconstituted in 8 M urea, 10 mM Tris HCl (pH 7.4), reduced in 10 mM DTT for 40 min at RT, and alkylated in 50 mM chloroacetamide for 30 min at RT. Samples were then diluted to 2 M urea with 10 mM Tris HCl (pH 7.4), and digested in 10 µg/ml Trypsin Gold (Mass Spectrometry Grade, Promega, V5280) overnight at RT. Excess trypsin was quenched by the addition of 1% (v/v) trifluoroacetic acid (TFA, Pierce, 28904) on ice for 15 min and centrifuged at 1,800 × *g* for 15 min at 4**°**C. The supernatants were adsorbed to C18 pipette tips (Bond Elut OMIX C18, Agilent, A7003100), and eluted in 50% acetonitrile (ACN, Optima^TM^ LC/MS Grade, Fisher Chemical, A955-4), 0.1% formic acid (FA, LC-MS Grade, Pierce, 28905). Eluted peptides were vacuum dried, reconstituted in 40 µl 5% ACN, 0.05% TFA, and analyzed by nLC-MS/MS.

### Treatment of TgSkp1 peptides with α-galactosidase

Trypsinates from above were centrifuged at 1800 × *g* for 15 min at 4**°**C. The supernatants were dried under vacuum, resuspended in 100 mM sodium citrate phosphate buffer (pH 6.0), and treated with 3.6 mU of green coffee-bean α-galactosidase (CalBiochem) for 18 h at 37**°**C. An additional 3.6 mU of α-galactosidase was added for an 8 h. After treatment, peptides were processed as above.

### Mass spectrometry of TgSkp1 peptides

Reconstituted peptides were loaded onto an Acclaim PepMap C18 trap column (300 μm, 100 Å) in 2% (v/v) ACN, 0.05% (v/v) TFA at 5 μl/min, eluted onto and from an Acclaim PepMap RSLC C18 column (75 μm × 150 mm, 2 μm, 100 Å) with a linear gradient consisting of 4-90% solvent B (solvent A: 0.1% FA; solvent B: 90% ACN, 0.08% (v/v) FA) over 180 min, at a flow rate of 300 nl/min with an Ultimate 3000 RSLCnano UHPLC system, into the ion source of an Orbitrap QE+ mass spectrometer (Thermo Fisher Scientific). The spray voltage was set to 1.9 kV and the heated capillary temperature was set to 280°C. Full MS scans were acquired from *m/z* 350 to 2000 at 70k resolution, and MS^2^ scans following higher energy collision-induced dissociation (HCD, 30) were collected for the Top10 most intense ions, with a 30-sec dynamic exclusion. The acquired raw spectra were analyzed using Sequest HT (Proteome Discoverer 2.2, Thermo Fisher Scientific) with a full MS peptide tolerance of 10 ppm and MS^2^ peptide fragment tolerance of 0.02 Da, and filtered to generate a 1% target decoy peptide-spectrum match (PSM) false discovery rate for protein assignments. All known glycoforms for TgSkp1 specific glycopeptides were manually searched for and verified.

### Enzyme assays

#### Sugar nucleotide hydrolysis

Recombinant TgGat1 (0.625-2.5 µM) was incubated with a given UDP-sugar (50 µM) (Promega) in 50 mM HEPES-NaOH (pH 7.4), 2 mM MnCl_2_, 5 mM DTT, in a final volume of 20 µl, for 1-16 h at 37°C. The UDP generated was quantitated using the UDP-Glo assay (Promega) as described (Sheikh et al., 2017).

#### Glycosyltransferase activity toward small glycosides

In the standard reaction, TgGat1 or PuGat1 was incubated with 2 mM synthetic glycosides [pNP-*α*-galactoside (pNP-*α*Gal), pNP-*β*-galactoside (pNP-*β*-Gal), pNP-*α*-glucoside (pNP-*α*Glc), pNP-*β*-glucoside (pNP-*β*Glc), pNP-*α*-maltoside, (pNP-malt), chloro-4-nitrophenyl-*α*-maltotrioside (ClpNP-trimalt), pNP-Skp1 trisaccharide (FGaGn-pNP), pNP-Skp1 tetrasaccharide (GlFGaGn-pNP)], 8 µM UDP-Gal (unlabeled), 0.17 µM UDP-[^3^H]Gal (15.6 µCi/nmol, American Radiolabeled Chemicals), 50 mM HEPES-NaOH (pH 7.0), 2 mM MnCl_2_, 5 mM DTT, in a final volume of 30 µl, for 1 h at 37°C. Pilot studies indicated a pH optimum of 7.0, with 50% activity at pH 8.0 and 75% activity at pH 6.0 for TgGat1. Salt dependence studies showed maximal activity with no added NaCl or KCl, and 35% activity at 800 mM of either salt. Activity showed a ∼6-fold preference for MnCl_2_ over MgCl_2_, with activity maximal at 2 mM MnCl_2_. The enzyme was essentially inactive in NiCl_2_, CoCl_2_ and CaCl_2_ (Fig. S9A-D). For kinetic studies, concentrations and times were varied as indicated, and kinetic parameters were analyzed according to the Michaelis-Menten model using Graph pad Prism software. Reactions were stopped by addition of 1 ml 1 mM ice-cold Na-EDTA (pH 8.0), and incorporation of radioactivity into pNP-glycosides was analyzed by capture and release from a Sep-Pak C18 cartridge and scintillation counting (Rahman et al., 2017).

#### Glycosyltransferase activity toward GlFGaGn-Skp1

To prepare Tg-GlFGaGn-Skp1, 2 nmols (40 µg) of recombinant TgSkp1 FGaGn-Skp1 (Rahman et. al 2017) was incubated with 1.3 nmols (88 μg) of Tg-His_6_-Glt1, 4 nmols UDP-Glc, 1.2 units alkaline phosphatase (Promega), 50 mM HEPES-NaOH (pH 8.0), 5 mM DTT, 2 mM MnCl_2_, 2 mM MgCl_2_, 2 mg/ml BSA in a final volume of 121 μl for 3.5 h at 37°C. The reaction was initiated by addition of UDP-Glc and terminated by freezing at −80°C. Reaction progress was monitored by Western blotting with pAb UOK104, which is specific for FGGn-Skp1 from either *D. discoideum* or *T. gondii* (Chinoy et al., 2015), followed by probing with pAb UOK75, which is pan-specific for all TgSkp1 isoforms, for normalization. Approximately 85% of total Skp1 was modified. GlFGaGn-Skp1 was purified from Glt1 on a mini-QAE-column using a Pharmacia Biotech SMART System. ∼19µg of GlFGaGn-Skp1 was applied to a mini QAE column pre-equilibrated with 50 mM Tris-HCl (pH 7.8), 5mM MgCl_2_, 0.1 mM EDTA (buffer A) and eluted with a gradient from 0% A to 100% buffer B (50 mM Tris-HCl (pH 7.8), 5 mM MgCl_2_, 0.1mM EDTA, 300 mM NaCl) in 40 min at a flow rate of 240 µl/min. GlFGaGn-Skp1fractions were identified based on A280 and Western blot probing with pAb UOK75.

TgGat1 (0.127 µM) was incubated with GlFGaGn-Skp1 or FGaGn-Skp1 (0.9-3.6 µM), 3.2 µM UDP-[^3^H]Gal (15.6 µCi/nmol), 40 µM UDP-Gal (unlabeled), 0.2% (v/v) Tween-20, 50 mM HEPES-NaOH (pH 7.0), 2 mM MnCl_2_, 5 mM DTT, in a final volume of 20 µl, for 1 h at 37°C. The reaction was stopped by addition of 4×-Laemmli sample buffer, 1 M DTT (50 mM final concentration), and 2 µg soybean trypsin inhibitor, and boiled for 3 min. SDS-PAGE and incorporation of radioactivity into Skp1 was performed as described above.

To detect Gat1 activity in cells, a cytosolic extract was prepared by hypotonic lysis, ultracentrifugation at 100,000 *g* **×** 1 h, and desalted as previously described (Rahman et al., 2016). 25 µg of desalted S100 protein was incubated with 10-50 nmol Tg-GlFGaGn-Skp1 and 1.0 µCi UDP-[^3^H]Gal (15.6 µCi/nmol) for 5 h, and incorporation of radioactivity into protein was assayed by SDS-PAGE and scintillation counting as described above.

#### Glycosyltransferase activity toward parasite extracts

To search for Gat1 substrates, cytosolic S100 fractions (180 µg protein) were incubated with 0.13 µM TgGat1, 2.0 µCi UDP-[^3^H]Gal (15.6 µCi/nmol) in a final volume of 60 µl containing 50 mM HEPES-NaOH (pH 7.0), 2 mM MnCl_2_, 5 mM DTT, 30 mM NaF, 0.2% Tween-20, at 37°C for 1 h, supplemented with 1.7 µg Tg-GlFGaGN-Skp1 as indicated. Incorporation of radioactivity was monitored by the SDS-PAGE assay as described above.

### Mass spectrometry of Skp1

To evaluate its glycosylation status, recombinant PuGat1 or TgGat1 (purified by gel filtration) were incubated with or without UDP-Gal or UDP-Glc in the absence of added acceptor substrate, and diluted to 50 ng/μl Skp1 with 2% acetonitrile, 0.05% (v/v) trifluoroacetic acid. 250-500 ng of protein (5-10 μl) was injected into an Acclaim PepMap C4 trap cartridge (300 μm × 5 mm) equilibrated with 0.05% trifluoroacetic acid, 2% acetonitrile, ramped up with an increasing gradient to 0.1% formic acid, 25% acetonitrile, and introduced into an Acclaim PepMap analytical C4 column (75 μm × 15 cm, 5 μm pore size) maintained at 35°C in an Ultimate 3000 RSLC system coupled to a QE+ Orbitrap mass spectrometer (Thermo Scientific). After equilibrating the analytical column in 98% LC-MS Buffer A (water, 0.1% formic acid) for 10 min and 6-min ramp up to 27% LC-MS Buffer B [90% (v/v) acetonitrile, 0.1% formic acid], separation was achieved using a linear gradient from 27% to 98% Buffer B over 20 min at a flow rate of 300 nl/min. The column was regenerated after each run by maintaining it at 98% Buffer B for 5 min. The effluent was introduced into the mass spectrometer by nanospray ionization in positive ion mode via a stainless-steel emitter with spray voltage set to 1.9 k, capillary temperature set at 250°C and probe heater temperature set at 350°C. The MS method consisted of collecting Full ITMS (MS^1^) scans (400-2000 *m*/*z*) at 140,000 resolution in intact protein mode (default gas P set to 0.2). PuGat1 species eluting between 17.5 and 21.5 min and TgGat1 species eluting between 18.5 and 22.5 min (∼60 to 80% acetonitrile) were processed with Xcalibur Xtract deconvolution software to generate monoisotopic masses from the multicharged, protonated ion series. Since TgGat1 MS spectra were not isotopically resolved, masses were extracted after MS spectra deconvolution using the ReSpect algorithm in the BioPharma Finder suite (Thermo Scientific), with a 20 ppm deconvolution mass tolerance and 25 ppm protein sequence matching mass tolerance. For consistency, PuGat1 was also deconvoluted and re-extracted using the ReSpect algorithm and the same conditions.

### Structure determination of PuGat1

A PuGat1:UDP:Mn^2+^ complex in 50 mM HEPES-NaOH, pH 7.4, 75 mM NaCl, 2 mM DTT, 5 mM UDP, and 5 mM MnCl_2_ was crystallized at 20°C using a hanging drop vapor diffusion method over a reservoir containing 8-12% (w/v) PEG4000, 0.4 M ammonium sulfate, and 0.1 M sodium acetate at pH 4.0. Crystals were obtained overnight and were transferred to a reservoir solution containing 15% (v/v) of a cryoprotectant mixture (1:1:1 ethylene glycol:dimethyl sulfoxide:glycerol), and flash cooled with liquid N_2_. The complex crystallized in space group P4_2_2_1_2 and diffracted to 1.76 Å (Table S3). X-ray data were collected remotely at the SER-CAT 22-BM beamline at the Argonne National Laboratory using a Fast Rayonix 300HS detector, and processed using XDS (Kabsch 2010), with 5% of the data omitted for cross validation.

PuGat1:UDP:Mn^2+^ crystals were soaked with platinum cyanide for heavy-atom phasing, cryoprotected, and frozen as above. PuGat1:UDP:Pt^2+^ crystals were isomorphous to PuGat1:UDP:Mn^2+^ crystals and diffracted to 2.1 Å (Table S3). PuGat1:UDP:Mn^2+^ crystals were alternatively soaked in UO_2_ (not shown). The crystals, which diffracted to 2.4 Å in the same space group, lacked UDP density, suggesting displacement by UO_2_.

The crystal structure of PuGat1:Pt^2+^ was solved using single-wavelength anomalous dispersion (SAD). The data was obtained at a wavelength of 1.85 Å for maximum anomalous signal. A single Pt^2+^ site was located using PHENIX (Adams et al., 2010), and the resulting phases had an acceptable figure of merit of 0.31. The model was subjected to iterative cycles of refinement and yielded a final model with R_work_/R_free_ of 0.21/0.24 (Table S3). The structure of PuGat1:UDP:Mn^2+^ was solved using rigid body refinement of PuGat1:Pt^2+^. The resulting model was subjected to iterative cycles of refinement and yielded a final model with R_work_/R_free_ of 0.18/0.21 (Table S3). Images were rendered in PyMol (Delano 2002).

### Glycan docking

The lowest energy conformation of the TgSkp1 tetrasaccharide (Glcα1,3Fucα1,2Galβ1,3GlcNAcα1-OH) was generated via GLYCAM (Woods, 2014). Hydrogen atoms were added, and the electrostatic surface was generated using AutoDockTools (Morris et al., 2009). A grid box with dimensions 26 Å × 26 Å × 34 Å was placed over the ligand binding site based on where the acceptor is bound on glycogenin/glucan complex. The ligand was kept rigid, since AutoDock Vina is not parameterized specific to glycan torsion angles. 100 binding modes were calculated with the lowest binding energy scored at −4.4 kcal/mol and the highest binding energy scored at −5.7 kcal/mol.

### Sedimentation velocity

PuGat1 was further purified on a Superdex S200 gel filtration column (GE Healthcare) equilibrated with 20 mM potassium phosphate (pH 7.4), 50 mM KCl, 0.5 mM TCEP. Protein concentration was calculated from *A*_280_ measured in an Agilent 8453 UV/Vis spectrophotometer, based on a molar absorptivity (ε_280_) of 60390 M^-1^cm^-1^, which was calculated from the PuGat1 sequence using ProtParam (Gastiger et al., 2005). Samples were diluted to 0.3-11 µM, loaded into 12 mm double-sector Epon centerpieces equipped with quartz windows, and equilibrated for 2 h at 20°C in an An60 Ti rotor. Sedimentation velocity data were collected using an Optima XLA analytical ultracentrifuge (Beckman Coulter) at a rotor speed of 50000 RPM at 20°C. Data were recorded at 280 nm for protein samples at 3.5-11 µM, and at 230/220 nm for samples at 0.3-1.5 μM, in radial step sizes of 0.003 cm. SEDNTERP (Laue et al., 1992) was used to model the partial specific volume of PuGat1 (0.73818 mL/g), and the density (1.0034 g/ml) and viscosity (0.0100757 P) of the buffer. Using SEDFIT (Schuck, 2000), data were modeled as continuous *c*(*s*) distributions and were fit using baseline, meniscus, frictional coefficient, and systematic time-invariant and radial-invariant noise. Predicted sedimentation coefficient (s) values for the PuGat1 monomer (2.8 S) and dimer (4.2 S) were calculated using HYDROPRO (Ortega et al., 2011). Data fit and *c*(*s*) plots were generated using GUSSI (Brautigam, 2015).

### Molecular dynamics simulations

The model for TgSkp1 was built as described previously for DdSkp1 (Sheikh et al., 2017). Briefly, a homology model of TgSkp1 was generated with the SWISS-MODEL web server (Arnold et al., 2006) based on the human Skp1 template from PDB ID: 2ASS (Hao et al., 2005), and missing residues were appended with UCSF Chimera (Petterson et al., 2004). Molecular dynamics simulations were performed as described previously. Briefly, MD simulations were performed with the pmemd.cuda version of AMBER14 (Götz et al., 2012). The amino acid and carbohydrate residues were parameterized with the FF12SB and GLYCAM06 (J-1) force fields, respectively (Case et al., 2005; Kirschner et al., 2008). The systems were neutralized with Na+ ions and solvated using the TIP3P water model (Jorgensen et al., 1983) in a truncated octahedral box with 15-Å distance from the solute to the end of the unit cell. Electrostatic interactions were treated with the particle mesh–Ewald algorithm, and a cut-off for non-bonded interactions was set to 8 Å (Darden et al., 1993). SHAKE was employed to constrain hydrogen-containing bonds, enabling an integration time step of 2 fs. Restraints were imposed in specific situations and were enforced with a 10-kcal/mol Å^2^ energy barrier in each case. Each minimization step consisted of 1000 cycles of the steepest descent method (1000 cycles), followed by 24,000 cycles using the conjugate gradient approach. The systems were heated to 300 °K under NVT conditions over 60 ps, employing the Berendsen thermostat with a coupling time constant of 1 ps. The subsequent simulations were performed under NPT conditions. A torsion term that corrects 4(*trans*)-hydroxyproline residue (Hyp) ring puckering was included in simulations of the O-linked residue type (OLP) based on previous studies that indicate that the ring is primarily exo when glycosylated (Park et al., 2005; Owens et al., 2007). This torsion term has been adopted in GLYCAM06 (version K).

A 50-ns simulation of the protein was performed with Cα cartesian constraints on all amino acids except those generated by Chimera. The fully glycosylated isoform was created by adding the TgSkp1 pentasaccharide to the exo-pucker conformation of hydroxyproline (residue 154). Six independent simulations were performed. Three ran for 250 ns directly, while the other three began with an additional 50 ns in which the protein was restrained to allow the glycan time to adapt to the protein conformation.

### Computational Analysis

Structural images were created with Visual Molecular Dynamics (Humphrey et al., 1996) and the 3D-SNFG plugin (Thieker et al., 2016). The structure depicted in Fig. 6 was created by identifying the frame from equil-1 that consisted of the lowest RMSD to the average structure as calculated by cpptraj (Roe et al., 2013). The cpptraj program was also used to distribute the latter 200 ns of the six simulations into 48 bins containing 250 frames each for analysis. Per-residue MMGBSA energies were calculated with MMPBSA.py.MPI with igb=2 and idecomp=3. A bash script was used to calculate the correlation coefficients (Miller et al., 2012).

### Phylogenetic analysis of enzyme sequences

Proteins related to TgGat1 were searched for using a BLASTP **(V 2.4.0)** search seeded with the full-length TgGat1 protein sequence against the NCBI non-redundant database (December 2016). The evolutionary relationship of Gat1-like sequences was investigated by using a Maximum Likelihood method (Le and Gascuel, 2008) and conducted in MEGA7 (Kumar et al., 2016). Catalytic domains from 43 CAZy GT8 sequences selected based on their relatedness to Gat1, glycogenin, or known function, and consisted of 196 positions. Sequence alignments were manually-curated in BioEdit (v 7.2.5). Initial tree(s) for the heuristic search were obtained automatically by applying Neighbor-Join and BioNJ algorithms to a matrix of pairwise distances estimated using a JTT model, and then selecting the topology with superior log likelihood value. A discrete Gamma distribution was used to model evolutionary rate differences among sites (5 categories (+*G*, parameter = 1.1608)). The rate variation model allowed for some sites to be evolutionarily invariable ([+*I*], 1.02% sites).

## Supporting information

Full Supplement

## ACKNOWLEDGMENTS

We thank Kentuan Hicks and Nathan Beattie for their assistance. MM was partially supported by NIH T32-AI060546; HWK was partially supported by NIH T32-GM107004; and KH was supported by NSF REU grant DBI-1426834. This project was supported in part by NIH RO1-GM084383 to CMW and Ira Blader, Grant #14-140 from the Mizutani Foundation for Glycoscience to CMW and Ira Blader, NIH P41-GM103490 (to LW, senior investigator), NIH 8P41-GM103390 (Resource for Integrated Glycotechnology to J. Prestegard), NIH R01-GM114298 to ZAW, and NIH P01-GM107012 to John Rose.

## ABBREVIATIONS

DBA: *Dolichos biflorus* lectin
Dd: *Dictyostelium discoideum*
FBP: F-box protein
GaGlFGaGn–: Galα1,3Glcα1,3Fucα1,2Galβ1,3GlcNAcα1–
GalT: galactosyl transferase
GT: glycosyltransferase
Hyp: (*2S*,*4R*)-4-hydroxy-l-proline
mAb: monoclonal antibody
P4H: prolyl 4-hydroxylase
pAb: polyclonal antibody
pNP: para-nitrophenol
Pu: *Pythium ultimum*
RHΔΔ: RHΔ*ku80*Δ*hxgprt*
SCF: Skp1/Cullin-1/F-box protein subcomplex of E3 Cullin-RING-1 ubiquitin ligases
Tg: *Toxoplasma gondii*
Ub: ubiquitin

## AUTHOR CONTRIBUTIONS

Msano Mandalasi, Investigation, Methodology, Writing–review and editing; Hyun W. Kim, Investigation, Methodology, Data curation, Writing–review and editing; Kazi Rahman, Investigation, Methodology; M. O. Sheikh, Methodology, Writing–review and editing; David Thieker, Methodology, Writing–review and editing; Elisabet Gas-Pascual, Methodology, Data curation; Peng Zhao, Methodology, Data curation; Nitin Daniel, Methodology, Data curation; Hanke van der Wel, Methodology; Travis H. Ichikawa, Methodology; John N. Glushka, Methodology, Data Curation; Lance Wells, Investigation, Methodology; Robert J. Woods, Conceptualization, Supervision, Editing; Zachary A. Wood, Conceptualization, Supervision, Writing–review, editing; Christopher M. West, Conceptualization, Supervision, Funding acquisition, Writing–original draft, Project administration

## SUPPLEMENT

The Supplement contains extended Experimental Methods, Figures S1-S14, and Tables S1-S3.

