## Supplementary material for "A glycogenin homolog controls *Toxoplasma gondii* growth via glycosylation of an E3 ubiquitin ligase": Full Supplement

**Table of Contents**

Table S1. Oligonucleotides employed
Table S2. Skp1 glycopeptide mass measurements (supports Fig. 1C)
Table S3. PuGat1 crystal parameters

Fig. S1. Nucleotide and amino acid sequences of TgGat1
Fig. S2. Nucleotide and amino acid sequences of PuGat1
Fig. S3 A-C. Disruption and complementation of Tg*gat1* in RHΔΔ
Fig. S4 A-B. Disruption and complementation of Tg*gat1* in Ku80+ strains
Fig. S5 A-G. nLC/MS of Skp1 glycopeptides (supports Fig. 1C, Table S2)
Fig. S6. Comparison of Gat1-like and glycogenin sequences
Fig. S7. Summary of Gat1-related sequences selected for phylogenetic analysis
Fig. S8. Alignment of Gat1-like, glycogenin-like, and other CAZy GT8 sequences (supports phylogenetic tree in Fig. 2)
Fig. S9 A-I. Characterization of the αGalT activity of Gat1 and biochemical complementation of *Toxoplasma* extracts (supports Fig. 4)
Fig. S10 A-E. Absence of Gat1 autoglycosylation
Fig. S11 A-C. NMR analysis of the *Tg* Skp1 pentasaccharide
Fig. S12 A-E. Computational comparison of the Skp1 glycans from *Toxoplasma* and *Dictyostelium*
Fig. S13. PuGat1 and Oc-glycogenin-1 ligand interactions
Fig. S14. A, B. Sedimentation velocity analyses of PuGat1

#### Table S1. List of primers

| Purpose | Code name | Primer name | Primer sequence | Location |
|---|---|---|---|---|
| *gat1* disruption in RHΔΔ and complementation in RH | Fa | a) Gat1F1 5'-flank 5'-end 5' | GGGGGCCCAACCAGCGGATCTTCTGAAC (ApaI) | Tg*gat1* homologous recombination disruption & complementation plasmids |
| | Ra | a') Gat1R1 5'-flank 3'-end 5 | GGCTCGAGACGCGTTGAGCGATTGA (Xho1) | |
| | Fb | b) Gat1F2 3'-flank 5'-end 5' | GCTCTAGAGAGGGAGAACCAAAGTGATGAT (XbaI) | |
| | Rb | b') Gat1R2 3'-flank 3'-end 5 | CGCGGCCGCTGCGTAGAACACAAGGAGAAC (NotI) | |
| PCR confirmation for *gat1* disruption in RHΔΔ | | | | |
| PCR1 | Fc | Forward | TACCCTGTTGACGGACAATT | Tg*gat1* genomic sequence |
| | Rc | Reverse | CTTTGCTGGTTGTTCCCAAG | |
| PCR2 | Fd | Forward | GAACCGAATGACAACGCATTAC | |
| | Rd | Reverse | AGTCGCGGAACATCTCGTTGAAGT | HXGPRT sequence |
| PCR3 | Fe | Forward | ATTTGCATCCTGAAAGGCTCTCGC | |
| | Re | Reverse | TCTGAAATGGAGTCGCCTTG | Tg*gat1* genomic sequence |
| Dual guide CRISPR plasmid for *gat1* disruption in RH, Me49-RFP | | | | |
| NsiI PCR | Ff | Plasmid 3 FOR | CGTGGGGATGCATTCACCGCGGCCACATGTTG | Dual guide *gat1* CRISPR disruption plasmid |
| | Rf | Plasmid 3 REV | GCGATGAGCGCAAGCCGTCCTGAGTTACG | |
| dg plasmid sequencing | Fg | gRNA FOR | CAAAGTGCGCGAGTTGAAATCG | |
| | Rg | gRNA REV | GAGACGATGATTCCTGATCACTCCG | |
| PCR confirmation for *gat1* disruption in RH and Me49-RFP | Fh | Gat1 63 seq Fw (P1) | CGTACGCTACCCTGTTGACG | Tg*gat1* genomic sequence |
| | Rh | Gat1 968 Seq Rv (P2) | AGAATCAGTTGGCACAGTGCC | |
| | Fi | DHFR F/R Fw (P4) | CCATTGCGGTGTCGTGGATT | DHFR sequence |
| | Ri | DHFR RO Rv (P3) | CCCCTGTGTCCTTTATCGAAG | |
| Complementation plasmid sequencing | Fl | TgGat1 seq Fw | GGACTGTTTCACCAAACTGCGTGTGTG | Tg*gat1* genomic sequence |
| | Rl | Gat1 3'UTR Rv | CTAGTCAGTCCCTAAGGCTAGT | |
| Ty tag insertion on complementation plasmids | | | | |
| Tg*gat1* Ty Tag insertion | Fn | TgGat1-Ty Fw | GAAGTACACACAAACCAAGACCCACTAGACTAGTGGAGGGAGA | Tg*gat1* genomic sequence & Ty tag |
| | Rn | TgGat1-Ty Rv | GTTTGTGTGTACTTCCACGATATCAGAATCAGTTGGC | |
| PCR confirmation for *gat1* complementation in RH | Fp | UPRT Fw | GTCCCAACGTCGCAAGTAA | UPRT genomic sequence |
| | Rp | UPRT Rv | ATGCGGACTTTCGGGTATTC | |
| | Fq | Gat1 check Fw | TGGGAACAACCAGCAAAGA | Tg*gat1* genomic sequence |
| | Rq | Gat1 FO Rv | GGGGTTGCAGCCTATGG | |
| TgGat1 *E. coli* expression plasmid | Fr | Gat1 Fw | AAGCTAGCATGTCTCCTCGGTACGCGTACGCT | Tg*gat1* genomic sequence & pET15b expression plasmid |
| | Rr | Gat1 Rv | AAGGATCCCTACACGATATCAGAATCAGTTGGCACAG | |
| DHFR amplicon with 45 bp *gat1* arms for CRISPR disruption in RHΔΔ | Fs | 63 Fw_dhfr Fw | CGGACAATTCTTTCTACTATGGTGTCGAGGCACTGCTCAAGTCACAAGCTTCGCCAGGCTGTAAAT | Tg*gat1* and DHFR |
| | Rs | 968 Rv_dhfr Rv | ATCAGAATCAGTTGGCACAGTGCCCGTAAGGAAGACTTTCCACCACATCCTGCAAAGTGCATAGAAG | |
| Gat1-HA complementation plasmid in RHΔΔ | Ft | 3HA Fw | GGTACCTACCCGTACGACGTC | pUPRT a1 WT cDNA shuttle Vector-Tub1-3xHA |
| | Rt | Tub-5'UTR Rv | GGCGCGCCGTGTCGAAAA | |
| | Fu | Tub-5'UTR-Gat1 Fw | CTTTTTCGACACGGCGCGCCATGTCTCCTCGGTACGCG | pmini-Tg*gat1* plasmid |
| | Ru | HA-Gat1 Rv | ACGTCGTACGGGTAGGTACCCACGATATCAGAATCAGTTGGC | |
| PCR confirmation for Gat1 complementation in RH | Fv | UPRT 5'Arm Fw | GCTGTGCCTAGTATCGAAAGCTGTA | UPRT genomic sequence |
| | Rv | Gat1 at STOP Rv | CTACACGATATCAGAATCAGTTGGCACA | Tg*gat1* genomic sequence |
| | Fl | TgGat1 seq Fw | GGACTGTTTCACCAAACTGCGTGTGTG | |
| | Rw | UPRT 3'Arm Rv | CGACGTCACTGTACGACATCC | UPRT genomic sequence |

#### Table S2. Skp1 glycopeptide mass measurements (supports Fig. 1C)

Isoforms of the Skp1 peptide 145-IFNIVNDFTPEEEAQVR were detected and quantified as described in Material and Methods.

The abundances of raw ion counts for the detected isoforms are shown for all the strains analyzed.

| Strain* | unmodified peptide | | | H-dH-H-HN-O-peptide[b] | | | H-H-dH-H-HN-O-peptide | | | All peptides |
|---|---|---|---|---|---|---|---|---|---|---|
| | abundance[a] | $[M+2H]^{2+}$<br>$[M+3H]^{3+}$ | Δm/z[c]<br>Δm/z | abundance | $[M+2H]^{2+}$<br>$[M+3H]^{3+}$ | Δm/z<br>Δm/z | abundance | $[M+2H]^{2+}$<br>$[M+3H]^{3+}$ | Δm/z<br>Δm/z | total abundance |
| RH, wt | 1.12E+07 | 1011.002 | 0.40 | nd[d] | | | 3.96E+04 | 1436.650 | 0.28 | 2.13E+07 |
| | 8.59E+06 | 674.337 | 0.56 | | | | 1.40E+06 | 958.103 | -0.30 | |
| RH + aGal*ase* | 4.71E+06 | 1011.000 | 2.37 | 1.07E+05 | 1355.62 | 3.61 | nd | | | 7.00E+06 |
| | 1.34E+06 | 674.335 | 3.53 | 8.41E+05 | 904.081 | 4.47 | | | | |
| Δ*gat1*/RH MM12.A8 | 9.32E+06 | 1011.001 | 1.38 | 9.99E+04 | 1355.62 | 1.40 | nd | | | 1.65E+07 |
| | 5.67E+06 | 674.336 | 2.05 | 1.46E+06 | 904.083 | 2.26 | | | | |
| *gat1*::gat1-ty/ *gat1*Δ/RH MM21.E12 | 1.42E+07 | 1011.000 | 2.37 | nd | | | 5.77E+04 | 1436.65 | 2.37 | 2.21E+07 |
| | 6.08E+06 | 674.336 | 2.05 | | | | 1.79E+06 | 958.101 | 1.78 | |
| Me49-RFP MM8.A10 | 1.81E+07 | 1011.000 | 2.37 | | | | 3.88E+04 | 1436.65 | 2.37 | 3.26E+07 |
| | 1.20E+07 | 674.336 | 2.05 | | | | 2.58E+06 | 958.100 | 2.83 | |
| Δ*gat1*/ME49 MM14.B5 | 8.27E+06 | 1011.000 | 2.37 | 4.57E+04 | 1355.61 | 10.99 | nd | | | 1.30E+07 |
| | 3.64E+06 | 674.336 | 2.05 | 1.08E+06 | 904.081 | 4.47 | | | | |

Notes:

* see Table 1 for other descriptions

Hydroxylated, mono, di and trisaccharide glycopeptides were not detected

[a] abundance from ion raw spectral counts

[b] H=Hex; dH=deoxyHex; HN=HexNAc

[c] Δm/z in ppm

[d] nd: not detected

Expected masses for each glycoform are as follows:

| unmodified peptide | |
|---|---|
| $[M+2H]^{2+}$ | 1011.002 |
| $[M+3H]^{3+}$ | 674.337 |

| tetrasaccharide-peptide | |
|---|---|
| $[M+2H]^{2+}$ | 1355.624 |
| $[M+3H]^{3+}$ | 904.085 |

| pentasaccharide-peptide | |
|---|---|
| $[M+2H]^{2+}$ | 1436.650 |
| $[M+3H]^{3+}$ | 958.103 |

**Table S3. Crystallographic data**

| **Data collection** | PuGat1:UDP:$Pt^{2+}$ (6MW5) | PuGat1:UDP:$Mn^{2+}$ (6MW8) |
|---|---|---|
| Wavelength (Å) | 1.85 | 1.0 |
| Space group | $P4_2 2_1 2$ | $P4_2 2_1 2$ |
| Unit cell dimensions (a, b, c) | 83.78, 83.78, 75.84, 90.00, 90.00, 90.00 | 84.06, 84.06, 76.08, 90.00, 90.00, 90.00 |
| Completeness (%) | 97.4 (94.8)[a] | 99.9 (99.8)[a] |
| Total number of reflections | 396057 (13045) | 800853 (52625) |
| Unique reflections | 29424 (2114) | 28125 (2042) |
| Redundancy | 13.5 (6.2) | 28.4 (25.8) |
| I/σ(I) | 28.6 (1.21) | 36.94 (1.85) |
| $R_{meas}$[b] (%) | 6.1 (145.4) | 6.2 (203.9) |
| CC1/2[c] (%) | 100.0 (49.0) | 100.0 (65.5) |
| **Refinement** | | |
| Resolution (Å) | 2.1 | 1.76 |
| $R_{work}/R_{free}$ | 0.196/0.242 | 0.181/0.210 |
| No. of atoms Protein/ Ligand / Water | 1957/30/54 | 1949/42/122 |
| Wilson B-factor (Å$^2$) | 45.2 | 39.9 |
| B-factors (Å$^2$) Protein/ Ligands and Water | 45.2/45.66 | 39.4/44.2 |
| **Stereochemical Ideality** | | |
| Bond lengths (Å) | 0.006 | 0.006 |
| Bond angles (°) | 0.786 | 0.784 |
| φ, ψ Most favored (%) | 97 | 99 |
| φ, ψ Additionally allowed (%) | 3 | 1 |
| **SAD Phasing statistics** | | |
| Heavy atom sites | 1 | |
| Figure of merit | 0.31 | |

[a] Values in parentheses are for highest-resolution shell

[b] $R_{meas}$ is the redundancy independent merging R-factor of Karplus and Diederichs (2012)

[c] $CC_{1/2}$ is the percentage of correlation between intensities from random half-data sets

**Figure S1.** Genomic sequence surrounding the open reading frame of Gat1 (TGME49_310400 model from Toxodb.org). Numbering begins at the A of the start codon ATG. Coding sequences, including those upstream and downstream of Gat1, are capitalized; non-coding sequences are lower case. Amino acid sequence of Gat1 is above its coding sequence. Sequences of oligonucleotides from Table S1 are shown and mapped. For forward PCR primers, cognate sequences are colored purple; cognate sequences of reverse PCR primers are in red; guide DNA sequences are in blue. nt differences observed in the type 1 RH strain are indicated.

```
>TGME49_chrXI:1317070..1322069
ATCGTCGCTATACATGAGGAAGCGAACGTCAGACAACCCAGGGCAAACGAAATTGGAAGA
AAGACACTTCCTAAACTCGTCGGCGGGGGTGCCAGTGAATCGATTGCGATTCGACTTGC
TCTTGAGTTGGATACTCCGGCCGGCCAATTTCGAACATTGTCATTATCCAGTCCATTGCT
AGGTAGAGTGTGGAGAAGAGAGAGAAAACCGCAATGCAAATCGGAACCCAAAAGGCTGCC
TGTCTACTTCCTGGCACGGCCATgatttccggaggaactgaggacgatctcctttttgcc TGGT1_310390 reverse
gtagataactttgccctgcgtcgtctcctctttgttccttctggccaccttgacctctct
tcttgctttaaattgaatctcgttcccgcacgtttctcctgtttcccttctttgcgttct
cccgtcactttgctgccgtctctcttttcggggtctttctccgtcgtaaacgggtccttg
tttcatcagccgcgacagttctgccttcgtcgtgttttcccccctctgttcttgtgtttcg
tgttgtctcactcgtcgtgtgtcactcctccctccgtcccgttcctgtttcctactaggt
tcccttccctctggctcatccaaaaaagggatctaaatgacagcgaacatcccgacaact
ttttctcagctaaaatcacgaactatgcccaacacacaaggcgcgtaccataattcccgt
ataggttccctcctacacggcaacacttaaacaggcgcgtttctaaactattacgaagtc
gaggcgagaaccccacaacttcctatccagtccacaagaaccgaatgacaacgcattact
                                  5′-gaaccgaatgacaacgcattac  Fd

tttacacaagtgctcagttgaacacactagatacacaaatttttgttgctgcatgttgag
tccaccagtgacaccacatcgaaggcgttgcgctacttttgacctcgttgagtgccctcg
                                                      5′-ggg Fa

cggaaaaccagcggatcttctgaacacgtcttctgcgaagaagaagcgtaagctaccact
ggcccaaccagcggatcttctgaac                                   Fa

tttcagtgacgcttattgaaagaacaaatgaaggtgacagaatcaaaaggaaacacagga
ccgcaaagccaacactcttccacattttcagcggaatttactcacgtaccatcgcctgac

       a                                                    RH
tgaggccgtaggctgcaaccccccctccaccgctcccgtcgtttctcagcctgacggcac
     ggtatccgacgttgggg-5′                                   Rq

tcagacggcgagtttcgggaagcagcctgccttttcaaacatatatcaatctactgcgtt
cccgttattattggcgcagaacgcgcaaaaagaagtcttcacatctacatccccgccacg
gctgtcgcttggaaaactgctctaaatgccaatgatggccattagtgcacacatgaaagc
cgatgttttctaagcgaataataccaaggggaactccgggttgtctttggaagacgaaaa
atccaatgcatgacttcgatgtaagtaaaagtgagctgaattgaaagcgagggaaagaga
cgcgaattctggcttgcttgacttccccccctccggccccccgcctccagcgaagcgaa
tgtgtcgcaggtgaccgtggaagccccattcgccaaccagcagcgtttcggagagtgaaa
tcttgattctcacaggagctagaattctgaagcttactcacagtgttgtagggacgcgtg
tgaagtgtcaggcattatttcactggtgggacgtggcggtgtttaataattcgttccggc
atgtgtgtgtctcaaatccccttttcttcgccggcacttcgaggcggaacgggaggggg
cacacgtttgctttgctgccaaagcagccgatttcgcgcccccctcaacggctcaaaggca
cttcaactgcacatgcatgcacagttcccgattcgtctgccggatcatgtgtgtgcatgt
ttgtatgttgtgggcatcattcacgttttccgtgtttatactggttcaagagcgtacatt
caggaagttgcactaaatggaaaattggcttcctgtgtggacacaggacacgggggttct
ctttgtctcccttcgctgtgacgcgttttgttgtgatcaagggcgctgtttaagcgtgat

                                                t           RH
gcattaagcggcacttcagtctgaatcaatcgctcaacgcgtcatttcctttcctttttt
                         agttagcgagttgcgcagagctcgg-5′       Ra
                                       gcagtaaaagaaaggaaaaaa Rk
                                              5′-cttttcgacac Fu
         M  S  P  R  Y  A  Y  A  T  L  L  T  D  N  S  F  Y     17
gctaaggagATGTCTCCTCGGTACGCGTACGCTACCCTGTTGACGGACAATTCTTTCTAC   51
cgattcctctactgacag-5′                                       Rk
 aagctagcatgtctcctcggtacgcgtacgct-3′                        Fr
```

```
 ggcgcgccatgtctcctcggtacgcg-3'                               Fu
                      5'-cgtacgctaccctgttgacg                Fh
                             5'-taccctgttgacggacaatt         Fc
                                        5'-CGGACAATTCTTTCTAC Fs

Y  G  V  E  A  L  L  K  S  L  E  A  T  K  T  P  Y  P  V  L     37
TATGGTGTCGAGGCACTGCTCAAGTCACTGGAGGCTACGAAGACGCCTTACCCCGTGCTT  111
        5'-ggcactgctcaagtcactgg                              gDNA-63
TATGGTGTCGAGGCACTGCTCAAGTCACAAGCTTCGCCAGGCTGTAAAT            Fs

L  L  H  T  S  D  V  S  Q  S  T  I  K  A  L  V  Y  Q  R  R     57
CTTTTGCACACATCTGATGTTTCTCAGAGTACAATAAAAGCGTTGGTTTATCAGCGTCGA  171

K  A  P  A  S  E  D  A  G  T  T  G  K  E  M  K  T  G  Q  E     77
AAAGCCCCGGCGAGTGAGGATGCGGGAACTACAGGGAAGGAAATGAAAACAGGGCAGGAA  231

V  I  P  S  S  Q  C  P  E  H  T  P  G  R  N  L  H  S  P  I     97
GTCATCCCAAGTTCACAGTGTCCAGAACACACCCCAGGTAGAAACTTGCACTCCCCCATT  291

G  R  K  G  V  N  P  V  S  C  S  V  T  Q  D  E  T  R  V  R    117
GGCAGGAAAGGGGTAAACCCTGTGAGTTGCTCCGTCACACAAGACGAGACTAGGGTTCGT  351

T  D  S  D  R  I  E  E  A  E  R  R  A  S  E  R  T  S  E  R    137
ACTGATTCAGATCGTATAGAAGAAGCAGAGCGTCGAGCCTCAGAGAGAACCTCGGAGCGA  411

A  R  A  G  E  T  E  E  Q  G  I  C  V  I  P  R  L  V  G  S    157
GCGAGAGCTGGGGAAACGGAGGAACAGGGCATTTGCGTTATTCCCCGACTCGTTGGTTCT  471

V  A  Y  P  K  A  E  R  D  T  C  P  V  E  G  W  K  D  C  F    177
GTCGCGTACCCTAAAGCGGAACGGGACACGTGCCCTGTTGAAGGGTGGAAGGACTGTTTC  531
                                               5'-ggactgtttc Fl

T  K  L  R  V  W  E  Q  V  D  F  D  V  I  V  Y  V  D  A  D    197
ACCAAACTGCGTGTGTGGGAGCAGGTTGACTTCGATGTGATTGTGTATGTCGACGCGGAC  591
accaaactgcgtgtgtg                                            Fl

C  I  V  L  R  P  V  D  E  L  F  L  R  Q  P  L  P  A  F  A    217
TGTATAGTTTTGCGGCCGGTAGACGAGCTTTTTCTTAGGCAGCCACTACCCGCCTTTGCA  651

P  D  I  F  P  P  D  K  F  N  A  G  V  A  V  L  K  P  D  L    237
CCAGATATCTTCCCTCCCGATAAATTTAACGCGGGAGTCGCAGTGCTGAAGCCCGACCTC  711

G  E  Y  G  N  M  V  A  A  V  E  R  L  P  S  Y  D  G  G  D    257
GGCGAATACGGAAATATGGTAGCCGCGGTCGAGCGTTTACCTTCATATGACGGAGGCGAC  771

T  G  F  L  N  A  Y  F  S  S  W  Y  E  N  A  A  G  A  R  L    277
ACAGGGTTTTTGAACGCGTATTTCTCATCGTGGTATGAAAACGCAGCTGGCGCCCGTTTG  831

P  F  R  Y  N  A  L  R  T  L  Y  H  M  T  Y  S  S  R  K  G    297
CCCTTTCGGTACAATGCTCTGCGCACACTGTATCACATGACGTACTCCAGTCGAAAAGGA  891

Y  W  N  A  V  K  P  I  K  I  L  H  F  C  S  S  P  K  P  W    317
TACTGGAATGCCGTCAAGCCGATCAAAATCCTGCACTTCTGCTCCTCCCCGAAGCCTTGG  951
                                                       gaacc Rc
                                                      5'-tgg Fq

E  Q  P  A  K  T  D  L  E  E  L  W  W  K  V  F  L  T  G  T    337
GAACAACCAGCAAAGACCGACCTCGAGGAACTATGGTGGAAAGTCTTCCTTACGGGCACT 1011
gaacaaccagcaaaga                                             Fq
cttgttggtcgtttc-5'                                           Rc
             5'-ccgacctcgaggaactatgg                         gDNA-968
                                                           a Rv
cacggttgactaagactatagcacc-5'                                 Rv
```

```
                                                      ccgtga Rh
                                                   5'-gaagta Fn
                                                          ct Rr
           gaagatacgtgaaacgtcctacaccacctttcagaaggaatgcccgtga Rs

V  P  T  D  S  D  I  V  *                                     345
GTGCCAACTGATTCTGATATCGTGTAGtggagggagaaccaaagtgatgatgaaagaatg 1071
cacggttgactaaga-5'                                           Rh
cacacaaaccaagacccactagactagtggagggaga                        Fn
cacggttgactaagactatagcacatcggatcctt-5'                       Rr
  gccttgactaagactatagcaccttcatgtgtgtttg-5'                   Rn
  cggttgactaagactatagcacccatggatgggcatgctgca-5'              Ru
               5'-cttggatagtggagggagaaccaaagtgatga           Fk
                    5'-gctctagagggagaaccaaagtgatgat          Fb
cacggttgactaagacta-5'                                        Rs

accgacttccaaaagaacggaaacgccggacagctgcctcgcggtaccttgggaaaagag
cgggacgtgtggaatcctgtcaactatctctttctgtgtcacctgtggacgaattgtaaa
tcttgtaaagtacaaacggagtacagcttaatcttgtaattttctttcttcgaaggacgc

                                                     a       RH
cagtgcgccgcaagcgtctagtggcctgcaaagactagccttagggactgactggttcgc
                                 tgatcggaatccctgactgatc-5'   Rl

gtacgcaaccatcacgcacaagcatgtttatttgccactggtgtgtcactcagctagacg
cgtcatgtttatgtatacgtacgtttcacagcctctcagagacatcccgcacaacgcatg
aaccgctgcaaccagaatactgaccgtcagcggtttcgcttgctttaaatcgggttgttt
tggaaaaactcaaaggtagtcgttctgtacatctccctaagttaagcggtaagttactcg
acgagcatacattgacaataagacggttctcacaatgaacatcccaaagagggcactaga
ccaaacaaaggagctaaaagacacgagcaagatgaagataaaacgcaccttagcgaaggc
catataacaaagtggatcttcacagtatcattctgtgtccgtaccagtcgctgcgaacaa
gaagacgcatgtgaacggttCTACTCTCGGAATTGAAAGCATTCAATACCGCCAGAGCTG TGGT1_310410 reverse
CCCACTACGCACACCCGAACACCGCCAGGAACCGGTTTTCCGTCCAGATGAGCAGCATCC
GCAACAGACGTTTTGATCGCGTCCACACGCTCCACCGAAAAAATTTGTGTCTCGTCGATG
GTTGTAGACTCGCCCCAATATCGCCGCTGCTGGCATTCACGCGACCGCGCATTTAACCTC
TGGGCGGTAAGCGATGGTTTTCTTGTGGCACATGAACGCTGGATAGGCTCCGCTCCTTTC
ACCAGTCGAGATGAACTATTTTCGTTCCCATCGCATCCACCAGAGGCCACACCCGGACGA
ATAGTCCACACCTGCACCCATCGATCCCAGCCTACAGTAAACAGAAGGGGGAAGAGAAGC
GAGCAACTTCGTGCTGACGACGCATGAGCGTTTTCTACTCTACAGGCGGATATCAACCTA

                         T                                   RH
AGGGAGAGAGACGCGTTGTTCTCCTCGTGTTCTACGCAAATAAACTGGACCGTTACTGAC
                 caagaggaacacaagatgcgtcgccggcgc-5'           Rb

TGGTCATCACCGCACGAGGCGACGCAGTAGCAAGGCGACTCCATTTCAGACTCTGCGAGG
                              gttccgctgaggtaaagtct-5'        Re

GCGGACGAACCCAGCCCTGCAGGAGATGCGACCGACGCGCCGCTTTCAACCTTTGTCCCC
AAAGCGTTTTGCGTTTCGGCCACTTCCAAGTCATTTACGCCACACTGGTGTGCGTGTAGA
GTGCCCAATCGCTCCAGTCGAGGCACCGCAGCGCTCGTGTCCCCCAGGAACACCGCGATT
TCTCCGGTGGTCATTCCGACCAGTACCAAGACCCCATGTGTGTTCGTTCTCGACGGTTCC
TCACTCCCGAACTCTGTCTGGGAGCTAGCAAAGGAGCAGCCTCCAGCATCGTTTGCATGC
GTGTCTAGACAAACGACACGAACGCAGAGAGCCGCGCCTTCCAGCGGGCATGATGCTCTT
TTTTCTAAGCGACTGTTCCGGCTCCGGTCACCATGAGCCAGCTGCCCGAATTCCTGAGAC
CTCGTCAGCACAAGTGTCGCTTCCACATACGCGACTTCGGCCGACGACGCAGCCGTCCAC
ATGTGCACAGTCAGAGACTC
```

**Fig. S2. *Pythium ultimum* Gat1 sequences**

**Bold: amino acid sequence**

Black: native genomic coding sequence, from PYU1_G002535-201 (UniProtK3WCV7)

Red: Synthetic codon optimized sequence

**M T V G T R R A A Y A T L I T S D A Y V**

atgaccgtcggcacgcgcagggcggcgtacgcaacactgatcacgtccgatgcgtacgtc

ATGACTGTCGGAACACGTCGTGCGGCTTATGCCACTTTGATCACCAGCGATGCGTACGTT 60

**M G V E A L V Y S L F K A R V A F P L V**

atgggcgtcgaggcgctcgtctactcgctcttcaaggcgcgcgtagcgtttccgctcgtg

ATGGGCGTCGAGGCATTAGTGTATAGCTTGTTTAAGGCGCGTGTTGCCTTCCCACTTGTG 120

**V L H S S Q V T Q P T V A K L T R F C A**

gtgctgcattcgtcgcaggtgacgcagcccacggtggccaaactcacgcgcttctgcgcg

GTGTTACACAGCAGCCAGGTTACTCAGCCAACGGTGGCGAAGCTTACCCGTTTCTGCGCC 180

**P F Q S S T W R I S F R S V P D I G I P**

ccattccagtcatcaacgtggcgcatttcgttccgctctgtcccagatatcggcatccca

CCCTTCCAAAGCAGCACATGGCGTATTAGCTTTCGTAGCGTTCCTGATATCGGTATCCCA 240

**D E V T D R S T V H V P G W V N S G Y T**

gacgaagtcactgataggagcacggtgcacgtgcctggatgggtcaactcggggtacacc

GACGAGGTAACTGATCGTAGTACCGTCCATGTGCCGGGATGGGTTAATTCAGGTTACACA 300

**K L H I F A M D D F E Q I V Y I D A D A**

aagctccacatcttcgccatggacgactttgagcagatcgtgtacattgacgccgacgcc

AAGCTTCATATCTTCGCTATGGACGACTTCGAGCAAATCGTCTATATTGACGCCGATGCC 360

**I V L Q N V D E L F D R S T S F A A A P**

atcgtcctacagaacgtcgacgagcttttcgatcgctcaacgagctttgcggctgcgccc

ATTGTTCTTCAAAACGTAGACGAGTTGTTCGACCGTAGTACCAGCTTCGCGGCGGCGCCT 420

**D V F P P D R F N A G V L V I R P N K Q**

gacgtgtttccacccgaccgcttcaacgccggcgtgctcgtgatccgtccgaacaagcag

GACGTATTTCCACCAGACCGTTTTAACGCGGGGGTGCTTGTCATTCGTCCTAACAAACAA 480

**L F A D L L A K A K E L K S Y D G G D T**

ctctttgcagacttactggcgaaagccaaggagctcaagtcgtacgatggcggcgacacg

CTTTTCGCCGACTTGTTAGCGAAGGCCAAGGAATTGAAAAGCTATGATGGGGGCGATACA 540

**G F L N A F F P K W F E S D A A S R L P**

ggcttcctcaatgcgtttttccccaagtggttcgaatcggacgccgcgtcgagactgccg

GGATTCTTAAACGCTTTTTTCCCCAAGTGGTTCGAGTCCGATGCCGCCTCACGTTTGCCT 600

**F G Y N A Q R T M Y W L V N G K N P G Y**

tttggatacaacgcgcagcgcacgatgtactggctcgtgaacggcaagaaccccgggtac

TTTGGTTACAATGCTCAGCGTACGATGTACTGGCTTGTGAACGGGAAGAACCCTGGGTAC 660

**W N A V Q P L K I L H Y S S N P K P W E**

tggaacgccgtccagccgctcaagatcctgcactactcgtcgaatccaaagccctgggag

TGGAACGCGGTCCAGCCTTTGAAGATTCTTCACTATTCATCCAATCCTAAACCCTGGGAG 720

**D P S R K G D L E I L W W Q M Y T E S R**

gatccgagtcgcaagggtgacctggagatcctgtggtggcaaatgtacacggaatccaga

GACCCAAGTCGTAAGGGTGACTTGGAGATTCTTTGGTGGCAAATGTATACCGAAAGTCGT 780

**C M S F L G ***

tgcatgagctttctggggtag

TGTATGAGCTTTCTTGGATAG 801

**Fig. S3.** Disruption and complementation of Tg*gat1* in the RHΔΔ type 1 strain. A. Disruption of *gat1* by double cross-over homologous recombination in RHΔΔ. The disruption DNA consisted of an HXGPRT cassette flanked by a 1177-nt 5'-upstream DNA (prepared by PCR using primers Fa and Ra) and a 1209-nt 3'-downstream DNA (prepared using primers Fb and Rb) of the *gat1* coding sequence (CDS), and an adjacent GFP-expression cassette. GFP-negative clones that grew in mycophenolic acid (MPA) and xanthine showed evidence of a gel shift of Skp1 relative to parental and *phyA*Δ cells based on Western blotting using pAb UOK75. Clone A1 was confirmed to have the desired integration by PCR reaction #1 (primers Fc and Rc), which showed loss of the *gat1* CDS, and PCR2 (Fd and Rd) and PCR3 (Fe and Re), which showed integration of the HXGPRT within the gat1 locus. **B.** Disruption of *gat1* using a double CRISPR/Cas9 strategy. RHΔΔ parasites were transiently transfected with a plasmid encoding gDNA-63 and gDNA-968 guide DNA's and Cas9, and a PCR amplicon expressing the DHFR resistance cassette flanked by 45-bp *gat1* homology arms. Pyrimethamine-resistant clones that replaced the *gat1* CDS with the DHFR cassette were confirmed using PCR4 (Fh and Rh), which showed loss of *gat1* CDS, and PCR2 and PCR3, which showed the integration of DHFR in the forward orientation. **C.** Complementation of clone B7 from panel B by replacement of the *uprt* locus with a *gat1* expression cassette consisting of a tubulin promoter, Tg*gat1* CDS modified with DNA encoding a C-terminal 3×HA tag, and *uprt* targeting sequences. Correct insertion of the gat1 expression cassette was assessed by PCR reactions PCR7, PCR8 and PCR9, and confirmed by Western blot analysis for a predicted $M_r$ 45,000 protein band that could be detected with mAb 12CA5 that recognizes the 3xHA epitope. A parallel gel containing samples without reducing reagent was Western blotted to detect Sag1 as a loading control.

**Figure S3.** Disruption and complementation of Tg*gat1* in the RHΔΔ type 1 strailn

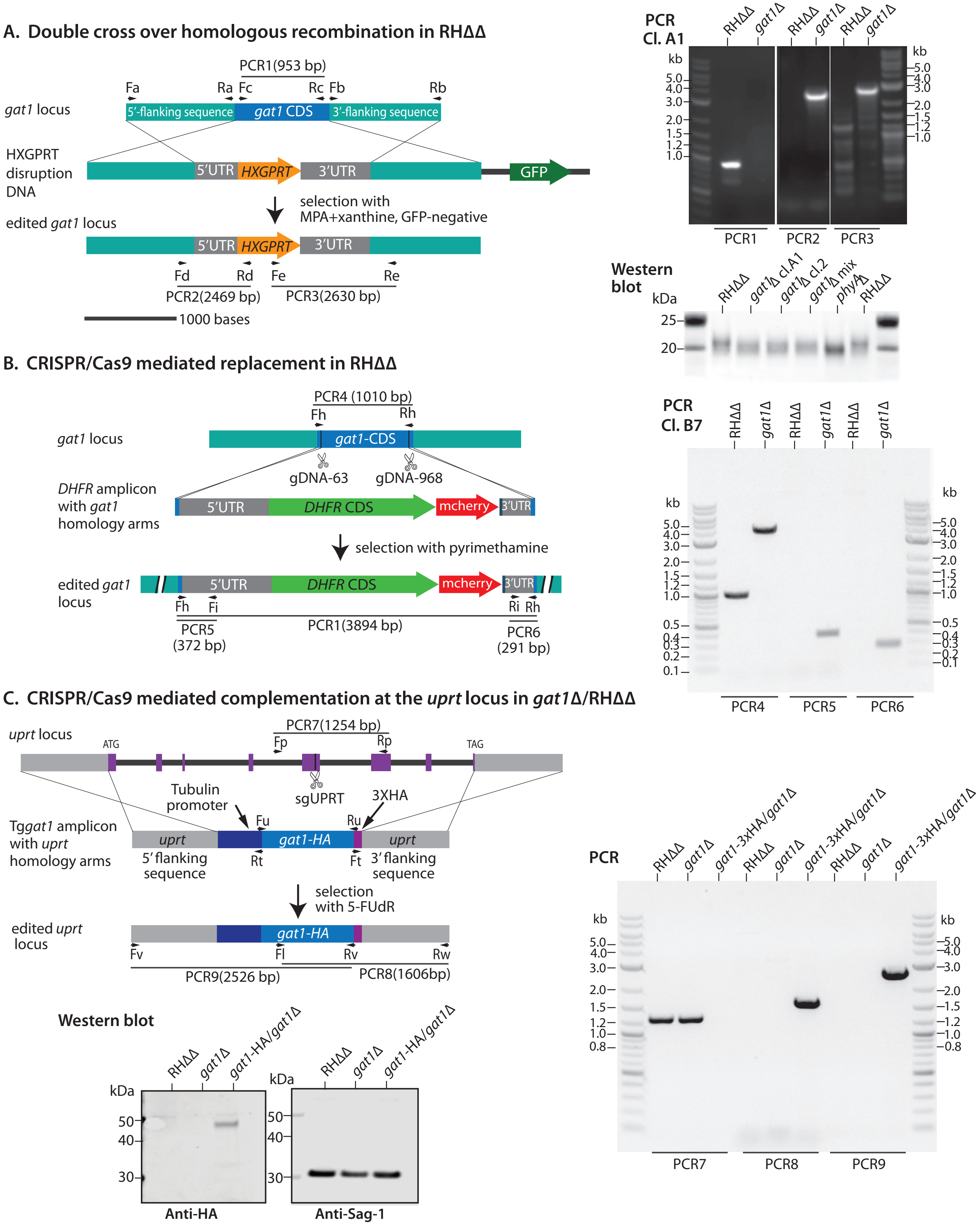

**Fig. S4.** Disruption and complementation of Tg*gat1* in Ku80+ type 1 and type 2 strains. A. Disruption of *gat1* was achieved using the double CRISPR/Cas9 strategy described in Fig. S3B, except that the DHFR amplicon lacked *gat1* homology arms owing to the presence of non-homologous end joining activity. Successful replacement was evaluated for strains RH and ME49, by PCR as in Fig. S3B. **B.** The RH *gat1*Δ strain was complemented by insertion of a genomic fragment of Tg*gat1* including its CDS, DNA encoding a C-terminal Ty-tag, and >1 kb of flanking DNA from both directions. Successful integration was verified using *uprt*-specific primers (Fp and Rp, Table S1) flanking the CRISPR/Cas9 cut site in PCR reaction #4. The identity of the integrated DNA was verified using primer pairs Fp and Rq, and Fq and Rp, in which Rq and Fq were specific to Tg*gat1* DNA. **C.** Extracts of *gat1*Δ and complemented clones from panels A and B were analyzed for Skp1 αGalT activity. Desalted S100 extracts were prepared by hypotonic lysis and gel filtration, and incubated in the presence of GlFGaGn-Skp1 and UDP-[$^3$H]Gal. The reactions were separated on SDS-PAGE gels, the Skp1 band was excised after Coomassie blue staining, and radioactivity determined by liquid scintillation counting. Error bars represent S.D. of two technical replicates of the same samples.

**Figure S4.** Disruption and complementation of Tg*gat1* in Ku80$^{+}$ type 1 and type 2 strains

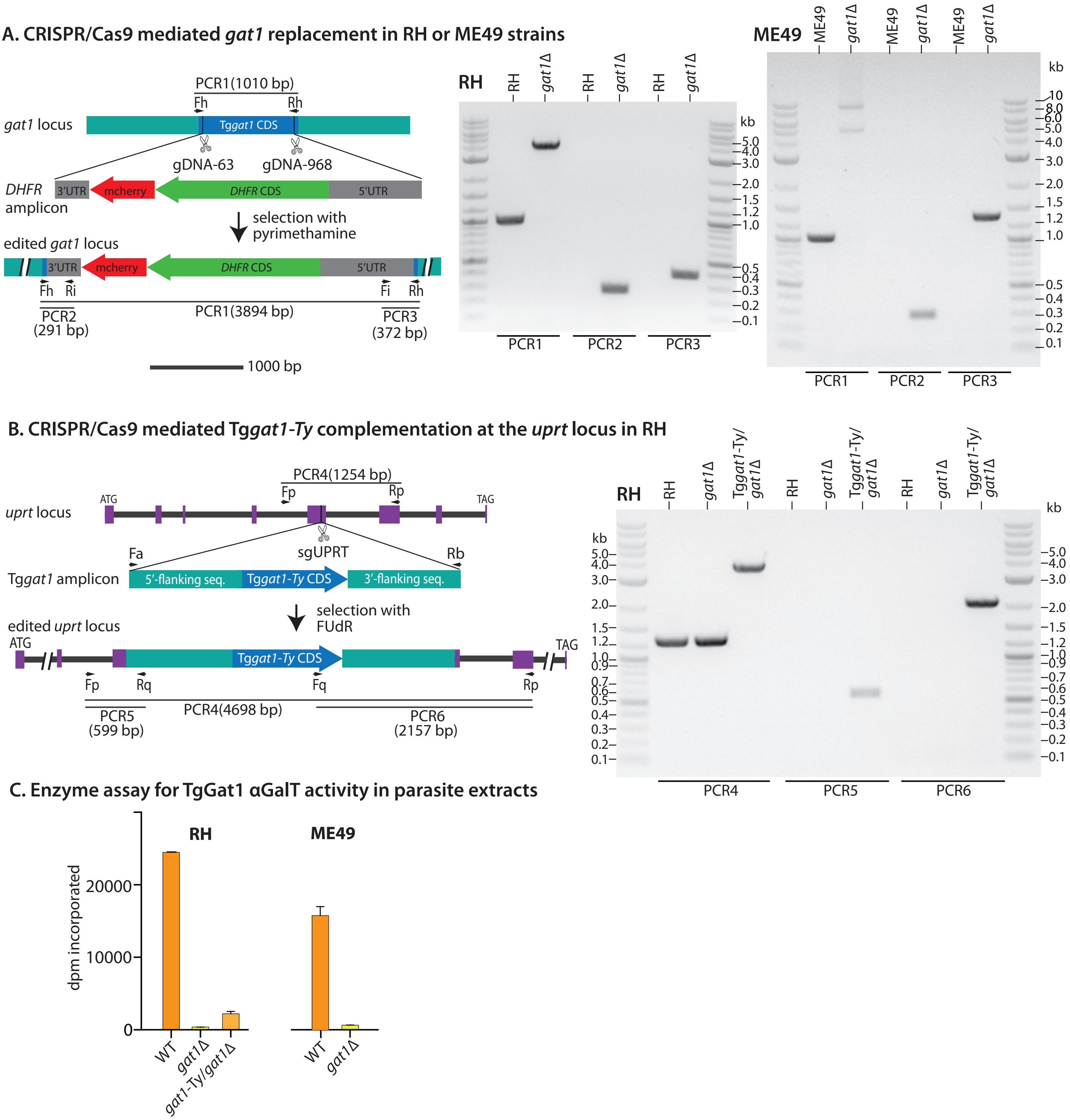

**Fig. S5.** nLC/MS of Skp1 glycopeptides (supports Fig. 1C, Table S2)
TgSkp1 isolated by immunoprecipitation from tachyzoite extracts were reduced and alkylated, trypsinized treated with green coffee bean a-galactosidase as indicated, and analyzed by a standard proteomics workflow consisting of separation on a C18 nLC column and analysis in an QE-Plus Orbitrap mass spectrometer.

Samples analyzed (as described in Fig. 1C):
RH (type 1 parental)
*gat1*Δ/RH
Tg*gat1*/*gat1*Δ/RH (complemented under the tubulin promoter in the uprt locus)
RH, incubated with α-galactosidase
Me49 (type 2 parental)
*gat1*Δ/Me49

**A.** Stacked extracted ion chromatograms for all isoforms of peptide(134-150), which contains the modifiable Pro143, that were detected in the RH and Me49 backgrounds.
**B.** Selected extracted ion chromatograms for unmodified peptide(134-150), and an example of an $MS^1$ spectrum, from the RH sample.
**C.** Selected extracted ion chromatograms for pentasaccharide-modified peptide(134-150), and an example of an $MS^1$ spectrum, from the TgGat1 complemented sample.
**D.** Selected extracted ion chromatograms for tetrasaccharide-modified peptide(134-150), and an example of an $MS^1$ spectrum, from the TgGat1 complemented sample.
**E.** $MS^2$ of unmodified peptide(134-15) from RH, with associated extracted ion chromatogram and MS1. Detected b and y fragment ions that define the peptide sequence are in bold in the list of predicted fragments ions at the bottom.
**F.** $MS^2$ of pentasaccharide peptide(134-15) from RH, with associated extracted ion chromatogram and $MS^1$. An expanded table of predicted b and y fragment ions, calculated to include the full pentasaccharide or a GlcNAc stub, is at the bottom. $MS^2$ fragmentation resulted in either loss of the full glycan leaving Hyp, or retention of a GlcNAc stub (encircled with a dashed green line).
**G.** $MS^2$ of tetrasaccharide peptide(134-15) from *gat1*Δ/RH, with associated extracted ion chromatogram and $MS^1$. An expanded table of predicted b and y fragment ions, calculated to include the full tetrasaccharide or a GlcNAc stub, is at the bottom. $MS^2$ fragmentation resulted in either loss of the full glycan leaving Hyp, or retention of a GlcNAc stub (encircled with a dashed green line).

### Figure S5A,B

A. Extracted ion chromatogram Summary: IFNIVNDFT(HyP)EEEAQVR (all glycoforms)

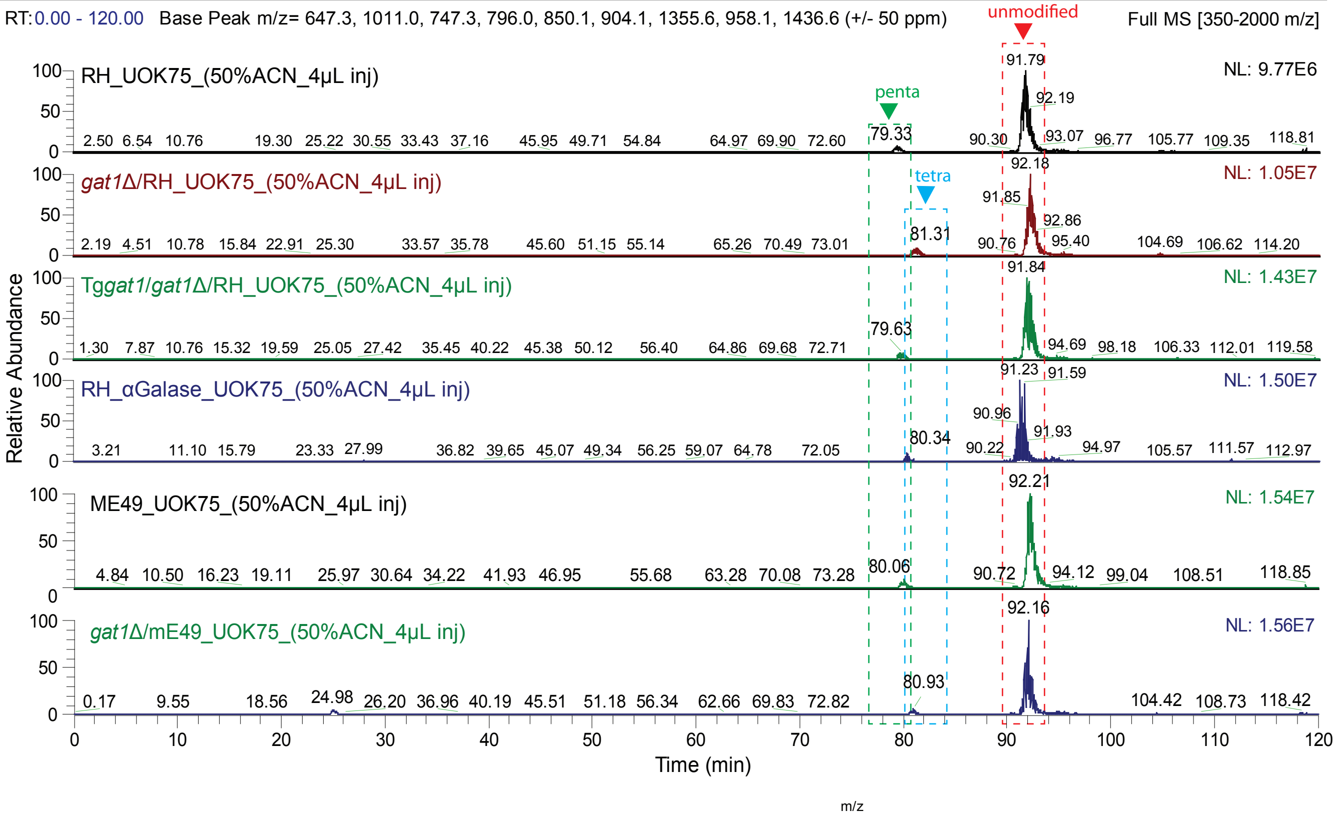


B. Extracted ion chromatograms, MS1: IFNIVNDFT(HyP)EEEAQVR (unmodified)

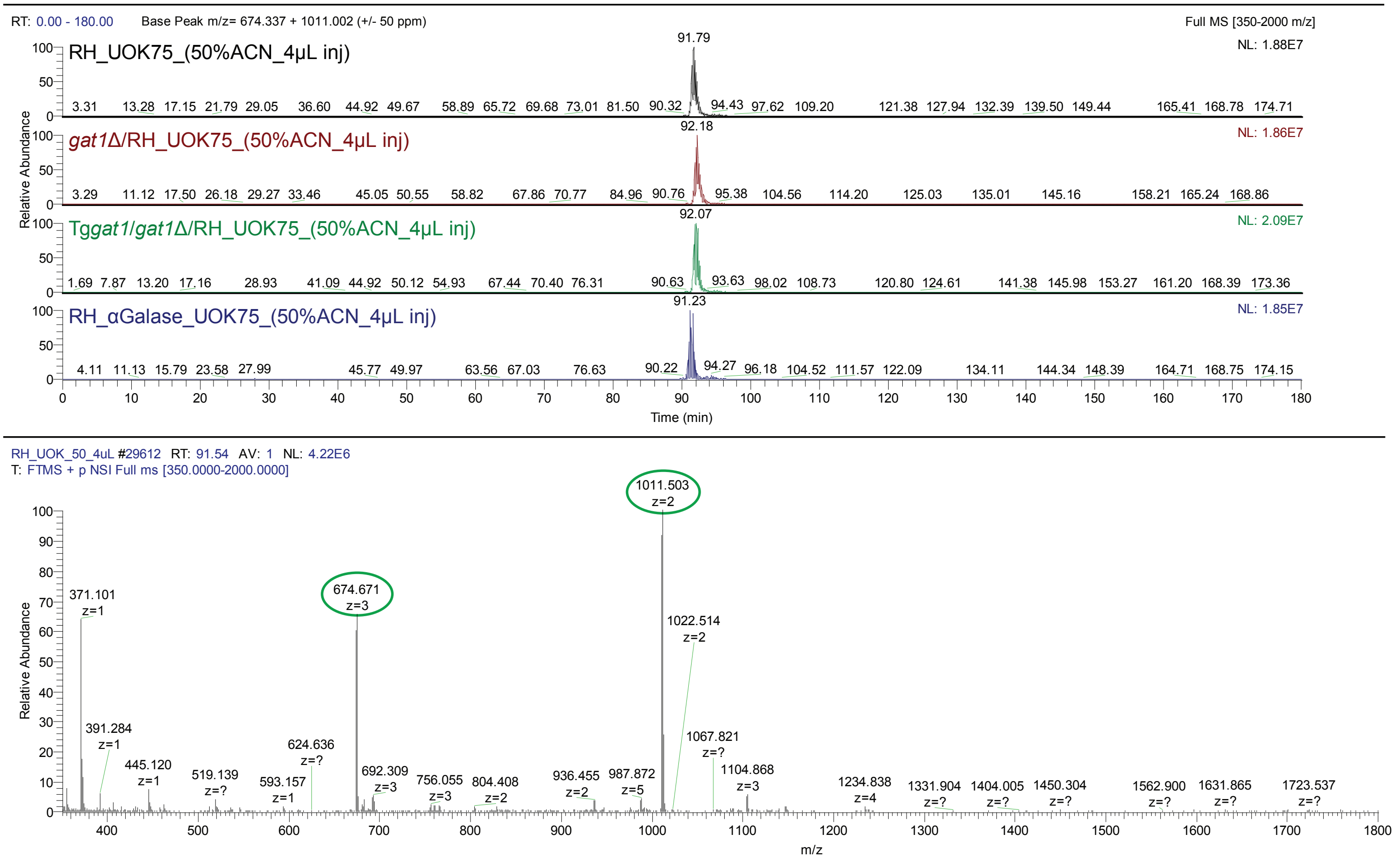

### Figure S5C,D

C. IFNIVNDFT(HyP)EEEAQVR + pentasaccharide

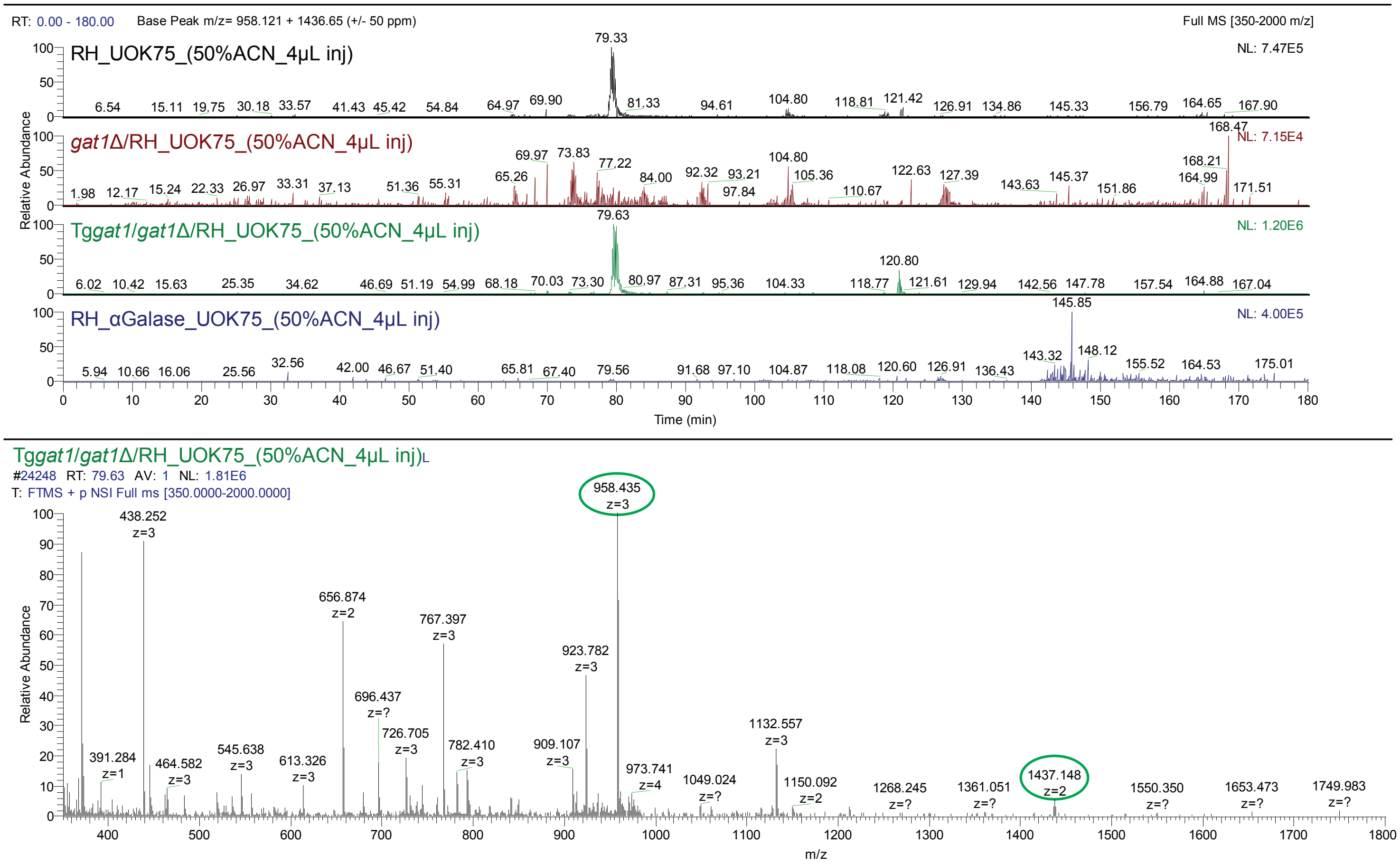


D. IFNIVNDFT(HyP)EEEAQVR + tetrasaccharide

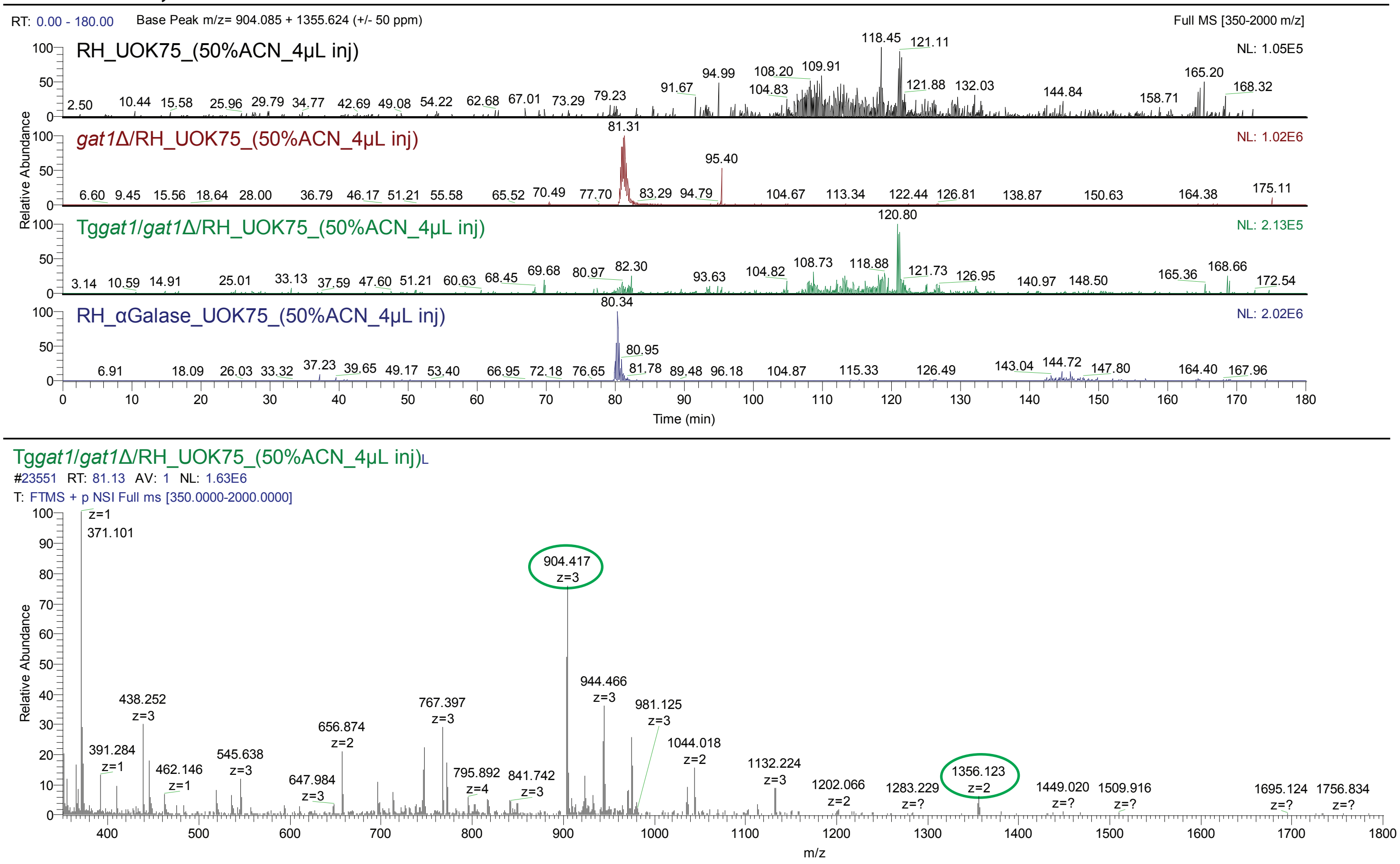

### Figure S5E. MS2 of unmodified peptide from RH

IFNIVNDFT(Pro)EEEAQVR (unmodified)

RH_UOK75_(50%ACN_4µL inj)

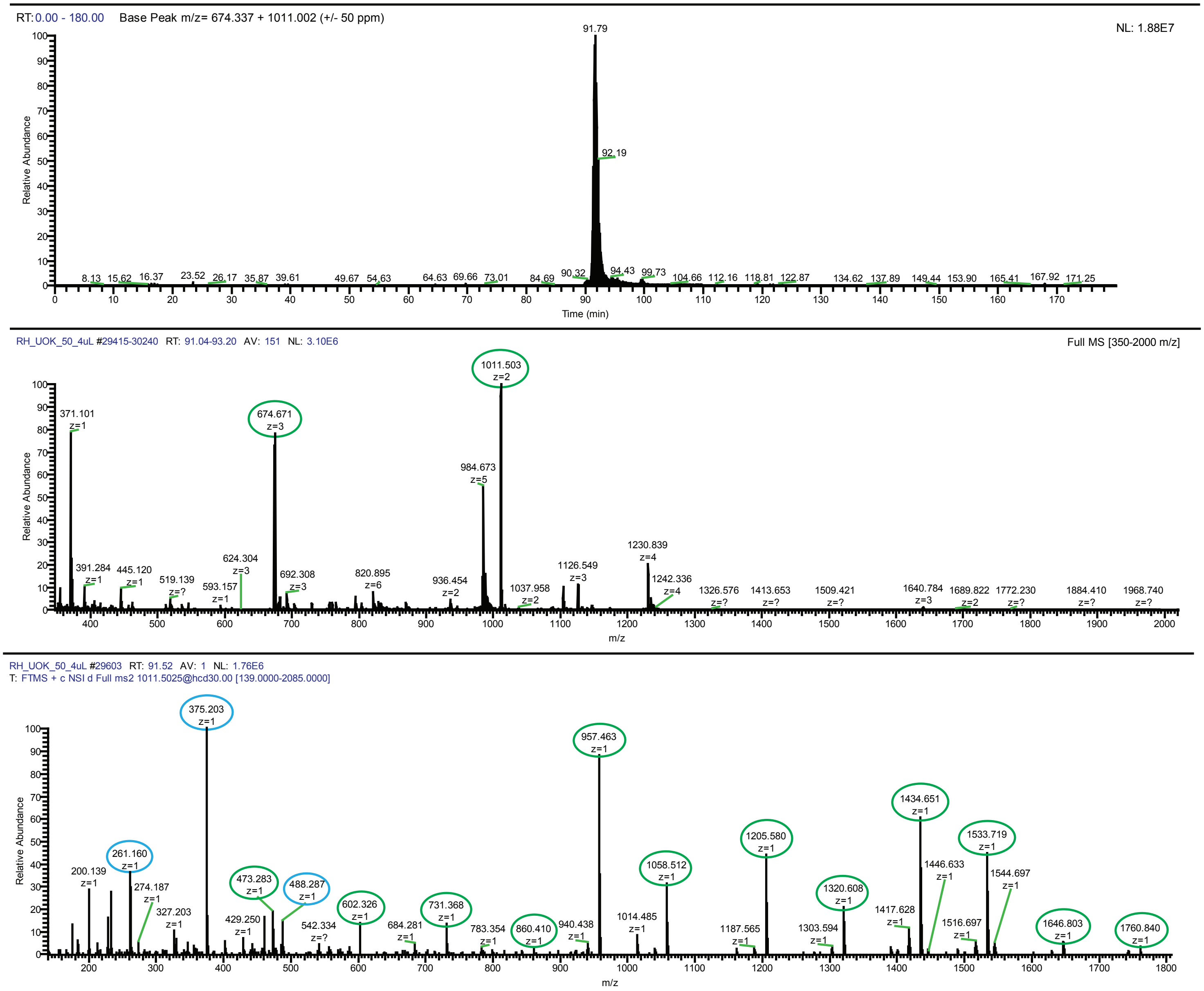


IFNIVNDFT(Pro)EEEAQVR

| | | unmodified | | |
|---|---|---|---|---|
| | | b | y | |
| I | 1 | 114.0914 | 2020.997 | 1 |
| F | 2 | **261.1598** | 1907.913 | 2 |
| N | 3 | **375.2027** | **1760.845** | 3 |
| I | 4 | **488.2868** | **1646.802** | 4 |
| V | 5 | 587.3552 | **1533.718** | 5 |
| N | 6 | 701.3981 | **1434.65** | 6 |
| D | 7 | 816.4251 | **1320.607** | 7 |
| F | 8 | 963.4935 | **1205.58** | 8 |
| T | 9 | 1064.541 | **1058.511** | 9 |
| P | 10 | 1161.594 | **957.4636** | 10 |
| E | 11 | 1290.637 | **860.4109** | 11 |
| E | 12 | 1419.679 | **731.3683** | 12 |
| E | 13 | 1548.722 | **602.3257** | 13 |
| A | 14 | 1619.759 | **473.2831** | 14 |
| Q | 15 | 1747.817 | 402.246 | 15 |
| V | 16 | 1846.886 | 274.1874 | 16 |
| R | 17 | 2002.987 | 175.119 | 17 |

b fragments in blue, y fragments in green; detected fragments bold.

### Figure S5F. MS2 of pentasaccharide peptide from RH

IFNIVNDFT(HyP+HexNAc+Fuc+Hex3)EEEAQVR

RH_UOK75_(50%ACN_4µL inj)

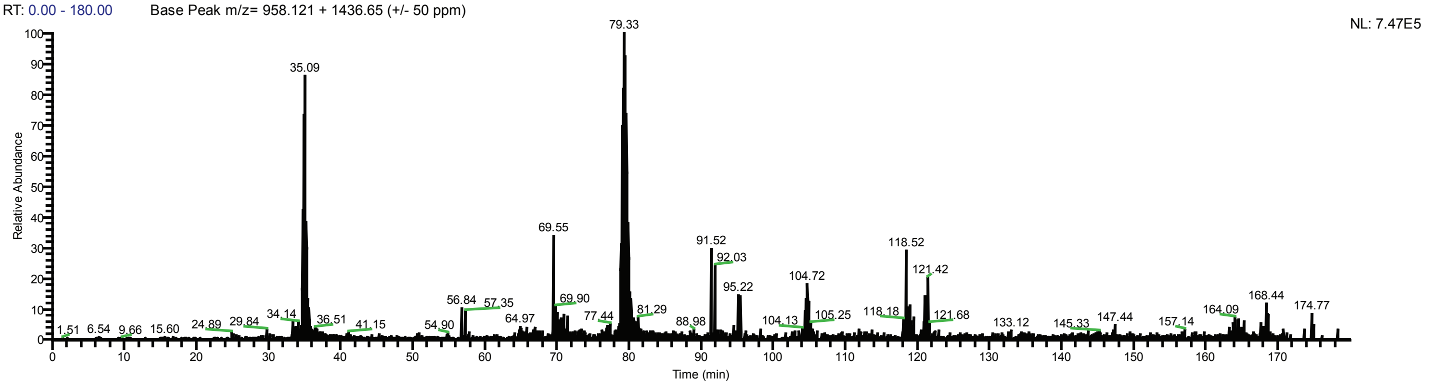


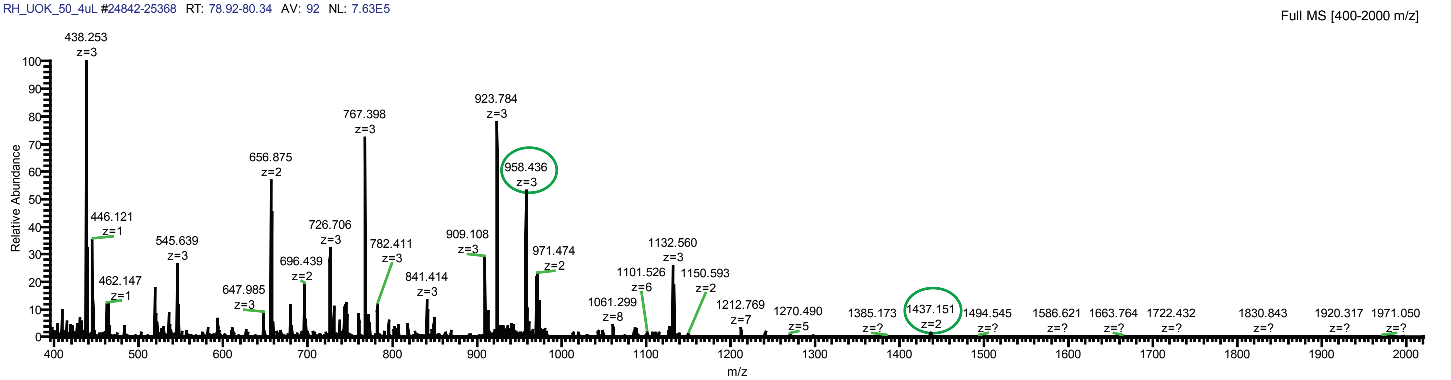


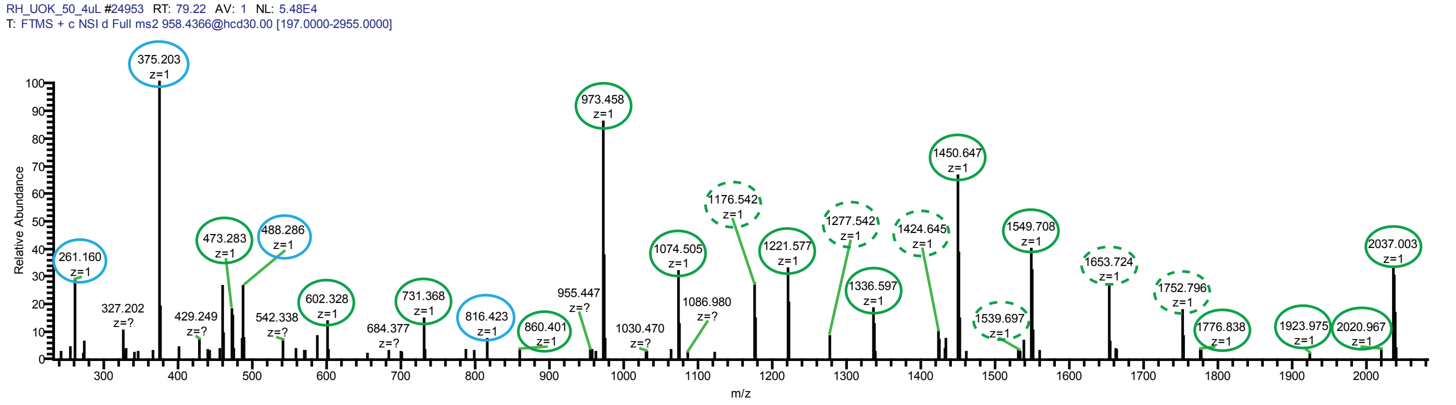


IFNIVNDFT(Pro)EEEAQVR

| | | unmodified | | Hyp | | Hyp+HexNAc | | Hyp+penta | | |
|---|---|---|---|---|---|---|---|---|---|---|
| | | b | y | b | y | b | y | b | y | |
| **I** | 1 | 114.0914 | **2020.997** | 114.0914 | **2036.997** | 114.0914 | 2239.997 | 114.0914 | 2872.293 | 17 |
| **F** | 2 | **261.1598** | 1907.913 | **261.1598** | 1923.913 | **261.1598** | 2126.913 | **261.1598** | 2759.209 | 16 |
| **N** | 3 | **375.2027** | 1760.845 | **375.2027** | **1776.845** | **375.2027** | 1979.845 | **375.2027** | 2612.141 | 15 |
| **I** | 4 | **488.2868** | 1646.802 | **488.2868** | 1662.802 | **488.2868** | 1865.802 | **488.2868** | 2498.098 | 14 |
| **V** | 5 | 587.3552 | 1533.718 | 587.3552 | **1549.718** | 587.3552 | **1752.718** | 587.3552 | 2385.014 | 13 |
| **N** | 6 | 701.3981 | 1434.65 | 701.3981 | **1450.65** | 701.3981 | **1653.65** | 701.3981 | 2285.946 | 12 |
| **D** | 7 | **816.4251** | 1320.607 | **816.4251** | **1336.607** | **816.4251** | **1539.607** | **816.4251** | 2171.903 | 11 |
| **F** | 8 | 963.4935 | 1205.58 | 963.4935 | **1221.58** | 963.4935 | **1424.58** | 963.4935 | 2056.876 | 10 |
| **T** | 9 | 1064.541 | 1058.511 | 1064.541 | **1074.511** | 1064.541 | **1277.511** | 1064.541 | 1909.807 | 9 |
| **P** | 10 | 1161.594 | 957.4636 | 1177.594 | **973.4636** | 1380.594 | **1176.464** | 2012.89 | 1808.76 | 8 |
| **E** | 11 | 1290.637 | 860.4109 | 1306.637 | 860.4109 | 1509.637 | 860.4109 | 2141.933 | 860.4109 | 7 |
| **E** | 12 | 1419.679 | **731.3683** | 1435.679 | **731.3683** | 1638.679 | 731.3683 | 2270.975 | **731.3683** | 6 |
| **E** | 13 | 1548.722 | **602.3257** | 1564.722 | **602.3257** | 1767.722 | 602.3257 | 2400.018 | **602.3257** | 5 |
| **A** | 14 | 1619.759 | **473.2831** | 1635.759 | **473.2831** | 1838.759 | 473.2831 | 2471.055 | **473.2831** | 4 |
| **Q** | 15 | 1747.817 | 402.246 | 1763.817 | 402.246 | 1966.817 | 402.246 | 2599.113 | 402.246 | 3 |
| **V** | 16 | 1846.886 | 274.1874 | 1862.886 | 274.1874 | 2065.886 | 274.1874 | 2698.182 | 274.1874 | 2 |
| **R** | 17 | 2002.987 | 175.119 | 2018.987 | 175.119 | 2221.987 | 175.119 | 2854.283 | 175.119 | 1 |

b fragments in blue, y fragments in green; detected fragments in bold.
Specific HexNAc fragments dashed in green.
No specific pentasaccharide fragments detected.

### Figure S5G. MS(2) of tetrasaccharide peptide fromm *gat1*Δ/RH

IFNIVNDFT(HyP+HexNAc+Fuc+Hex2)EEEAQVR

*gat1*Δ/RH_UOK75_(50%ACN_4μL inj) 03/28/19 20:20:12

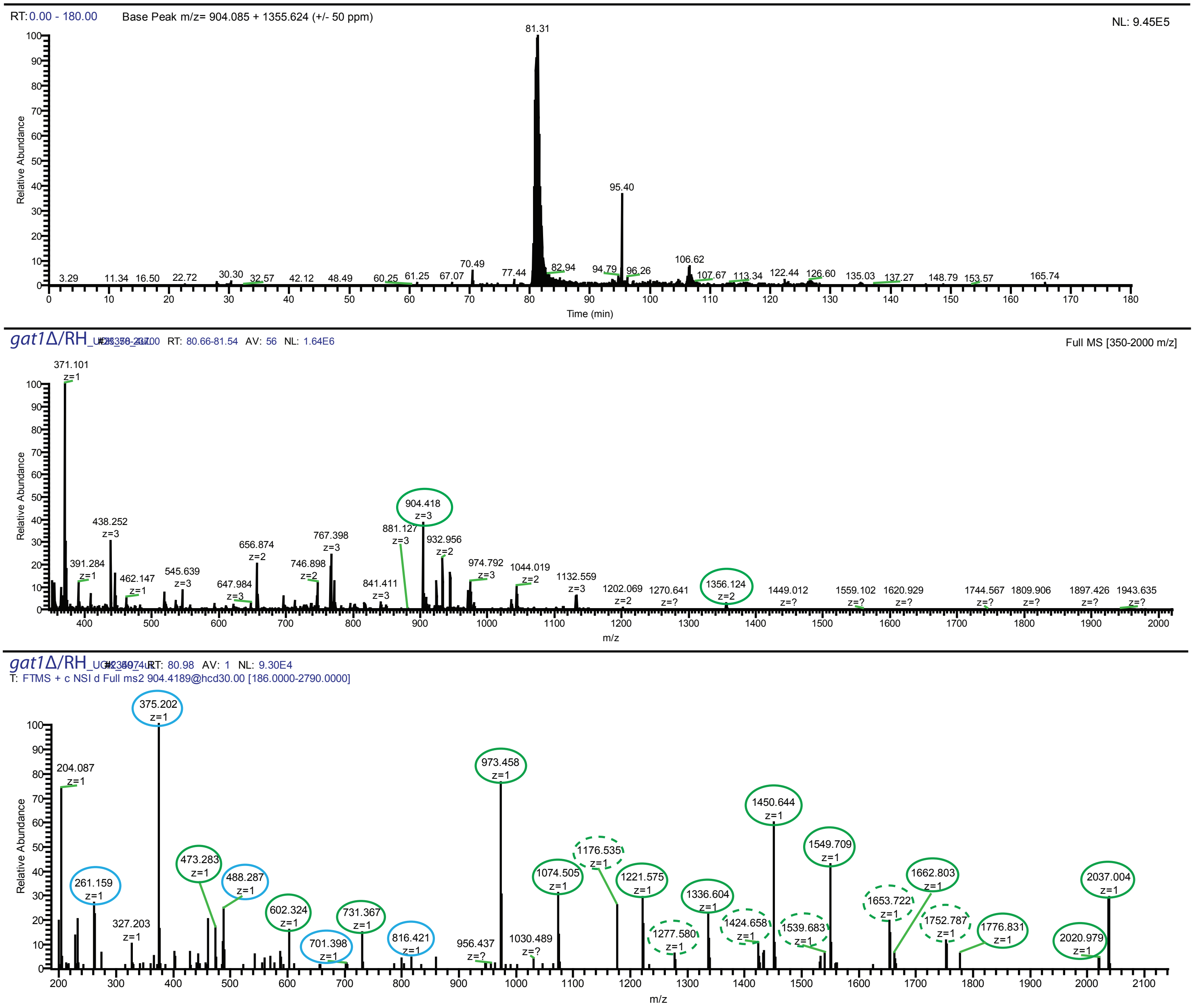


IFNIVNDFT(Pro)EEEAQVR

| | | unmodified | | Hyp | | Hyp+HexNAc | | Hyp+tetra | | |
|---|---|---|---|---|---|---|---|---|---|---|
| | | b | y | b | y | b | y | b | y | |
| I | 1 | 114.0914 | **2020.997** | 114.0914 | **2036.997** | 114.0914 | 2239.997 | 114.0914 | 2710.24 | 17 |
| F | 2 | **261.1598** | 1907.913 | **261.1598** | **1923.913** | **261.1598** | 2126.913 | **261.1598** | 2597.156 | 16 |
| N | 3 | **375.2027** | 1760.845 | **375.2027** | **1776.845** | **375.2027** | 1979.845 | **375.2027** | 2450.088 | 15 |
| I | 4 | **488.2868** | 1646.802 | **488.2868** | **1662.802** | **488.2868** | 1865.802 | **488.2868** | 2336.045 | 14 |
| V | 5 | 587.3552 | 1533.718 | 587.3552 | **1549.718** | 587.3552 | **1752.718** | 587.3552 | 2222.961 | 13 |
| N | 6 | **701.3981** | 1434.65 | **701.3981** | **1450.65** | **701.3981** | **1653.65** | **701.3981** | 2123.893 | 12 |
| D | 7 | **816.4251** | 1320.607 | **816.4251** | **1336.607** | **816.4251** | **1539.607** | **816.4251** | 2009.85 | 11 |
| F | 8 | 963.4935 | 1205.58 | 963.4935 | **1221.58** | 963.4935 | **1424.58** | 963.4935 | 1894.823 | 10 |
| T | 9 | 1064.541 | 1058.511 | 1064.541 | **1074.511** | 1064.541 | **1277.511** | 1064.541 | 1747.754 | 9 |
| P | 10 | 1161.594 | 957.4636 | 1177.594 | **973.4636** | 1380.594 | **1176.464** | 1850.837 | 1646.707 | 8 |
| E | 11 | 1290.637 | 860.4109 | 1306.637 | 860.4109 | 1509.637 | 860.4109 | 1979.88 | 860.4109 | 7 |
| E | 12 | 1419.679 | 731.3683 | 1435.679 | **731.3683** | 1638.679 | **731.3683** | 2108.922 | **731.3683** | 6 |
| E | 13 | 1548.722 | 602.3257 | 1564.722 | **602.3257** | 1767.722 | **602.3257** | 2237.965 | **602.3257** | 5 |
| A | 14 | 1619.759 | 473.2831 | 1635.759 | **473.2831** | 1838.759 | **473.2831** | 2309.002 | **473.2831** | 4 |
| Q | 15 | 1747.817 | 402.246 | 1763.817 | 402.246 | 1966.817 | 402.246 | 2437.06 | 402.246 | 3 |
| V | 16 | 1846.886 | 274.1874 | 1862.886 | 274.1874 | 2065.886 | 274.1874 | 2536.129 | 274.1874 | 2 |
| R | 17 | 2002.987 | 175.119 | 2018.987 | 175.119 | 2221.987 | 175.119 | 2692.23 | 175.119 | 1 |

b fragments in blue, y fragments in green; detected fragments in bold.
Specific HexNAc fragments dashed in green.
No specific tetrasaccharide fragments detected.

**Figure S6.** Alignment catalytic domains of Gat1-like sequences and glycogenins

green names- glycogenin like
blue names- Gat1-like

```
                   ‡‡‡‡  ‡                                                                            ¥ ¥
Oc   1-----MTDQAFVTLTTNDAYAKGALVLGSSLKQHRTSR-------RLAVLTT---PQVSDTMRKALEIVFDEV----ITVDILD------SGDSAHLTLMK
Dm   1-----MSKFAWVTLTTNDTYSLGALVLAHSLKRAKTAH-------QLAVLVT---PNVSQAMRDRLKEVYNVV----QEVNVLD------SQDAANLALLS
Ta   1---MSEKREAFVTLATNDSYAVGAFVLGNSLRNVKTTR-------ELVVLIT---DEVTHHYRYRLRHVFDIV----KLVDPFD------SGDEKHLRLLG
Sc   1--MGMYKKLAIATLLYSADYLPGVFALGHQVNKLLEEAGKKG-DIETCLIVT[5]GTLSELAKNILQSIYTKI----VLVEPLNCQEESIQKNSENLALLE
Tg   1----MSPRYAYATLLTDNSFYYGVEALLKSLEATKTPY-------PVLLLHT---SDVSQSTIKALVYQRRKA[97]RLVGSVA-----YPKAERDTCPVE
Ot   1-------------MITDDGYLPGLQVLHYTLRKFTSRL-------LVIILA---ENVKKITEMQIKKLSNVM---IKRVKPIL-----NPHEKSQTDNAS
Tp  37-APFPPRPKAIATFLSSADFLPGCQTLLHSLKKQLPQTPKDEYPPEIIVLLS---SKTSN--RQAIESRLHPTFC--ISVDHIP[11]DKGSSEKQMSHVQ
Pu   1-MTVGTRRAAYATLITSDAYVMGVEALVYSLFKARVAF-------PLVVLHS---SQVTQPTVAKLTRFCAPFQS--ISFRSVP-DIGIPDEVDRSTVHVP

     ¥¥       §               #‡#                      ¥   A ¥¥  ¥  ¥                            ‡   ¥    ¥
Oc   RPELGVTLTKLHCWSLTQYSKCVFMDADTLVL-ANIDDLFE-----REELSAAPDPGWPDCFNSGVFVYQPSVETYNQLLHVA---SEQGSFDGGDQGLLNT
Dm   RPELGVTFTKLHCWRLVQFEKCVFLDADTLVL-QNCDELFE-----REELSAAPDVSWPDCFNSGVFVFKPSVDTFAQITEFA---VKNGSFDGGDQGLLNQ
Ta   RPDLGITLTKLHCWRLTEFSKAVFLDADTLVI-GNIDDLFT-----RPELSAAPDVGWPDCFNSGVFVYKPSMQTYQTIVAFA---LQFGSFDGGDQGLLNE
Sc   RPELSFALIKARLWELTQFEQVLYLDSDTLPLNKEFLKLF[5]-QTTSQVGAIADIGWPDMFNSGVMMLIPDADTASVLQNYI---FENTSIDGSDQGILNQ
Tg   GWKD--CFTKLRVWEQVDFDVIVYVDADCIVL-RPVDELFL----RQPLPAFAPDIFPPDKFNAGVAVLKPDLGEYGNMVAAV---ERLPSYDGGDTGFLNA
Ot   SWVG-SGYTKLYIWTLIQFQKVFYIDADCLIS-SNPENAFD----RNSDFAAAPDVFPPDRFNAGVLLIKPSMTVFRDMISKI---LTFPAYDGGDTGFLNA
Tp   AWDENCGWAKLRLFELDGYDTILYIDADCLVV-KDVSHLL[19]QRSGLLAAAPDIFPPDKFNAGVMVLCPSKAVFNDMMARL[5]NSCTSYDGGDTGFLNS
Pu   GWVN-SGYTKLHIFAMDDFEQIVYIDADAIVL-QNVDELFDRS----TSFAAAPDVFPPDRFNAGVLVIRPNKQLFADLLAKA---KELKSYDGGDTGFLNA

                                                           ‡
                          ¥ ¥A   A * AA ¥¥¥  ¥          # ¥¥ ¥‡                         ¥
Oc  FFNSWATTD---IRKHLPFIYNL--SSISI-YSYLPAFK-AFG---ANAKVVHFLGQTKPWNYTYDTKTK[5]GHDPTMTHPQFLNVWWDIFT rabbit
Dm  FFADWSTAD---IKKHLPFVYNV--TAYAS-YCYLPAFK-QFR---DKIKILHFAGKLKPWLIQFNSETKVASVSSEYAHAQDLIQLWWNIFC fruit fly
Ta  FFNTWATSD---INTHLPFTYNM--TATSA-YWYAPALN-RFS---KDIKVVHFIGALKPWHHLYNKDTGHL[8]GQQPFLTNYVQRWWEIYT early metazoan
Sc  FFNQNCCTD[9]EWVQLSFTYNV--TIPNLGYQSSPAMN-YFK---PSIKLIHFIGKHKPWSLWS----------QKNFIKNEYHDQWNEVYE yeast
Tg  YFSSWYENA---AGARLPFRYNALRTLYHMTYSSRKGYW-DAV---KPIKILHFCSSPKPWEQ---------------PAKTDLEELWWKVFL apicomplexan parasite
Ot  YYPDWYLKD---SDSRLPYGYNAQRTLYWFTIKRTDGYW-KEI[4]EGLVIIHYSSSPKPWVG---------------QPKGDLELLWFQTYM ciliated protozoan
Tp  YYPNWFGGMP--EYSRLSFGYNAQRFMHHCTYEKQPKYWDDGI---DDVYIVHFSSSPKPWETKSSNEASD[15]QKAVKHGTLESKWQLAFD marine diatom
Pu  FFPKWFESD---AASRLPFGYNAQRTMYWLVNGKNPGYW-NAV---QPLKILHYSSNPKPWED--------------PSRKGDLEILWWQMYT plant parasite
```

### metal binding
§ present in all GT8 sequences and catalytically essential
‡ & ¥ sugar nucleotide binding (either nucleotide or sugar)
¥ group-specific
A & ¥ hydrogen bond and hydrophobic packing contacts with GlFGaGn- acceptor in PuGat1
* autoglucosylation site in glycogenin

Oc: *Oryctolagus cuniculus*
Dm: *Drosophila melanogaster*
Ta: *Trichoplax adhaerens*
Sc: *Saccharomyces cerevisiae*

Tg: *Toxoplasma gondii*
Ot: *Oxytricha trifallax*
Tp: *Thalassiosira pseudonana*
Pu: *Pythium ultimum*

**Fig. S7.** Summary of Gat1-related sequences selected for phylogenetic analysis. The best scoring hits (based on BLAST) from different categories of Gat1-like sequences were selected for manual alignment and phylogenetic analysis. **A.** Predicted Gat1 sequences, from protists that have PgtA-like sequences but not AgtA-like sequences. **B.** Glycogenin and glycogenin-like sequences. **C.** Closest CAZy GT8 sequences from vascular plants. **D.** Closest CAZy GT8 sequences from organisms (protists) that possess Gnt1-like but not PgtA-like sequences. **E.** Closest CAZy GT8 from protists that possess PgtA-like sequences but lack apparent Gat1. **F.** Closest CAZy GT8 sequences from prokaryotes. Expect values, gene IDs, and known functions are indicated.

| **A. Gat1-like sequences from PgtA containing Protists** | **B. Glycogenin-like sequences** |
|---|---|
| *Toxoplasma gondii* EPR60889.1 | *Trichoplax adhaerens* ($E^{-27}$) XP_002116183.1<br>*Amphimedon queenslandica* ($E^{-30}$) XP_003383748.1<br>*Nematostella vectensis* ($E^{-24}$) XP_001625718.1<br>(Simplest animals) |
| *Hammondia hammondii* XP_008886569.1 | |
| *Neospora caninum Liverpool* XP_003885051.1 | |
| *Ectocarpus siliculosus* CBJ26265.1 | *Saccharomyces cerevisiae* ($E^{-13}$) (yeast) E7QGE5<br>(known function: primes glycogen synthesis) |
| *Albugo laibachii* CCA19642.1 | *Monosiga brevicollis* ($E^{-27}$) (choanoflagellate) XP_001744585.1 |
| *Vitrella brassicaformis* CEM34465.1 | *Capsaspora owczarzaki* (Filesterea) XP_004349815.2 |
| *Nannochloropsis gaditana* EWM28655.1 | *Helobdela robusta* ($E^{-27}$) (anneleid) XP_009013909.1 |
| *Oxytricha trifallaxs* EJY67427.1 | *Drosophila melanogaster* ($E^{-26}$) (fruit fly) NP_001163232.2<br>(known function: primes glycogen synthesis) |
| *Stylonychia lemnae* CDW86810.1 | *Mus Musculus* ($E^{-23}$) (animal) NP_038783.1 |
| *Thalassiosira pseudonana* XP_002291959.1 | *Homo sapiens* ($E^{-23}$) (animal) AAH31096.2<br>(known function: primes glycogen synthesis) |
| *Stylonychia lemnae* CDW86810.1 | **D. Gat1-like sequences (E value <$10^{-5}$) from the protists that have Gnt1 but not PgtA** |
| *Reticulomyxa filosa* X6P0J2 | *Acanthamoeba castellanii* ($E^{-10}$) XP_004352787.1 |
| *Bigowiella natans* JGI: aug1.92_g19606 | *Cyanidioschyzon merolae* ($E^{-22}$) (red Alga) XP_005535960.1 |
| *Sarcocystis neurona* SN3_01500095 | *Galdieria sulphuraria* ($E^{-21}$) (red Alga) XP_005708321.1 |
| *Karenia brevis* EX959504.1 | *Volvox carteri* ($E^{-15}$) (green algae) XP_002954821.1 |
| *Pythium ultimum* K3WC47 | *Phytophthora infestans* ($E^{-13}$) XP_002997946.1 |
| *Aphanomyces euteiches*<br>(Aphanodb2: Ae201684_9096.1) | *Naegleria gruberi* ($E^{-13}$) XP_002672734.1 |
| **C. Closest Gat1-like sequences from plants** | *Saprolegnia diclina* ($E^{-7}$) XP_008603979.1 |
| *Arabidopsis thaliana* ($E^{-16}$) NP_175891.1 | *Chlorella variabilis* ($E^{-10}$) XP_005850943.1 |
| *Oryza sativa* ($E^{-17}$) A2XDA4 | *Trichomonas vaginalis* ($E^{-10}$) XP_001309036.1 |
| **E. Gat1-like GT8 sequences from organisms that have PgtA but not Gat1** | **F. Gat1 like sequence from bacteria** |
| *Dictyostelium discoideum* ($E^{-6}$) Q54L24 | *Rhizobium meliloti* ($E^{-19}$) WP_029616784.1 |
| *Albugo laibachii* ($E^{-14}$) F0W520 | |
| *Bigowiella natans* ($E^{-5}$) | |
| *Guillardia theta CCMP2712* ($E^{-15}$) L1J9Y4 | |

**Figure S8.** Alignment of glycogenin-like, Gat1-like, and other CAZy GT8 sequences used to construct the phylogenetic tree in Fig. 2. The amino acid sequence of Gat1-like proteins described in Fig. S7 (middle panel) were aligned with the amino acid sequences of representative known and predicted glycogenins (top panel) or CAZy GT8 sequences (bottom panel) as described in “Experimental Procedures”. Species names are spelled out at the bottom, and sequence sources are listed in Fig. S7. Amino acids are color-coded with respect to chemical similarities that guided the alignments, giving preference to the registration of hydrophobic residues: green, hydrophobic; blue, acidic; dark red, basic; black, polar; bright red, secondary structure breaking (P or G). Positions occupied by identical amino acids across all the organisms are bolded. Unique motifs that are specific for glycogenins are boxed in blue color, and Gat1-specific motifs are boxed in red.

```
     ....|....| ....|....| ....|....| ....|....| ....|....| ....|....| ....|....|
             10         20         30         40         50         60         70
Hs   QAFVTLTTND AYAKGALVLG SSLKQHRTTR RLVVLATLTL MKRPELGVTL TKLHCWSLTQ YSKCVFMDAD
Mm   QAFVTLTTND AYAKGALVLG SSLKQHRTTR RMVVLTSLTL MKRPELGITL TKLHCWSLTQ YSKCVFMDAD
DM   FAWVTLTTND TYSLGALVLA HSLKRAKTAH QLAVLVTLAL LSRPELGVTF TKLHCWRLVQ FEKCVFLDAD
Mb   QAYVTLCTND AYVVGAMLLA HSLRRTGTRR QIVCMITLGL LQRPELGVTL TKLHAWKLTH YDNCVFLDAD
Hr   -AYVTMATND VYAVGALVLA ETLRQTNTQQ DLVIMITLSL LQRSELGVTF TKIQAWRLVE YRKCVFMDAD
Ta   EAFVTLATND SYAVGAFVLG NSLRNVKTTR ELVVLITLRL LGRPDLGITL TKLHCWRLTE FSKAVFLDAD
Nv   EAFVSLVTND NYANGALVLG YSLRRVNTTR KLALLVTLAL LSRPELGITF TKIRCWNLTH YQKCVFMDAD
Co   EAFVTLVTND GYALGALVLA KSLRDVNTTR KIAVLITLAL LGRPELGVTL TKIYAWKLTQ FTKCVFLDAD
Pm   ETYMTLVLTD SYLIGSQVLA WSLRDSGSKK HLTALVTLYL LGRPDLRSSF TKIHIWAQEK FKKIIYLDAD
Aq   EAYVSLATNN DYCHGAIALA CSLRLTNTSR KLCLLISLAL IKRPELGVTF SKLHIWRLVH YSKCVFLDAD
Sc   LAIATLLYSA DYLPGVFALG HQVNKLKGDI ETCLIVTLAL LERPELSFAL IKARLWELTQ FEQVLYLDSD
-------------------------------------------------------------------------------
Tg   YAYATLLTDN SFYYGVEALL KSLEATKTPY PVLLLHTVGS VAYPKAEDCF TKLRVWEQVD FDVIVYVDAD
Hm   YAYATLLTDN SFYYGVEALL KSLEATKTPY PVLLLHTVGS VAYPKAEDCF TKLRVWEQVD FDVIVYVDAD
Nc   YAYATLLTDN SFYYGVEALL KSLEATKTPY PVLLLYTVGS IAYPEKENCF TKLRAWEQVD FDVIVYIDAD
Sn   KAYATLLLDD SFFYGVAALI RSLAKTRTRY PLLLLHTVEE VRGPAKARLY TKLRLWEQED FDLLVYIDAD
Kb   EAYVSLLTSD SFLMAVQALI ASLKATGTAR RLLLLHTVAA IPNPHQTSGF TKLRVWEQVD FDKLVYIDAD
Vb   CAYITLLTSD SFAIGVETLA FSLRKTGTPH PFIVLVGVDG IANPHAESGF TKLHVWSLTE FQRVVYIDAD
Tp   KAIATFLSSA DFLPGCQTLL HSLKKQLPQT PIIVLLSDNN NSDNNDKCGW AKLRLFELDG YDTILYIDAD
Rf   YAVVSLVTSE SYVVGAQVLI HSLHRNGGLK GSNVLVTVSE IPNPLEKSGY TKLRIFEMVQ LKKLFYIDAD
Bn   YGYVSLLTND GFLPGAIVLA KSLLKVEARY PGAVMVTIPI EPLPCPNVGL TKLRVWQLGD FAKVVYIDAD
Pu   AAYATLITSD AYVMGVEALV YSLFKARVAF PLVVLHSVPD IGIPDEVSGY TKLHIFAMDD FEQIVYIDAD
Ot   -----MITDD GYLPGLQVLH YTLRKF-TSR LLVIILAVKP ILNPHEKSGY TKLYIWTLIQ FQKVFYIDAD
Sl   -----MITED SYLPGLQVMH YSLRKF-TQR TLVVIMTVKP IGNPNEKSGY TKFYIWSLTQ YKRIFYIDAD
Ws   -----MVTSD DFVIGAEVML HSLREHSTRR PLVVMVTVEP IAMPMKRVGY TKLRVWGLIQ FRCVVYIDAD
Ae   KTFATLVTSD DFVIGVQVLA YSLRKHGAKY PLIVLYTVEA LPNPNVHSGY TKLHVFNLVE FSTVFYIDSD
Al   QAYATMITSD DFQMGVEALL YSWSCTHSSI NFLILYTVDS IPIPASSSAY TKLNIFGLEE YQKIVYIDAD
Ng   HAFVTLLTGP GAQVLLHSLR TSISAKVAIR PVVVLVTVEP IANPYAESGF TKLQIWGLTQ FERVVYLDAD
-------------------------------------------------------------------------------
Gt   EAYATLITTK EYIQGAIVLS RIVKSTDEER PFIALVLVPR VKRPTGATTY SKLFVWNLTA YRLVLYLDAD
At   EAYATILHAH VYVCGAIAAA QSIRQSGSTR DLVILVDNPK AEKDAYNWNY SKFRLWQLTD YDKIIFIDAD
Os   EAYATVLHSD TYLCGAIVLA QSIRRAGSTR DLVLLHDNPR AERGTYNYNY SKFRLWQLTD YDRVVFVDAD
Dd   NVYVTFADNA EYLKGIVALR MSMINTKCNY GLIVFVTIEM VDIPKEVPAF TKFRAWQLVE YERVIWLDSD
Tv   YAFATVT-TP AFCMGAVVLG YTLRKYGNDY SYLCLVTVND A-KPYLWRSW IKLELWTFTE YEKIVYLDTD
Cv   MARRGSTWPD SYLMGVQALA RSLLAAQAQH PLLVMYTVER YV-PAGHECW NKLRIWELEE YERLAYLDAD
Sd   RAYATLVCTD AYAIGAQVLR ASLHRVGSTL PLVVLVTYDV APIPLRSHAW AKLRVFELEM FDTIVFLDAD
Ac   EAFVTLLSSR SYYPGVVALA RSLRQFSA-R ELLVLTTVPV ERVPPPEDCF TKFRMFELKN YTKFVYLDAD
Ba   EAYVTHLTND QYIKGAQVLA ESLREAGATR PPLAMITVPE FGDGRKDGFF TKLEAWRLPC -TRVIYLDTD
Ab   FAYVTVHYDQ EYVLGIQVLM QSIKLSGTRH DLVVLVSVVD ITNPFLNHTL NKLHVWNLLE YDRVVYLDAD
Pi   FAYVTVHYDA EYVLGVQVMM HSIKLTGSPY DLVVLASVTN IDNPFVGYTL NKLHVWNMLE YERVVYLDAD
Ng   YAYATLVSSE GYLSGALAMY KSIIARGGKY DLVLVVTASY IDNPNAKDTY NKLHIWKLDQ YKRLVFVDSD
Vc   EAYATLVYGE DFVLAARVLG QSLRESGTTR DMVALTTVAP VKNPGTGYVY TKLYIFQMTE YKKIVFLDAD
Gs   YAYATLLCDD VMLPATRAWL QSLKMTNTSF PIVVLVLVTP LEYPFTLCRY SKLHLWNLLN YDKVVYMDSD
Cm   YAYATLLCDE RMLRAVAALV HSLRVRNTSY PILVLTTREP LPYPFALCRY AKLHLWSLTT YEKIVFLDGD
Rg   YAYITLVTNA DYAKGATALV RSLRLTKTAA NIVVLHTIAL APLADLGCNF CKLRLWQLTE YERIVFIDAD
```

```
          ....|....| ....|....| ....|....| ....|....| ....|....| ....|....| ....|....|
                  80         90        100        110        120        130        140
Hs        TLVLANIDDL FDREELSAAP DPGWPDCFNS GVFVYQPSVE TYNQLLHLAS EQGSFDGGDQ GILNTFFSSW
Mm        TLVLSNIDDL FEREELSAAP DPGWPDCFNS GVFVYQPSIE TYNQLLHLAS EQGSFDGGDQ GLLNTYFSGW
DM        TLVLQNCDEL FEREELSAAP DVSWPDCFNS GVFVFKPSVD TFAQITEFAV KNGSFDGGDQ GLLNQFFADW
Mb        TLVLTNIDEL FERNCFAAAP DIGWPDCFNS GVFVFQPSSA KFEDLVRLLA STGSFDGGDQ GLLNEYFADW
Hr        TLVLQNVDDL FSRDPFAAAP DAGWPDCFNS GIFLYQPSFE MYGDLLQFAL KIGSFDGGDQ GLLNLFFSDW
Ta        TLVIGNIDDL FTRPELSAAP DVGWPDCFNS GVFVYKPSMQ TYQTIVAFAL QFGSFDGGDQ GLLNEFFNTW
Nv        MLVLQNCDEL FDRCELSAVP DIGWPDCFNS GMFVFEPSRA THEALLKYAI DHGSFDGGDQ GLLNSFFSQW
Co        TLVVQNVDEL FDRPEIAAAP DVGWPDCFNS GVFVFVPSAA TFEKLAEHAV STGSFDGGDQ GLLNTFFDYW
Pm        AFCLKNIDEL FDLDTFAAVP DVGWPDIFNS GVFITKPNIS VYNSLLNLAK NSISFDGGDQ GLLNFYFSNW
Aq        TLVLTNVDEL FEREEMSAAP DIGWPDLFNS GVFVFRPSLE TFASLLELAD KEGSYDGGDQ GLLNLYWRDW
Sc        TLPLNKEFLL FDIMSVGAIA DIGWPDMFNS GVMMLIPDAD TASVLQNYIF ENTSIDGSDQ GILNQFFREW
Tg        CIVLRPVDEL FLRQPPAFAP DIFPPDKFNA GVAVLKPDLG EYGNMVAAVE RLPSYDGGDT GFLNAYFSSW
Hm        CIVLRPIDDL FLRQPPAFAP DIFPPDKFNA GVAVLKPDLD EYGKMVAAVE RLPSYDGGDT GFLNAYFSSW
Nc        CIVLGPVDEL FLRKPPAFAP DIFPPDKFNA GVVVLKPDLG EYGKMIAAIE RLPSYDGGDT GFLNAYFSSW
Sn        CVVLQNVDEL FERLSPAFAA DVFPPDRFNA GVIVLQPNVE LFSRMLRAAG LLPAADGGDT GFLNSFFSDW
Kb        CVVLERVDEL FERPSPAFCP DVFPPDKFNA GVIVLSPSRE LFEKMQERIA ELPSHDGGDT GFLNAFFPDW
Vb        CIVMRKIDCL FDPAAPAFAP DVFPPDRFNA GVMVIEPSLA VYEDLLAKRT VLRSYDRGDT GFLNAYFSGW
Tp        CLVVKDVSHL LRVDSLAAAP DIFPPDKFNA GVMVLCPSKA VFNDMMARLN SCTSYDGGDT GFLNSYYPNW
Rf        CIVVRDISDI FKLPDFAAAP DLCPPDHFNA GVLFIQPNVQ TFQQLLRNVA YVNSYDGGDT GFLNSYFNDW
Bn        AIVVRNVDHL FKMIPFAAAP DIFPPDRFNA GVLVLQPNSV MFAYILRLAY GLGSYDGGDT GFLNRIFPRW
Pu        AIVLQNVDEL FDRSTFAAAP DVFPPDRFNA GVLVIRPNKQ LFADLLAKAK ELKSYDGGDT GFLNAFFPKW
Ot        CLISSNPENA FDRNSFAAAP DVFPPDRFNA GVLLIKPSMT VFRDMISKIL TFPAYDGGDT GFLNAYYPDW
Sl        CLIMQNPENI FLRDTFAAAP DVFPPDKFNA GVLYIEPSMK IFTDLISKIQ ILSTYDGGDT GFLNAYFPNW
Ws        ALVMEDLDEL FDREVFAAAP DVFPPDKFNA GVMVVVPSLI VLEDMMSKVE ELPSYDGGDT GFLNAYFADW
Ae        AFVLANVDEV LERDIFAAAP DIFPPDRFNA GVLLLHPNAE LFQRLVSQSA QFQSYDGGDT GYLNAVFPDW
Al        ALILTNIDEL FEMDTFAAAP DIFPPDRFNA GVLVIKPGKD VFENLLAKAK TIKSYDGGDT GFLNLVFSDW
Ng        CLVVEDIQEL FSADVFAAAP DIFPPDRFNA GVMLVRPNLD VYEDMLRAVG ALPSYDGGDT GFLNAFFPKW
Gt        LLPLSSLAPL FDRDVVAAVP DISLPDHFNS ALVLLRPNLL HLQRLLALSS SLEPYDGGDQ GLLNEFFNAW
At        LLILRNIDFL FSMPEISATG NNGTL--FNS GVMVIEPCNC TFQLLMEHIN EIESYNGGDQ GYLNEVFTWW
Os        ILVLRDLDAL FGFPQLTAVG NDGSL--FNS GVMVIEPSQC TFQSLIRQRR TIRSYNGGDQ GFLNEVFVWW
Dd        MLLLKSLDHL FDLVDLYAAI DADANSCINS GIMLLSPSID VYNLLIDGMK LPNQSTVNDQ DVINTTLPHW
Tv        TLPTQRIDEL FNHSELSCVS DPMPPQICNT GLLVLEPNLT TFKHMKKLSD LYANNPPGDQ GFINFFFGQF
Cv        MLVLRNIDHL FALPPFYAAP DCTAGRQFNA GFFLVTPSRA ELARFQSLLV RIGGY--AEQ DLLNEVLHEF
Sd        MLCVRNMDDL FDAIAAASRA CTCNPQRFNS GMLVLHPSCA TLESLLAKLR SVERFVFSDQ CFLNEAFPDF
Ac        MLVVGDVDEL FSYPSFAAAP NFQLKKSFNA GLFVVDRDEG LHRQFLDHYH YDKAWSWADQ SLLNDFFKKW
Ba        ILAVGNPDVL FELAQFAVQD SQPHMQGPNT GVMVLKPDIR VYARIVETLT PLHEMPFYEQ GFIGKFFAKW
Ab        NIVLRNADEL FMCGPFCAVF MNPCH--FHT GLLVVTPDKE EYQRLLHQLE YQSSFDGADQ GFLSSVYSEL
Pi        NVLIRNSDEL FLCGEFCAVF MNPCH--FHT GLLVVTPSAA EYQRLLSALG HLESFDGADQ GFLSSMYSML
Ng        CIIFKNVDLL FNCVGVCSGS DMGNTEFFNG GIMVLEPSTK TYDDMMDKMP AYKSYDGGEQ GFINLYFDFH
Vc        VLVIRNMDVI FKCPGFCAAL RHSER--FNT GVMSLVPSLE MYDDMMAKMR SMPSYTGGDQ GFLNSYFPSF
Gs        MLVMQNIDNL FVEFDLSACA DLYPDT-FNS GIMVIQPNET TFRNMKAVYK NVSSYNVGDQ GFLNWFFGEW
Cm        TLVLAPIDDL FEKYDLAAAP DLYPET-FNS GVMVLEPRHD VYASMLARYR ETPSYNLGDQ GFLNSFFGQW
Rg        AIILKNIDKL FAYPEFSAAP NVYETRRMNS GVFVARPSEE TFGRMLAMLD QPDAFRRTDQ TFLEAFFPDW
```

```
      ....|....| ....|....| ....|....| ....|....| ....|....| ....|.
              150        160        170        180        190
Hs    ATTHLPFIYN LYSYLPAFKV FGASA----- -KVVHFLGRV KPWNYTHPEF LILWWN
Mm    ATTHLPFVYN LYSYLPAFKA FGKNA----- -KVVHFLGRT KPWNYTHPEF LNLWWD
DM    STAHLPFVYN VYCYLPAFKQ FRDKI----- -KILHFAGKL KPWLIQAQDL IQLWWN
Mb    ATQRLPFAYN MYGYAPAFER FKADI----- -KVIHFIGAR KPWMGM---- ------
Hr    ATKHLPFTYN LYSYKPALKK FGDEI----- -KIVHYLGKP KPWDHENMEL LQLWWD
Ta    ATSHLPFTYN MYWYAPALNR FSKDI----- -KVVHFIGAL KPWHHLLTNY VQRWWE
Nv    SHEHLSFIYN MYTYAPAYKE FGKNV----- -KIVHFIGPV KPWQYSERSY IQLWWD
Co    PTARLSFLYN MYSYKPAFQK YGHLV----- -KIIHFIGQF KPWHWASEFH VQQWWN
Pm    K--RLPFTYN VYQYFPAYYH FKSKI----- -SVIHFAGTK KPWMLSYNEL IEKWKS
Aq    SIRRLPFTYN VYSYPPAFLR HRKDM----- -KIIHFLGAI KPWHHRAEEF IRKWWE
Sc    V--QLSFTYN VYQSSPAMNY FKPSI----- -KLIHFIGKH KPWSLWKNEY HDQWNE
Tg    YENRLPFRYN ALRTLYHMTY SSRKGYWDAV IKILHFCSSP KPWEQPKTDL EELWWK
Hm    YENRLPFRYN ALRTLYHMTY SSRKGYWNAV IKILHFCSSP KPWEQPKTDL EELWWK
Nc    YESRLPFRYN ALRTLYHMTY CSHKGYWNAV IKILHFCSSP KPWEQPKTDL EDLWWK
Sn    YMWRLPFKYN AQRSVYRFTG AAYRGYWEAI IKILHFTSTP KPWERPQTEL EDIWWS
Kb    YRWRLPFRYN ALRTMYWFTH KN-PGYWDSL IKILHFCSSP KPWDPEKGDL EQLWWE
Vb    YGWRLAFHYN AQRTMHWMTH SKQPGYWDEC LSVLHLSSSP KPWESPKGPT EWLWWN
Tp    FGGRLSFGYN AQRFMHHCTY EKQPKYWDDG VYIVHFSSSP KPWETKHGTL ESKWQL
Rf    YHGRLDFGWN AQRIMEWYTR DK-PAYWDHI VRILHFSSSP KVWDIPSNRL HRQWHS
Bn    HSWRLHFGYN AQRTLHWFTK -KNPKYWEWS LHIIHYASSP KPWEVPTDKL EKIWWK
Pu    FESRLPFGYN AQRTMYWLVN GKNPGYWNAV LKILHYSSNP KPWEDPKGDL EILWWQ
Ot    YLKRLPYGYN AQRTLYWFTI KRTDGYWKEI LVIIHYSSSP KPWVG-KGDL ELLWFQ
Sl    FESRLPYGYN AQRTLYWFTI KRTDGYWKEV IIIIHYSSSP KPWSSQKGDL ELEWFK
Ws    FSRRLPFAYN ALRTVYWTTH EKNPGYWEAI VKIIHFCSSP KPWEETKGDL EMTWWQ
Ae    YTYRLPFAYN AQRTMHWLTY AKKPGYWDAV VKVLHCSSSP KPWESPKGDL EMLWWQ
Al    FQRRLPFRYN AQRTMYWMVN SKNPGYWKAV LKILHFSSSP KPWEEPIGDL EMIWWM
Ng    YSSRLPFIWN AQRTLHWMTH AVAPGYWGAV VKILHFSSSP KPWEPEKGEL EVKWWT
Gt    YESRLGLELN LSRLHPRSWL RTLPRQRSNL SQVIHFSGGR RPWGIASVAA AALVWH
At    HRILKHFWIG DRKKTELFGA EPPVL----- -YVLHYLG-M KPWLCYTDIA HRKWWM
Os    HRLLKNFWAN TRALKERLFR ADPAE----- -WSIHYLG-L KPWTCYSDAA HARWWQ
Dd    RSLEYGVQIT HCTSEPRLWN F--------- -TFLHFTAGP KPWSLLPTCI EQIYLN
Tv    N--PLPTLYN VDTNFEFLYE QKLI------ -KVVHFVC-K KPWKCGMYSL NQVWWD
Cv    SAPPLPHTFN ARRHHPQLWR ---------- -QHWHAVAVA KPWQEGYQDL VQLWWR
Sd    I--DVPYVFN APIAHPRLWQ LEDV------ -KAIHYIL-E KPWHVEYDDL YALWWE
Ac    N--QVPHYFN MFLYRPDLWE VDKI------ -KIIHYTG-G KPWQTPPYEP LFALWR
Ba    V--QLPAKYN FYLNRPLYQD IRHDN----- KVFIHYAK-C KPWDLSFGKE YLRYIR
Ab    RKARLSVGYN IYEQYHWKLF YLRHFATMTS RPIPAITIGL KPW------- --YWWA
Pi    RKARLPVGYN IYEQYHWKLF YLRQFASMTS RPIPALTVGL KPW------- --YWWA
Ng    RKSRIPYTWN TYYFFKYAYI QRLKK----- FRIIHYNLPI KPWKFLILDA SYYWYE
Vc    AHSRLPTTFN ALYVVGSNRW MLPRS----- LYVIHYTLGF KPWVWWRENP AWQAYR
Gs    SQRHIPLKYN VLKYRDTIMW GHVKD----- IKVLHFTGET KPWNFYEMRS YYAWVR
Cm    RANHLPLEYN TLKLRETILW ASLQR----- VRVVHFTGET KPWSWHDRHI DPVFYY
Rg    HG--LPVYFN MLQYVWFTMP AL---WDWKS ISVLHYQYE- KPWEKDHPKL IDLWHS
```

**Species names. Sequence IDs are in Fig. S7.**

**Glycogenin-like:**
*Hs: Homo sapiens*
*Mm: Mus musculus*
*DM: Drosophila melanogaster*
*Mb: Monosiga brevicollis*
*Hr: Helobdella robusta*
*Ta: Trichoplax adherens*
*Nv: Nematostella vectensis*
*Co: Capsaspora owczarzaki*
*Pm: Pneumoycystis murina*
*Aq: Amphimedon queenslandica*
*Sc: Saccharomyces cerevisiae*
**Gat1-like**
*Tg: Toxoplasma gondii*
*Hm: Hammondia hammondi*
*Nc: Neospora canimum*
*Sn: Sarcocystis neurona*
*Kb: Karenia brevis*
*Vb: Vitrella brassicaforma*
*Tp: Thalassiosira pseudonana*
*Rf: ReticulomYxa folisa*
*Bn: Bigelowiela natans*
*Pu: Pythium ultimum*
*Ot: Oxytricha trifallax*
*Sl: StylonYchia lemnae*
*Ws: Ectocarpus siliculosus*
*Ae: Aphanomyces euteiches*
*Al: Albugo laibachii*
*Ng: Nannochloropsis gaditana*
**Other CAZy GT8 family**
*Gt: Guillardia theta*
*At: Arabidopsis thaliana*
*Os: Oryza sativa*
*Dd: Dictyostelium discoideum*
*Tv: Trichomonas vaginalis*
*Cv: Chlorella variabilis*
*Sd: Saprolegnia diclina*
*Ac: Acanthamoeba castellanii*
*Ba: Bigelowiella natans GT8*
*Ab: Albugo laibuchi GT8*
*Pi: Phytophthora infestans*
*Ng: Naegleria gruberi*
*Vc: Volvox carteri*
*Gs: Galdieria sulphuraria*
*Cm: Cyanidioschyzon merolae*
*Rg: Rhizobium gallicum*

**Fig. S9.** Characterization of Gat1 enzyme activity and biochemical complementation of *T. gondii* extracts. TgGat1 glycosyltransferase activity was assayed using 20 mM maltose-pNP as an acceptor in the presence of 4 µM UDP-Glc, 2 mM $MnCl_2$, pH 7.0, and varying concentrations of NaCl or KCl (**A, B**). (**C**) TgGat1 was assayed using 20 mM maltose-pNP in the presence of 5.2 µM UDP-Gal, no added salt, 2 mM $MnCl_2$, pH 7.0, and the indicated divalent metal ions. (**D**) TgGat1 was assayed using 20 mM maltose-pNP, 8 µM UDP-Gal, no added salt, 2 mM $MnCl_2$, at different pH values. **E.** Donor specificity of TgGat1, based on the UDP-Glo assay in an overnight reaction that consumed all UDP-Gal. **F.** UDP-Gal concentration dependence of TgGat1 and PuGat1 Gal-transferase activity toward 20 mM maltose-pNP. **G.** Concentration dependence of TgGat1 Gal-transferase activity on GlFGaGn-Skp1 concentration. Error bars represent ±S.D. of 3 technical replicates of the same reaction. See Fig. 4 for related data.

### Figure S9. αGalT activity studies

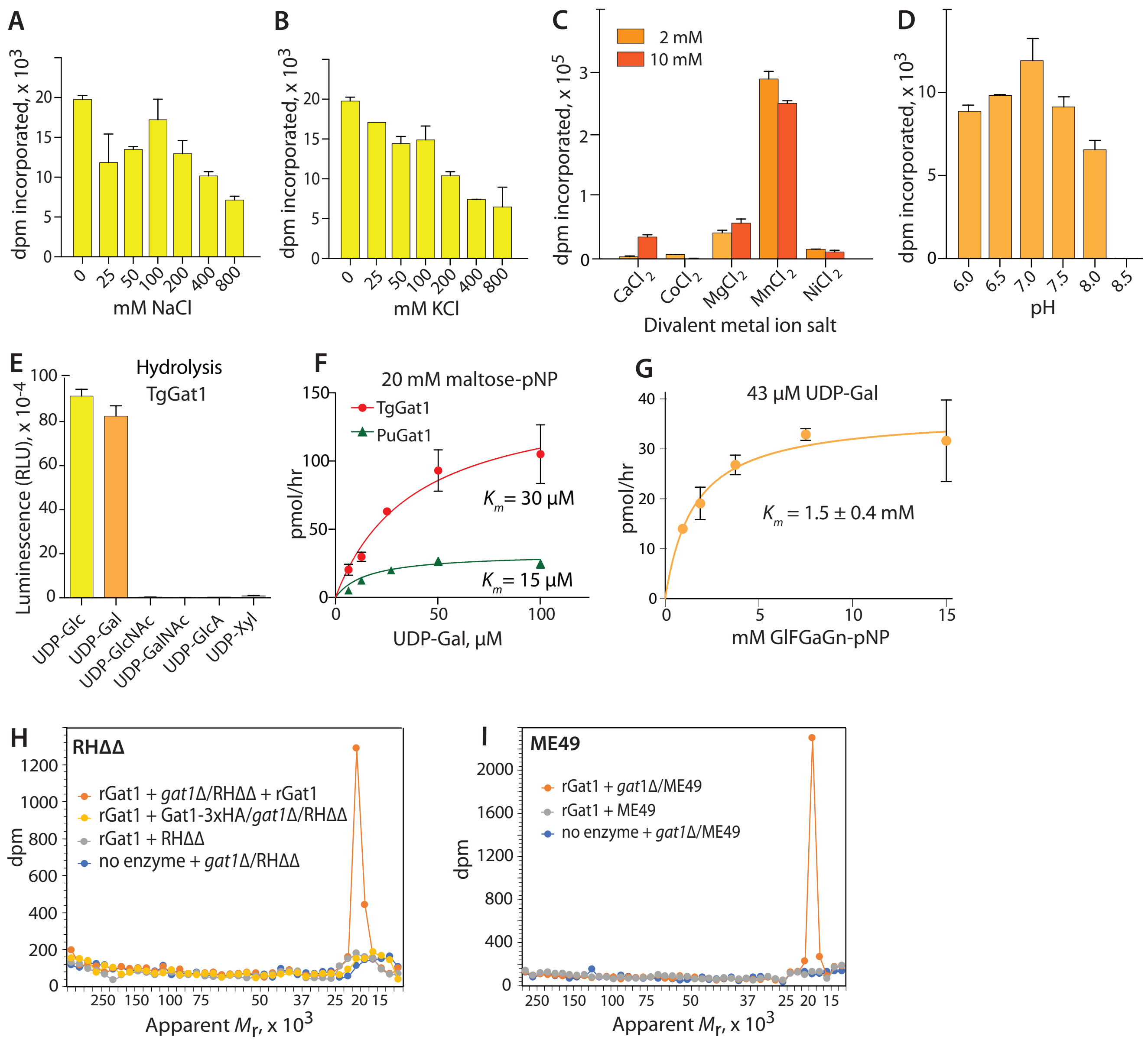

**Fig. S10.** Test for TgGat1 auto-glycosylation activity. PuGat1 and TgGat1 were prepared in *E. coli*, purified to near homogeneity (Fig. 4A), and analyzed by nLC/MS analysis. **(A)** Total ion current for elution of PuGat1 in a gradient of acetonitrile from a C4 column. **(B)** Mass spectrum showing multiply protonated species. Xtract deconvolution yielded virtually only one species with an $M_r$ 30251.2596, which closely matched the predicted theoretical monoisotopic mass of $M_r$ 30251.2603 (error= 0.02 ppm). **(C)** Deconvolution of data in panel B using the ReSpect algorithm in BioPharma to yield a measurement of the average mass. **(D)** SDS-PAGE and Coomassie blue staining of TgGat1, before and after incubation with UDP-Glc or UDP-Gal for 30 min. **(E)** Summary of average mass measurements of TgGat1 and PuGat1 based on ReSpect deconvolution. After isolation from *E. coli*, both TgGat1 and PuGat1 yielded predominantly only the unmodified versions of the recombinant proteins, with $M_r$ 39051.9687 for TgGat1 (theoretical average mass: 39051.9161, error= 1.3 ppm) and $M_r$ 30269.1738 for PuGat1 (theoretical average mass: 30269.2901, error= 3.8 ppm). Their masses were essentially unaffected by *ex vivo* reaction in the presence of UDP-Gal or UDP-Glc.

#### Figure S10. Autoglycosylation test

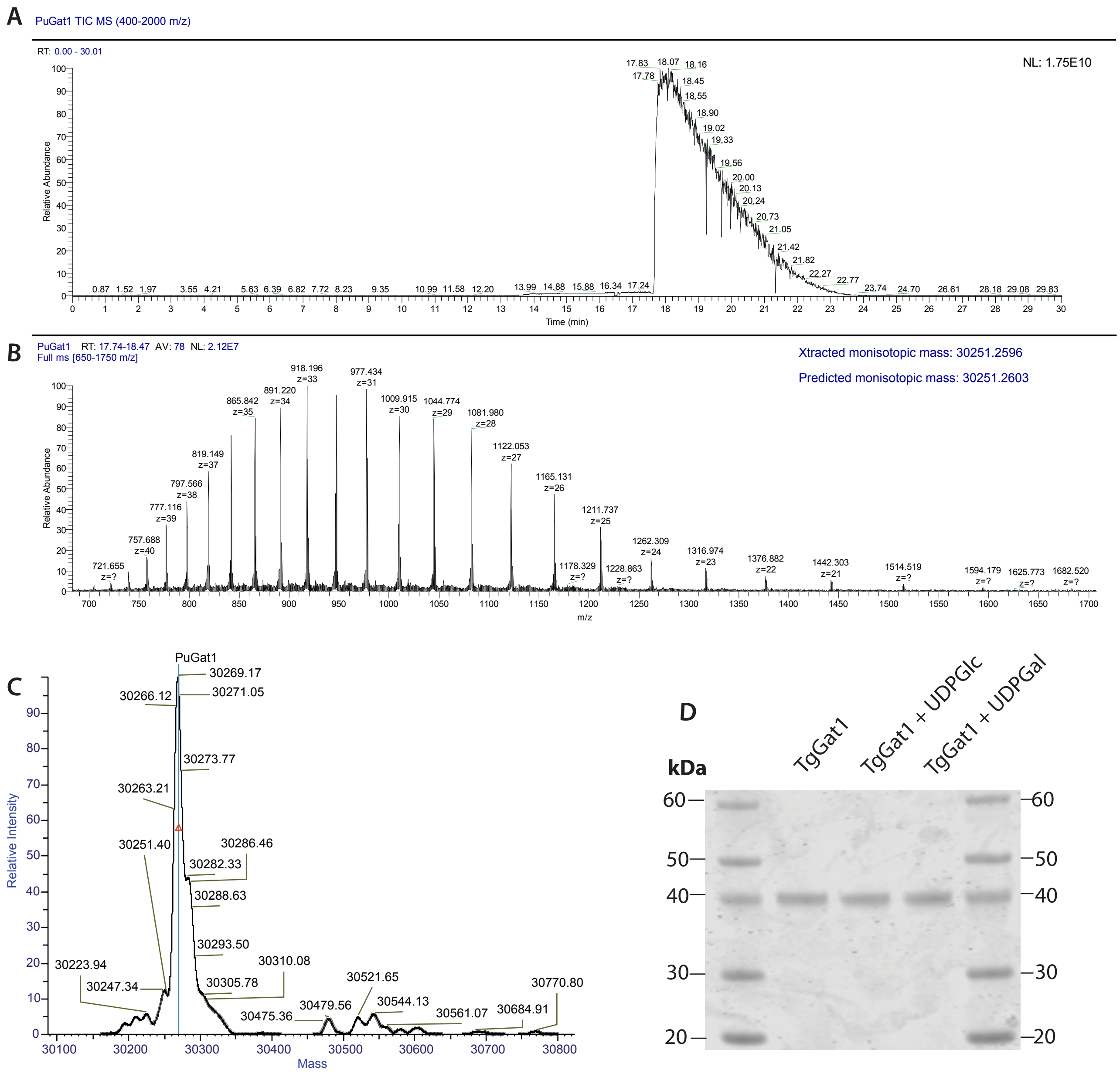


E

| Sample Name | Protein Name | Average Mass | Theoretical Mass (Da) | ΔMass (ppm) | Intensity | Fractional Abundance | Score | Number of Charge States | Charge State Distribution | Mass Std Dev | PPM Std Dev |
|---|---|---|---|---|---|---|---|---|---|---|---|
| TgGat1 | TgGat1 | 39051.9688 | 39051.9161 | 1.3 | 1.51E+08 | 5.44 | 136.39 | 34 | 21 - 54 | 1.68 | 42.93 |
| TgGat1+Glc | TgGat1 | 39051.3359 | 39051.9161 | 14.9 | 4.73E+07 | 2.25 | 103.70 | 28 | 26 - 53 | 1.92 | 49.12 |
| TgGat1+Gal | TgGat1 | 39052.7539 | 39051.9161 | 21.5 | 7.38E+07 | 4.73 | 83.04 | 31 | 22 - 52 | 2.39 | 61.19 |
| PuGat1 | PuGat1 | 30269.1738 | 30269.2901 | 3.8 | 1.86E+09 | 20.86 | 92.16 | 26 | 17 - 42 | 0.98 | 32.34 |

!

**Fig. S11.** NMR Analysis of the TgSkp1 pentasaccharide. ($^{13}C_6$)GlFGaGn-pNP (Glc is U-$^{13}$C-labeled) was partially (20%) modified by TgGat1 or PuGat1 in the presence of UDP-[U-$^{13}$C]Gal. **(A)** 1D 900 MHz $^{1}$H-NMR spectrum of. Magnification shows the region of $^{13}$C-anomeric carbons in the mixture of modified and unmodified tetrasaccharide. A cartoon diagram of the TgSkp1 pentasaccharide attached to pNP (Galα1,3Glcα1,3Fucα1,2Gal-β1,3GlcNAcα1-pNP) is shown at the top using glycan symbols from Varki et al. (2015). **(B)** $^{1}$H –$^{13}$C-HSQC, -HMBC, and –HSQC TOCSY spectra demonstrating anomeric carbon to ring proton correlations. The Gal-H1/C1 doublet peaks were too weak to observe and are indicated by boxes. **(C)** $^{1}$H –$^{1}$H –COSY and $^{1}$H –$^{13}$C-HMBC spectra. Identical results were obtained using ($^{13}C_6$)GlFGaGn-pNP modified with PuGat1 (not shown).

### Figure S11

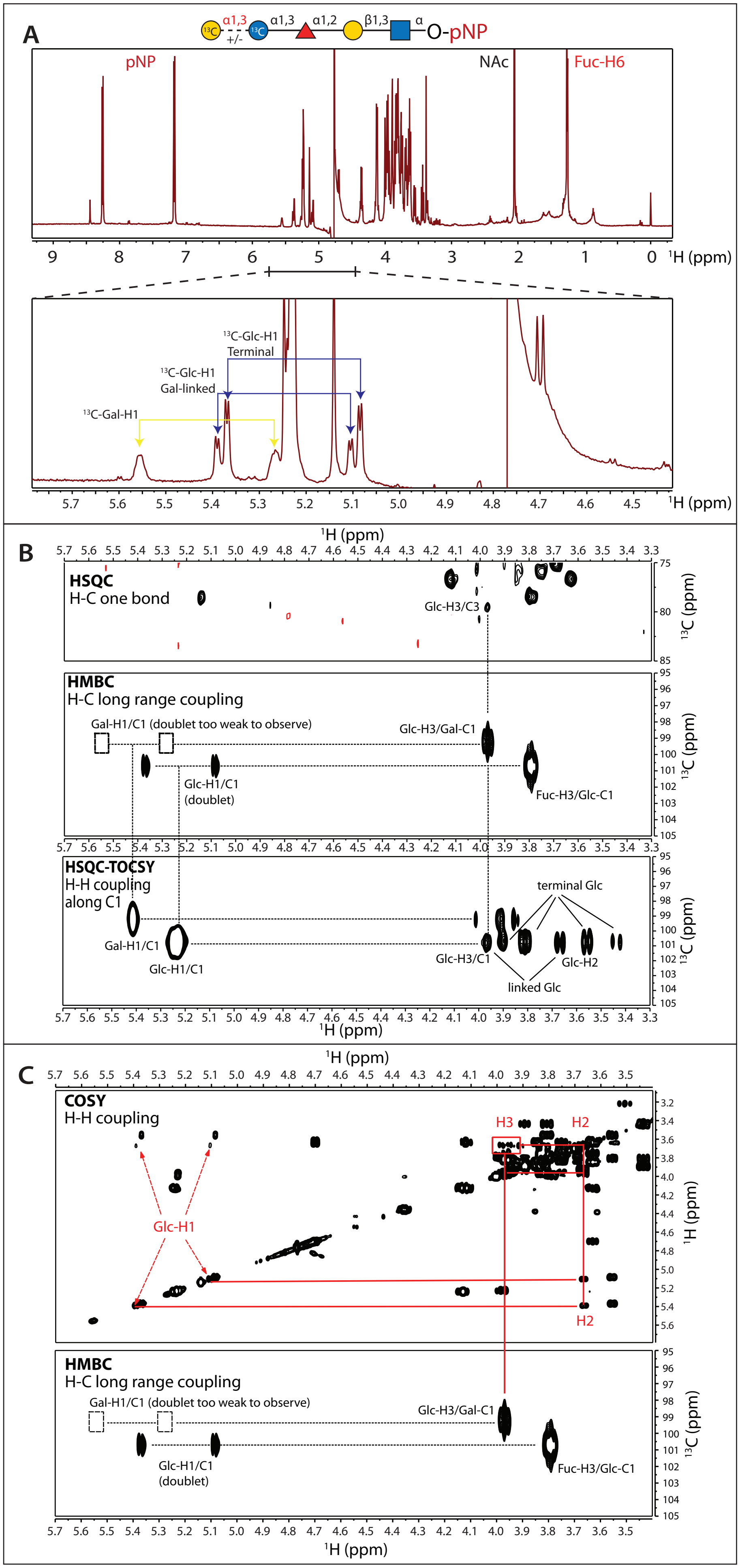

A
α1,3
+/-
α1,3
α1,2
β1,3
α
O-pNP
pNP
NAc
Fuc-H6
¹H (ppm)
¹³C-Glc-H1
Terminal
¹³C-Glc-H1
Gal-linked
¹³C-Gal-H1
B
HSQC
H-C one bond
Glc-H3/C3
HMBC
H-C long range coupling
Gal-H1/C1 (doublet too weak to observe)
Glc-H3/Gal-C1
Glc-H1/C1
(doublet)
Fuc-H3/Glc-C1
HSQC-TOCSY
H-H coupling
along C1
terminal Glc
Gal-H1/C1
Glc-H1/C1
Glc-H3/C1
Glc-H2
linked Glc
¹³C (ppm)
C
COSY
H-H coupling
H3
H2
Glc-H1
H2
HMBC
H-C long range coupling
Gal-H1/C1 (doublet too weak to observe)
Glc-H3/Gal-C1
Glc-H1/C1
(doublet)
Fuc-H3/Glc-C1

**Fig. S12.** Computational comparison of the Skp1 glycans from *Toxoplasma* and *Dictyostelium*. **(A)** Superimposition of the two energy-minimized glycan structures produced by the Glycam webserver (Woods, 2014). Residues are colored according to the SNFG system. The differing Glc (blue) and Gal (yellow) residues (arrowhead) mark the difference between *Toxoplasma* and *Dictyostelium* glycans, respectively. **(B)** Superposition of the glycans in the context of Skp1 (orange ribbon); note that the linkage to Hyp is not shown. **(C)** Illustration of hydrogen bonds present at >25% occupancy over all simulations (1.5 µs) in *Toxoplasma* Skp1. **(D)** Comparison of amino acid sequences of TgSkp1 and DdSkp1 over the region depicted. Red asterisks indicate residues involved in hydrogen bonds that correlate best with extension of helix-8 (see panel E), green asterisks indicated residues that contribute most to non-polar packing interactions (see Table 2), and the black asterisk indicates the attachment site after hydroxylation. Residues are labeled from below according to the hydrogen bond with which they are associated. **(E)** Summary of the six 250-ns trajectories (3 pre-equilibrated; 3 were not). Left bars of each pair summarize the average distances for each trajectory between C156, near the C-terminus, to the center of mass of residues 1-136 (dashed green line in Fig. 6A), scaled to the highest average distance (Equil-3, in which the average value was >50 Å in the observed range of 18-61 Å for at least 85% of the time sampled at 0.1 ns increments). Right bars summarize the 5 most frequent hydrogen bonds between the glycan and Skp1, normalized to the highest level of hydrogen bonds observed in a single trajectory (Equil-1, in which at least one of the hydrogen bonds was occupied >99.7% of the time sampled in 0.1 ns increments). At the right is shown a time-resolved analysis of the correlation of helix-8 extension with the occupancy of each hydrogen bond over the entire 1.5 µs of simulation time, based on the Pearson's correlation coefficient (linear regression $R^2$).

#### Figure S12

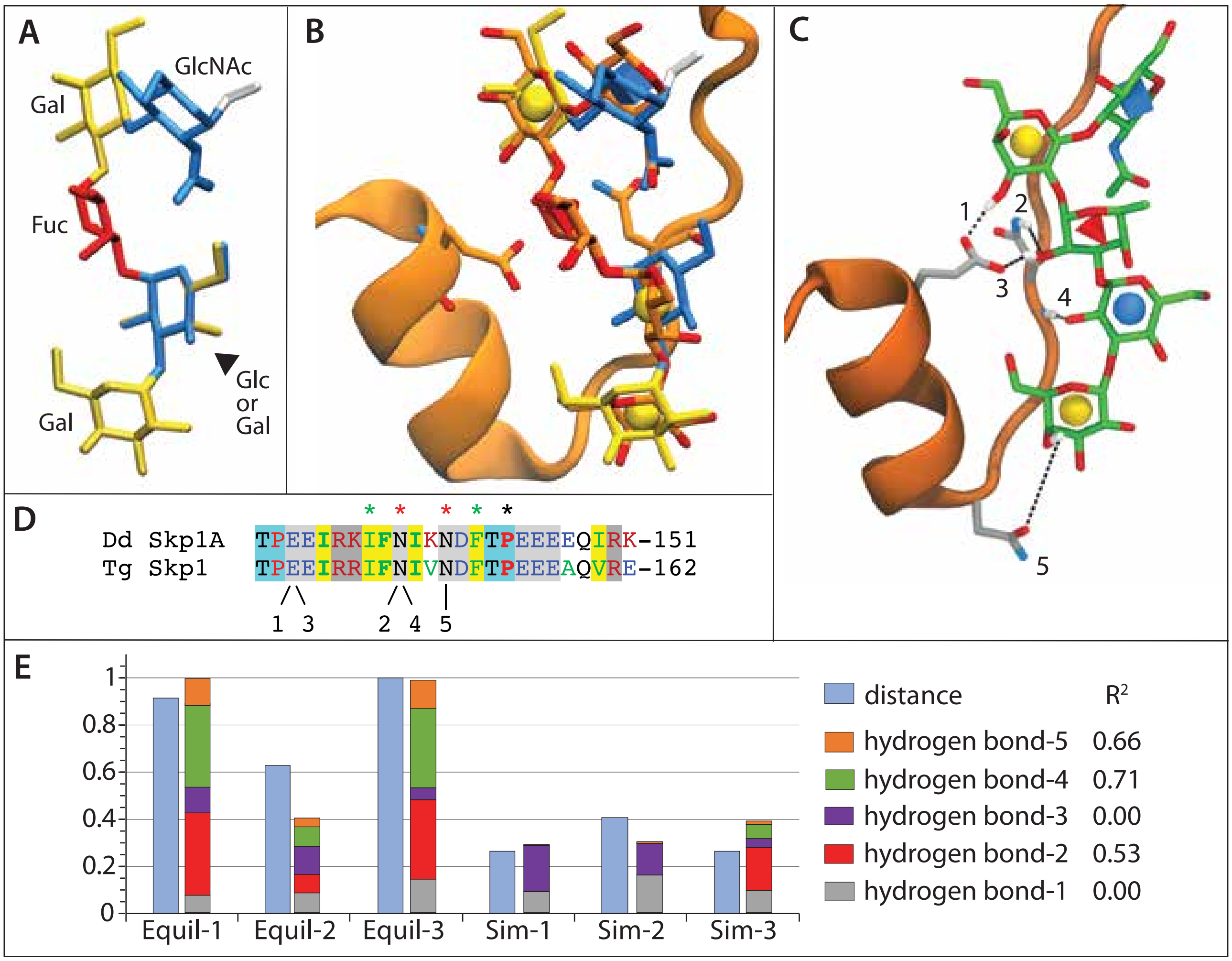
A
GlcNAc
Gal
Fuc
Glc
or
Gal
Gal
B
C
1
2
3
4
5
D
Dd Skp1A TPEEIRKIFNIKNDFTPEEEEQIRK-151
Tg Skp1 TPEEIRRIFNIVNDFTPEEEAQVRE-162
1 3
2 4 5
E
1
0.8
0.6
0.4
0.2
0
Equil-1
Equil-2
Equil-3
Sim-1
Sim-2
Sim-3
distance
R2
hydrogen bond-5 0.66
hydrogen bond-4 0.71
hydrogen bond-3 0.00
hydrogen bond-2 0.53
hydrogen bond-1 0.00

**Fig. S13.** PuGat1 and Oc-glycogenin-1 ligand interactions. PuGat1:UDP:$Mn^{2+}$ **(A)** and Oc-glycogenin-1:UDP **(B)**, from PDB 1LL2, are displayed as Ligplots (Laskowski and Swindells, 2011). Green dotted lines represent the interactions between the protein and the ligand, and red arcs represent packing interactions.

**Figure S13**

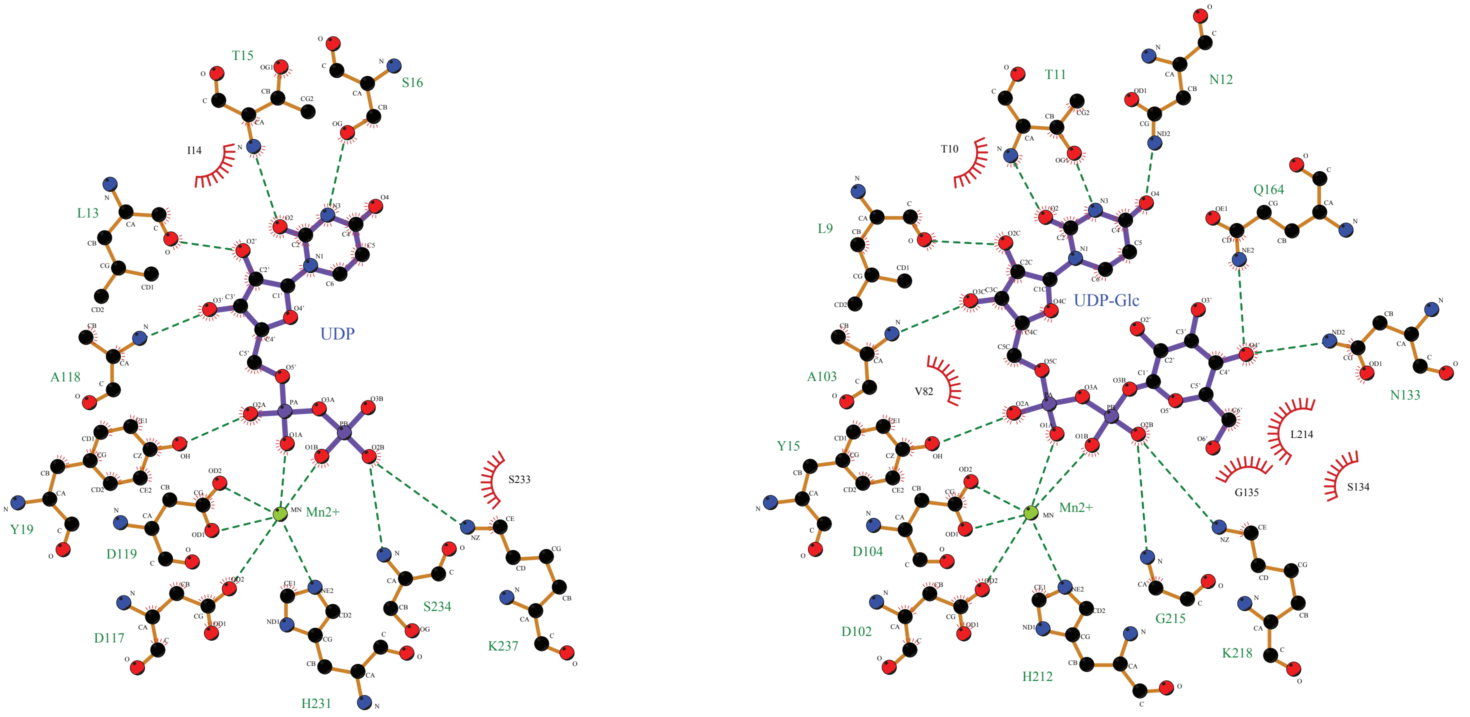


**A.** PuGat1:UDP:$Mn^{2+}$ **B.** Oc-glycogenin-1:UDP-Glc:$Mn^{2+}$

**Fig. S14.** Sedimentation velocity analyses of PuGat1. **(A)** Sedimentation profiles of different concentrations of PuGat1 are displayed with fit data and residuals. 11 μM, 6.5 μM, and 3.5 μM concentrations were detected at 280 nm, 1.3 μM data were collected at 230 nm, and 0.65 μM and 0.3 μM data were collected at 220 nm. **(B)** Data modeled as continuous $c(s)$ distributions are shown (normalized to a value of 1 for the tallest peak). Black dashed line represents the determined S-value, and the red and green dashed lines respectively represent the predicted monomer and dimer S-values. The peak appearing at a near-zero S-value at the lower concentrations may be due to a buffer mismatch that became apparent at lower wavelengths.

#### Figure S14

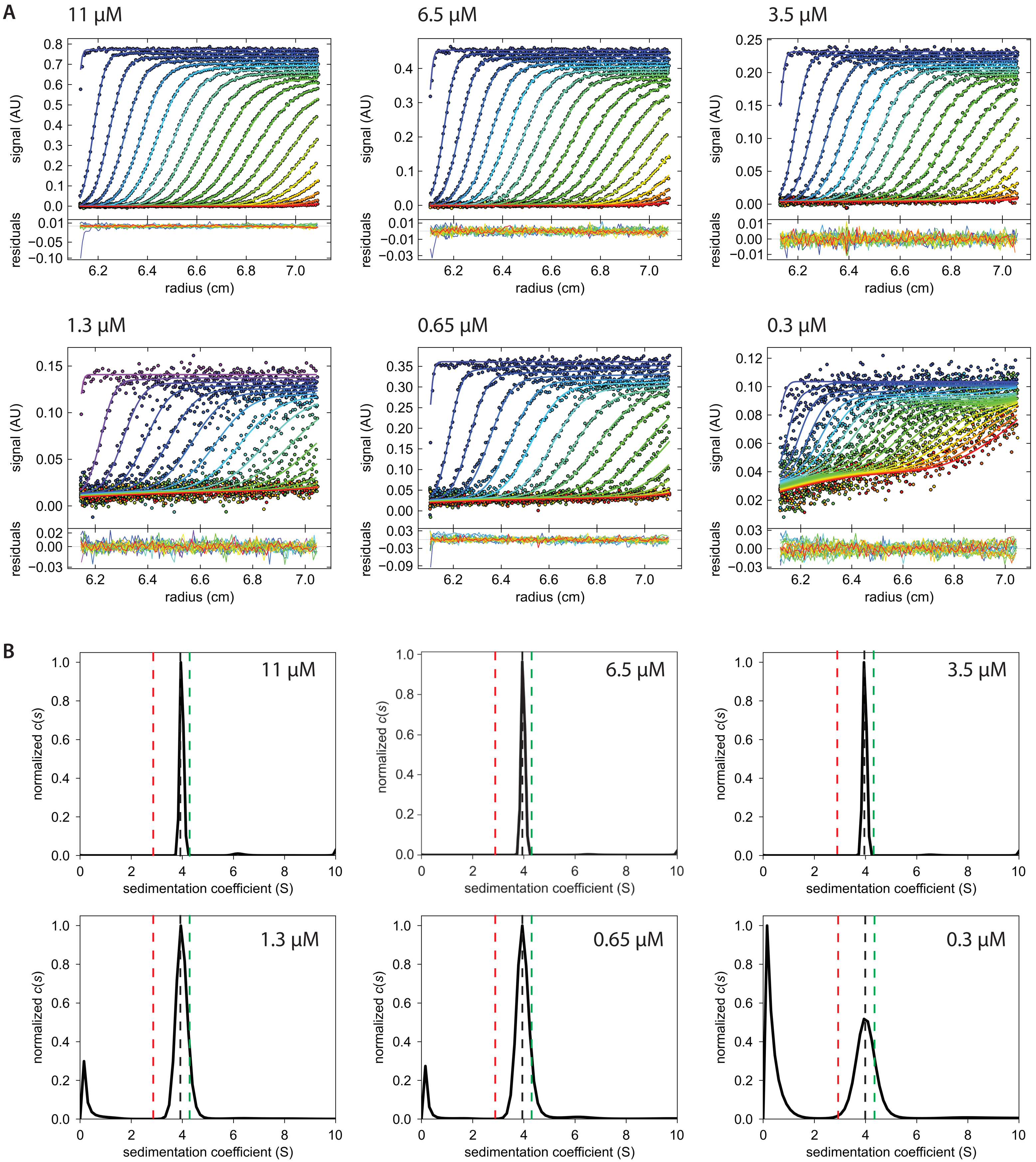
A
11 µM
6.5 µM
3.5 µM
1.3 µM
0.65 µM
0.3 µM
signal (AU)
residuals
radius (cm)
B
11 µM
6.5 µM
3.5 µM
1.3 µM
0.65 µM
0.3 µM
normalized c(s)
sedimentation coefficient (S)
